# Population-specific and transethnic genome-wide analyses reveal distinct and shared genetic risks of coronary artery disease

**DOI:** 10.1101/827550

**Authors:** Satoshi Koyama, Kaoru Ito, Chikashi Terao, Masato Akiyama, Momoko Horikoshi, Yukihide Momozawa, Hiroshi Matsunaga, Hirotaka Ieki, Kouichi Ozaki, Yoshihiro Onouchi, Atsushi Takahashi, Seitaro Nomura, Hiroyuki Morita, Hiroshi Akazawa, Changhoon Kim, Jeong-sun Seo, Koichiro Higasa, Motoki Iwasaki, Taiki Yamaji, Norie Sawada, Shoichiro Tsugane, Teruhide Koyama, Hiroaki Ikezaki, Naoyuki Takashima, Keitaro Tanaka, Kokichi Arisawa, Kiyonori Kuriki, Mariko Naito, Kenji Wakai, Shinichiro Suna, Yasuhiko Sakata, Hiroshi Sato, Masatsugu Hori, Yasushi Sakata, Koichi Matsuda, Yoshinori Murakami, Hiroyuki Aburatani, Michiaki Kubo, Fumihiko Matsuda, Yoichiro Kamatani, Issei Komuro

**Author notes:** Correspondence should be addressed to K.I, Y.K, and I.K.

## Abstract

To elucidate the genetics of coronary artery disease (CAD) in the Japanese population, we conducted a large-scale genome-wide association study (GWAS) of 168,228 Japanese (25,892 cases and 142,336 controls) with genotype imputation using a newly developed reference panel of Japanese haplotypes including 1,782 CAD cases and 3,148 controls. We detected 9 novel disease-susceptibility loci and Japanese-specific rare variants contributing to disease severity and increased cardiovascular mortality. We then conducted a transethnic meta-analysis and discovered 37 additional novel loci. Using the result of the meta-analysis, we derived a polygenic risk score (PRS) for CAD, which outperformed those derived from either Japanese or European GWAS. The PRS prioritized risk factors among various clinical parameters and segregated individuals with increased risk of long-term cardiovascular mortality. Our data improves clinical characterization of CAD genetics and suggests the utility of transethnic meta-analysis for PRS derivation in non-European populations.

---

Coronary artery disease (CAD) is a leading cause of global morbidity and mortality^1^. CAD is a common and complex disease, which is also known to be highly heritable^2^. To identify genetic factors underlying CAD, continuous efforts have been undertaken^3–9^, which have resulted in the discovery of over 160 CAD susceptibility loci.

These achievements have been mainly led by microarray-based genotyping technology in combination with haplotype imputation by publically available reference panels derived from large-scale sequencing projects. Recent advances in high-throughput sequencing technology have now made it possible to reveal the genomes of specific populations, including a substantial number of subjects with relevant diseases. These efforts have successfully discovered population-specific and disease-specific rare haplotypes^10–12^ and it is now possible to assess the pathological roles of these variants at the population level.^13^ Another critical achievement in complex-disease genetics is the advancement of the polygenic risk score (PRS). As recently reported, accumulating genetic knowledge and data enables us to derive a PRS which shows clinically relevant performance for case-control discrimination^14–17^.

Clinical implementation of these technologies is considered promising for the future of precision medicine in CAD. However, these data are mainly derived from large-scale GWAS predominantly conducted with subjects of European descent. Therefore, before these findings can be widely implemented in clinical settings, it is essential to confirm their relevance and robustness in diverse ethnic groups. Such transethnic analyses could further be used to explore novel findings that are otherwise difficult to discover in a single population^18–20^.

In the present study, to elucidate the genetic architecture of CAD and its transethnic heterogeneity, we conducted a large-scale GWAS of CAD in the Japanese population (25,892 cases and 142,336 controls) in combination with whole-genome sequencing of 4,930 Japanese individuals including rare disease haplotypes. This was followed by a transethnic meta-analysis (total of 121,234 cases and 527,824 controls). Additionally, using the PRS derived from these results and biobank-based phenome-wide analyses, we provide insights into the clinical utility of the PRS which may pave the way to the precision medicine.

## Results

### Construction of a CAD reference panel and imputation

We sequenced the genomes of 1,782 individuals with early-onset CAD and 3,148 non-CAD controls of Japanese ancestry from the Biobank Japan (BBJ) project and the Nagahama study (Supplementary Fig. 1a). After stringent quality control (Methods section), we established 48,305,902 variants [45,505,776 single nucleotide polymorphisms (SNPs) and 2,800,126 indels]. We found several pathogenic or likely pathogenic variants of familial hypercholesterolaemia (FH)^21,22^, which is the most important underlying genetic condition for CAD (Supplementary Table 1). To leverage the enriched disease-relevant haplotype information, we constructed a reference panel for haplotype imputation from the whole-genome sequence data (referred as BBJ_CAD_). For comparison, we constructed two reference panels from the 1000 Genomes Project (1KG)^23^ dataset under the same pipeline [1KG all subjects (1KG_ALL_) and 1KG East Asian subjects (1KG_EAS_)]. To assess the performances of these three different imputation panels, we imputed 179,320 Japanese genotypes with these three reference panels. After filtering by imputation quality (R^2^ ≥ 0.3), almost double the number of variants remained in the new panel (1KG_ALL_: 10,284,722; 1KG_EAS_: 8,482,456; BBJ_CAD_: 22,136,930, Supplementary Table 2). This difference was based on the improved imputation quality, especially when considering rare [1% ≤ minor allele frequency (MAF) < 5%] to very rare (MAF < 1%) variants (Supplementary Fig. 2, 3). We revealed a substantially increased number of variants with R^2^ ≥ 0.3 in the mechanistically annotated constrained class or curated pathogenic variants (Supplementary Fig. 4, Supplementary Table 3). For example, the BBJ_CAD_ panel imputed a 6 × higher number of stop-gain variants and a 5 × higher number of “Pathogenic” variants with R^2^ ≥ 0.3 compared to the 1KG reference panels.

### Nine novel loci for CAD detected in Japanese

Using the densely imputed genotypes with BBJ_CAD_ reference panel, we performed a GWAS on the case-control dataset from the BBJ, including 25,892 CAD cases and 142,336 controls (Supplementary Fig.1a), testing 19,707,525 variants. Although the lambda GC value was inflated at 1.24, the linkage disequilibrium (LD) score regression intercept^24^ was 1.06, indicating that 84% of the inflation was likely due to the polygenic nature of CAD. The liability scale heritability was estimated at 6.5%, which is slightly decreased compared to the results of a recent European study. Forty-eight loci reached genome-wide significance (Methods section); of them, nine loci were previously unreported (Table 1, Supplementary Table 4, Supplementary Fig. 5, Supplementary Data Set 1,2). A novel locus on 9q31 habours *ABCA1*, a critical gene for high-density lipoprotein cholesterol (HDLC) homeostasis, whose disruption causes a severe deficiency in serum HDLC^25^. The lead variant rs35093463 is located in the intron of *ABCA1*. The higher frequency of this variant in Japanese (MAF 36%) compared to Europeans (MAF 5%) might have contributed to the discovery of this loci in this study. Besides, we detected a relatively strong signal [MAF = 1.0%, odds ratio for CAD development (OR_CAD_) =1.61 and its 95% confidence interval (95%CI) = 1.46 – 1.78, *P* = 2.3 × 10^−21^] on the chromosome 17q25, where the lead variant rs112735431 causes a nonsynonymous substitution in the *RNF213* gene. rs112735431 is also known as an established susceptibility variant for moyamoya disease^26^, which is a rare cerebrovascular disease with abnormal vascular formation or obstruction. Since this variant is highly specific to the East Asian population, previous European studies could not assess its importance in CAD.

**Table 1.**
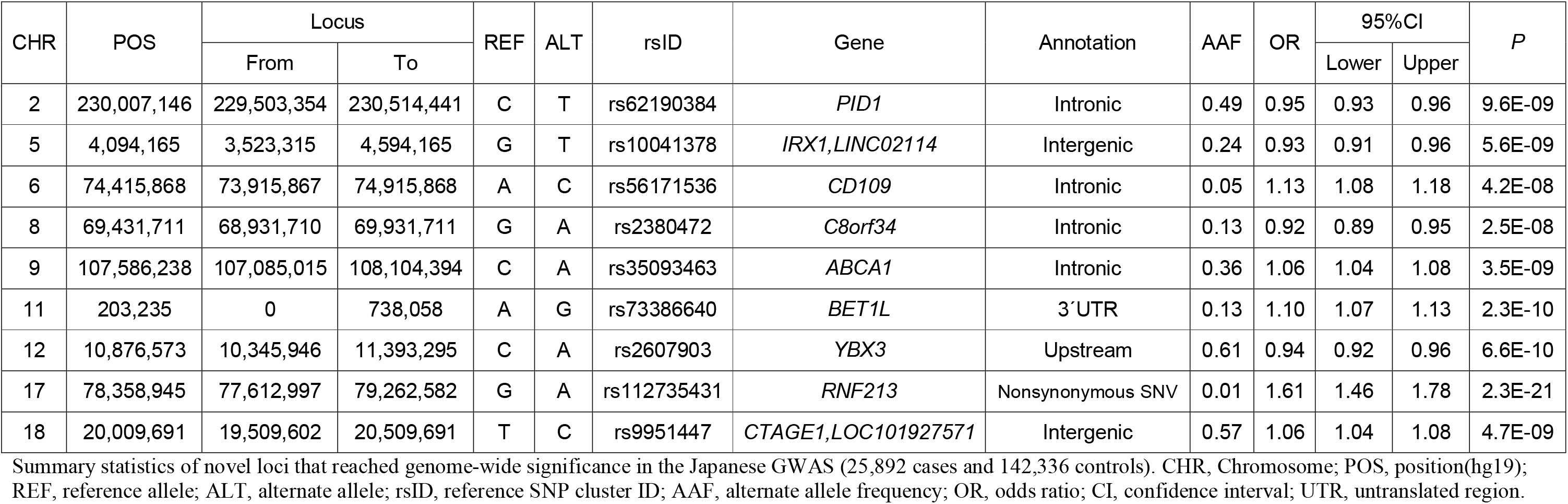
Novel loci identified by Japanese GWAS.

| CHR | POS | Locus |  | REF | ALT | rsID | Gene | Annotation | AAF | OR | 95%CI |  | *P* |
| --- | --- | --- | --- | --- | --- | --- | --- | --- | --- | --- | --- | --- | --- |
|  |  | From | To |  |  |  |  |  |  |  | Lower | Upper |  |
| 2 | 230,007,146 | 229,503,354 | 230,514,441 | C | T | rs62190384 | *PID1* | Intronic | 0.49 | 0.95 | 0.93 | 0.96 | 9.6E-09 |
| 5 | 4,094,165 | 3,523,315 | 4,594,165 | G | T | rs10041378 | *IRX1,LINC02114* | Intergenic | 0.24 | 0.93 | 0.91 | 0.96 | 5.6E-09 |
| 6 | 74,415,868 | 73,915,867 | 74,915,868 | A | C | rs56171536 | *CD109* | Intronic | 0.05 | 1.13 | 1.08 | 1.18 | 4.2E-08 |
| 8 | 69,431,711 | 68,931,710 | 69,931,711 | G | A | rs2380472 | *C8orf34* | Intronic | 0.13 | 0.92 | 0.89 | 0.95 | 2.5E-08 |
| 9 | 107,586,238 | 107,085,015 | 108,104,394 | C | A | rs35093463 | *ABCA1* | Intronic | 0.36 | 1.06 | 1.04 | 1.08 | 3.5E-09 |
| 11 | 203,235 | 0 | 738,058 | A | G | rs73386640 | *BET1L* | 3'UTR | 0.13 | 1.10 | 1.07 | 1.13 | 2.3E-10 |
| 12 | 10,876,573 | 10,345,946 | 11,393,295 | C | A | rs2607903 | *YBX3* | Upstream | 0.61 | 0.94 | 0.92 | 0.96 | 6.6E-10 |
| 17 | 78,358,945 | 77,612,997 | 79,262,582 | G | A | rs112735431 | *RNF213* | Nonsynonymous SNV | 0.01 | 1.61 | 1.46 | 1.78 | 2.3E-21 |
| 18 | 20,009,691 | 19,509,602 | 20,509,691 | T | C | rs9951447 | *CTAGE1,LOC101927571* | Intergenic | 0.57 | 1.06 | 1.04 | 1.08 | 4.7E-09 |

Summary statistics of novel loci that reached genome-wide significance in the Japanese GWAS (25,892 cases and 142,336 controls). CHR, Chromosome; POS, position(hg19); REF, reference allele; ALT, alternate allele; rsID, reference SNP cluster ID; AAF, alternate allele frequency; OR, odds ratio; CI, confidence interval; UTR, untranslated region.

### Very rare, high-impact signals in the genome-wide significant loci and their pleiotropy

To detect additional CAD-associated signals independent of the lead variants among the genome-wide significant loci, we performed conditional analysis for each locus of interest. This conditional analysis revealed 25 additional independent signals (locus-wide *P* < 1.0 × 10^−5^) in 48 genome-wide significant loci. They included rare variants with a high impact on the development of CAD (Fig. 1a, Supplementary Table 5). Of these genome-wide or locus-wide significant variants, we found one stop gain, five missense, and one in-frame deletion variants. In particular, a stop-gain variants (rs879255211) in the *LDLR* gene showed a dramatically high OR_CAD_ [MAF = 0.038%, OR_CAD_ (95% CI) = 4.97 (2.49 – 9.93), *P* = 5.5 × 10^−6^]. We also found two missense variants in *PCSK9* [rs151193009, MAF = 0.99%, OR_CAD_ (95% CI) = 0.64 (0.57 – 0.72), *P* = 1.21 × 10^−14^; rs564427867, MAF = 1.1%, OR_CAD_ (95% CI) = 1.31 (1.19 – 1.44), *P* = 2.7 × 10^−8^], and one in *APOB* [rs13306206, MAF = 3.8%, OR_CAD_ (95% CI) = 1.38 (1.31 – 1.46), *P* = 4.8 × 10^−35^] surpassing genome- or locus-wide significance thresholds.

**Fig. 1.**
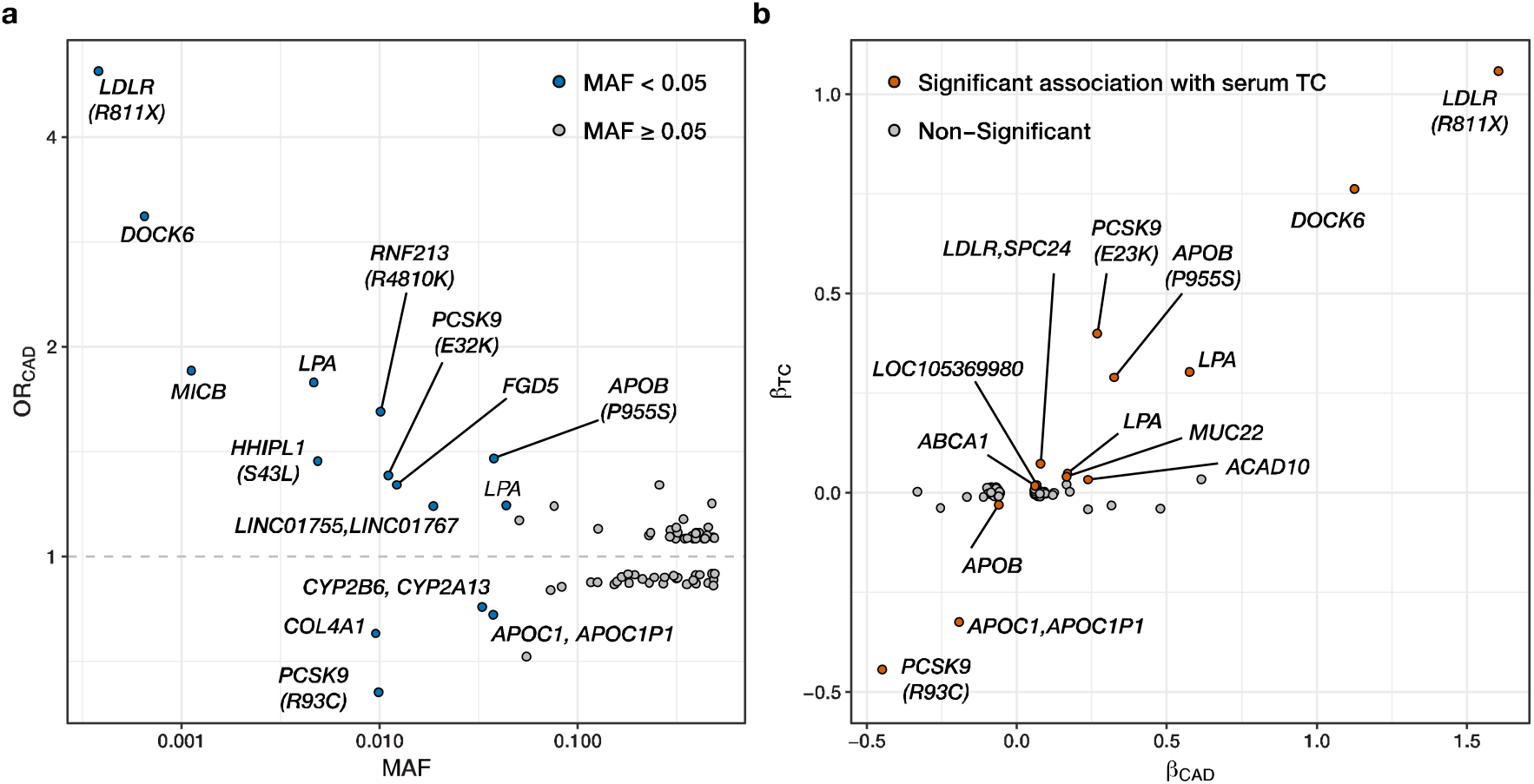
Distinct signals in CAD development and serum TC levels. **a**, The odds ratio for CAD development of the 73 independent signals in Japanese GWAS (25,892 cases, 142,336 controls) are plotted against the minor allele frequency. Rare variants (MAF < 0.05) are plotted in blue, and others in grey. **b**, Beta values for serum TC levels estimated by the linear regression model (n = 134,314) are plotted against beta values for CAD development. Variants showing significant associations for serum TC levels are plotted in orange. CAD, coronary artery disease; TC, total cholesterol, GWAS, genome-wide association study; MAF, minor allele frequency; OR, odds ratio.

To explore the biological pathways in which these independent signals play roles in CAD development, we performed association analyses for 34 clinical indices, including clinical measurements [body mass index (BMI) and blood pressure], serum laboratory measurements [e.g., total cholesterol (TC), low density lipoprotein cholesterol (LDLC)] and lifestyles (cigarette smoking and alcohol drinking) in the BBJ dataset. We found 121 significant associations (*P* < 0.05 / 2,482) in 29 phenotypes among the 34 tested (Supplementary Fig. 6, Supplementary Table 6). The most frequent association was found between serum TC and these variants. The CAD risk-increasing alleles of these variants are completely matched with the TC increased allele (14/14, Fig. 1b). Especially, rs879255211 showed a large impact on serum TC levels [MAF = 0.038%, β_TC-raw_ (standard error (SE)) = 74.6 (8.90) mg/dL per allele, *P*_TC-adjusted_ = 1.4 × 10^−10^] in accordance with its large impact on CAD development. In addition, rs553577764, a rare intronic variant in *DOCK6* with a high OR_CAD_, also showed a large impact on serum TC levels [OR_CAD_ (95% CI) = 3.08 (2.22 – 4.26), *P* = 1.2 × 10^−11^, β_TC-raw_ (SE) = 57.9 (4.30) mg/dL per allele, *P*_TC-adjusted_ = 1.2 × 10^−21^]. We noted that this variant was in high LD (R^2^ = 0.639) with rs746959386, which is a missense variant of *LDLR* registered in the ClinVar database as “Likely pathogenic” for FH [MAF = 0.057%, OR_CAD_ (95% CI) = 3.11 (2.21 – 4.37), *P* = 4.8 × 10^−11^, β_TC-raw_ (SE) = 63.4 (4.48) mg/dL per allele, *P*_TC-adjusted_ = 8.3 × 10^−21^, Supplementary Table 7]. We also noted that a splicing variant of *LDLR* showed a nominal association with CAD development [rs778408161, MAF = 0.074%, OR_CAD_ (95% CI) = 2.15 (1.44 – 3.21), *P* = 1.6 × 10^−4^] and a significant association with TC levels [β_TC-raw_ (SE) = 43.9 (4.85) mg/dL per allele, *P*_TC-adjusted_ = 2.5 × 10^−10^].

The minor allele of rs151193009 in *PCSK9* showed a protective effect against CAD development and a negative effect on serum TC levels [β_TC-raw_ (SE) = −24.3 (1.06) mg/dL per allele, *P*_TC-adjusted_ = 2.3 × 10^−111^]. This variant results in the amino acid substitution, Arg93Cys, which decreases the affinity of *PCSK9* for *LDLR*^27^, and could have negative impacts on CAD development and TC levels. In contrast, the minor allele for rs564427867 in *PCSK9* showed an increased risk for CAD development and increased TC levels [β_TC-raw_ (SE) = 26.4 (1.06) mg/dL per allele, *P*_TC-adjusted_ = 6.0 × 10^−91^]. rs13306206 is a missense variant of *APOB*, and its minor allele increased the risk of CAD development. In concordance, the risk allele of rs13306206 was associated with an increased level of serum TC [β_TC-raw_ (SE) = 19.9 (0.59) mg/dL per allele, *P*_TC-adjusted_ = 8.6 × 10^−155^]. rs112735431, a missense variant of *RNF213* that was described above, showed a significant association with systolic blood pressure (SBP) [β_SBP-raw_ (SE) = 2.48 (0.66) mmHg per allele, *P*_SBP-adjusted_ = 2.2 × 10^−22^]. Supplementary Table 8 lists the genome-wide or locus-wide associated coding variants including all of the abovementioned variants. We found that these variants were highly specific to the East Asian population and not found in the European population. Notably, rs879255211 and rs746959386 are not found even in the East Asian population in the 1000 Genomes Project^23^ or gnomAD dataset^28^.

### Individuals with rare coding variants in monogenic genes for FH showed the worse cardiovascular outcome

Epidemiological studies have shown that patients with FH have premature myocardial infarctions and worse outcomes because of recurrent ischemia^29^. However, the definition of FH in previous epidemiological studies was mainly based on clinical criteria. To confirm the association between clinical outcomes and deleterious variants in established FH genes, we assessed the association of the carrier status of the variants detected in the current study (rs564427867 in *PCSK9*, MAF = 1.1%; rs13306206 in *APOB*, MAF = 3.8%, and rs879255211, rs746959386, and rs778408161 in *LDLR*, MAF = 0.038%, 0.057%, and 0.074%, respectively) with clinical indicators. We found that these variants are significantly enriched in the patients with acute coronary syndrome [ACS, defined as a composite of acute myocardial infarction (AMI) and unstable angina] compared to those with stable angina pectoris (SAP) [OR_ACS/SAP_ (95% CI) = 1.23 (1.14 – 1.33) per allele. *P* = 1.2 × 10^−7^, Fig. 2a, Supplementary Fig. 7a]; ACS tends to be associated with a more aggressive clinical course than SAP. In addition, individuals with these variants developed AMI at a younger age than non-carriers [effect size on onset age per allele (95% CI) = −1.52 (−2.06; −0.98), *P* = 4.7 × 10^−8^, Fig. 2b]. In concordance with these results, as illustrated in Fig. 2c, carriers of these variants showed significantly increased cardiovascular mortality than non-carriers [adjusted hazard ratio (HR) = 1.16, 95% CI = 1.07 – 1.27, *P* = 3.5 × 10^−4^, Supplementary Fig. 7b).

**Fig. 2.**
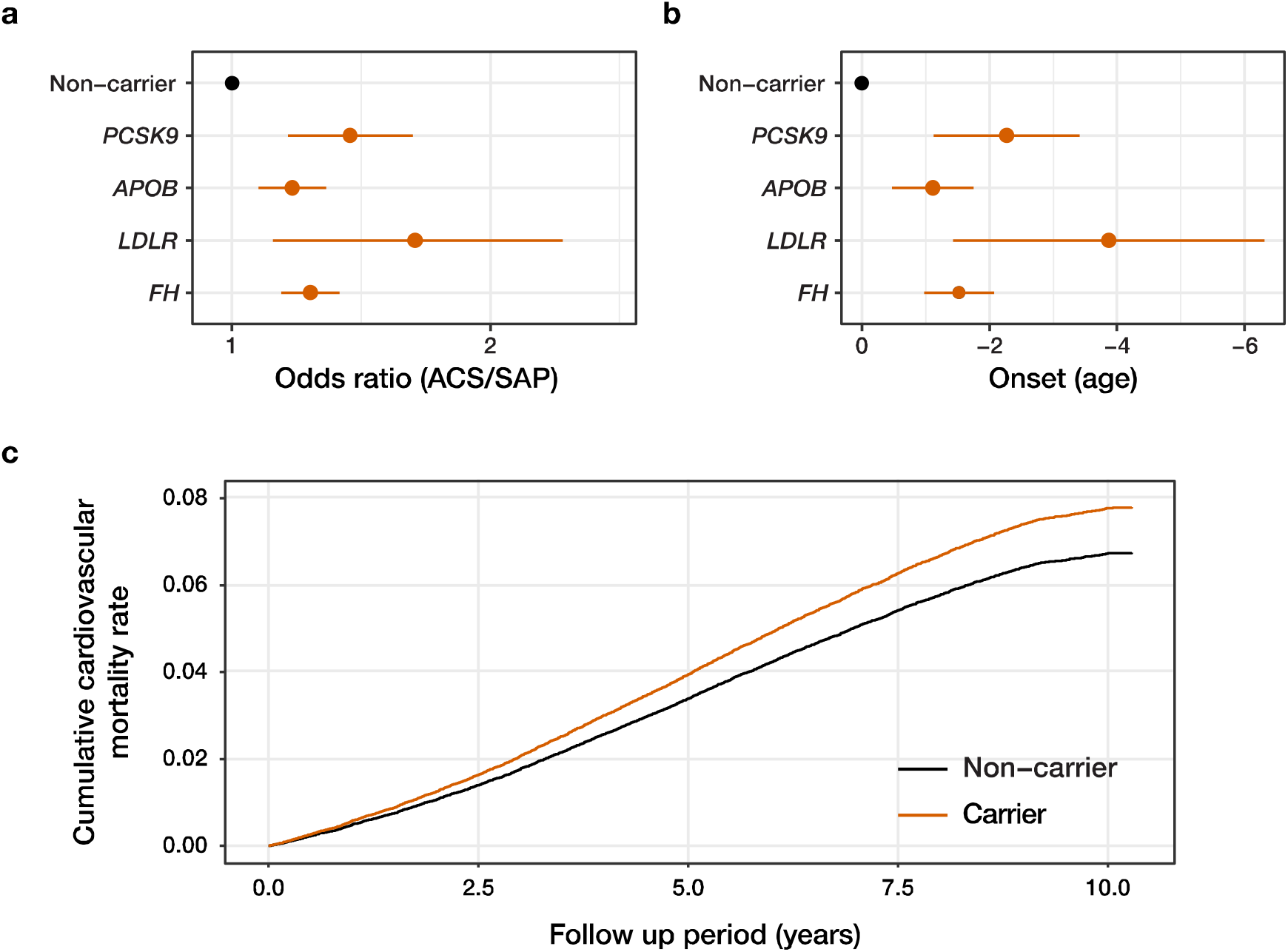
Impact of the deleterious variant in FH genes in CAD subtypes, age of onset of AMI, and long-term cardiovascular mortality. **a**, Each point indicates the odds ratio for developing ACS with the error bar indicating its 95% confidence intervals for alternate allele dosage of deleterious variants for each gene. Individuals with SAP were used as controls. **b**, Each point indicates effect sizes for the onset age of AMI of alternate allele dosage with the error bar indicating the 95% confidence interval. **c**, Adjusted curves for mortality from diseases of the circulatory system (ICD10.I) stratified by carrier status of deleterious variants in established FH genes (rs564427867 in *PCSK9*; rs13306206 in *APOB*; rs879255211, rs746959386, and rs778408161 in *LDLR*) are shown. ACS, acute coronary syndrome; SAP, stable angina pectoris; AMI, acute myocardial infarction; FH, familial hypercholesterolaemia.

### Transethnic meta-analysis identified 37 novel CAD-associated loci

To increase the power for detecting further associations with CAD, we conducted a transethnic meta-analysis combining the current Japanese GWAS data and previously published data from two large-scale CAD GWAS, CardiogramPlusC4D (C4D) and UK Biobank (UKBB), mainly involving subjects of European descent (Supplementary Fig. 1b)^7,9^. To account for the ancestral heterogeneity in each study, we applied the MANTRA algorithm in the analysis^30^. By combining all three datasets, there was a total of 121,234 CAD cases (BBJ: 25,892, C4D: 60,801, UKBB: 34,541) and 527,824 controls (BBJ: 142,336, C4D: 123,504, UKBB: 261,984). A total of 4,804,024 SNPs was tested and 176 loci reached the genome-wide significance threshold [log_10_ Bayes factor (BF) > 6, Fig. 3, Supplementary Table 9, Supplementary Data Set 3,4]^31^. Forty-three of these loci were not previously reported (Table 2), including six loci that we detected in the current Japanese GWAS. In total, we found 46 previously unreported loci in the Japanese GWAS and the transethnic meta-analysis. Among these loci, we found a novel association on chromosome 5q13, which harbours *HMGCR* encoding the late limiting enzyme in endogenous cholesterol synthesis, 3-hydroxy-3-methylglutaryl coenzyme A (HMG-CoA) reductase. This is the target enzyme of statins, the most prevalent lipid lowering agents. The lead variant rs13354746 is located in 13kb upstream of *HMGCR*, and the risk allele of this variant is highly associated with increased serum TC levels in the Japanese population^32^.

**Fig. 3.**
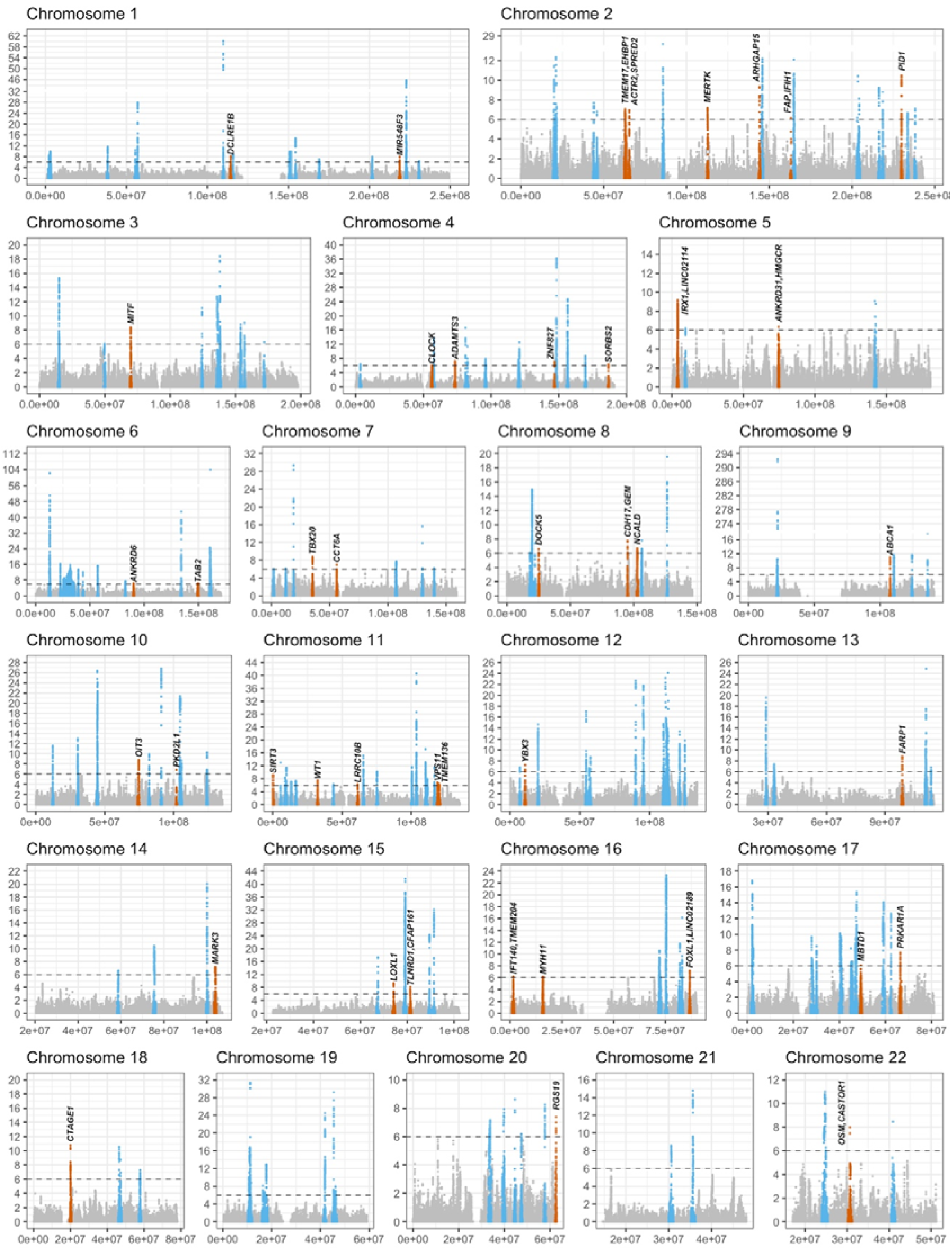
Manhattan plots for transethnic meta-analysis. The results of the case-control association study in the transethnic meta-analysis (CAD case 121,234, control 527,824) are shown. The log_10_BF on the y-axes are plotted against the genomic position (hg19) on the x-axes. Variants in 43 novel and 133 previously reported loci are presented in orange and blue, respectively. Dashed lines indicate genome wide significant thresholds (Log_10_BF = 6). BF, Bayes Factor.

**Table 2.** Novel loci identified by Transethnic meta-analysis.

| CHR | POS | Locus |  | REF | ALT | rsID | Gene | Annotation | MANTRA |  |  | Fixed effect |  |  |  |  |
| --- | --- | --- | --- | --- | --- | --- | --- | --- | --- | --- | --- | --- | --- | --- | --- | --- |
|  |  | From | To |  |  |  |  |  | Beta | SE | Log<sub>10</sub> BF | Beta | SE | P | Q | P<sub>Q</sub> |
| 1 | 114,448,389 | 113,926,000 | 114,950,964 | C | T | rs11552449 | *DCLRE1B* | Nonsynonymous SNV | 0.037 | 0.006 | 8.1 | 0.037 | 0.006 | 3.7E-10 | 0.2 | 8.9E-01 |
| 1 | 218,827,855 | 218,317,531 | 219,358,746 | T | C | rs1481345 | *MIR548F3* | ncRNA Intronic | -0.031 | 0.005 | 7.9 | -0.031 | 0.005 | 4.7E-10 | 2.0 | 3.7E-01 |
| 2 | 62,878,928 | 62,357,974 | 63,679,076 | G | C | rs7420881 | *TMEM17, EHBP1* | Intergenic | 0.032 | 0.005 | 7.1 | 0.032 | 0.005 | 3.9E-09 | 0.5 | 7.7E-01 |
| 2 | 65,499,468 | 64,982,115 | 66,030,347 | G | A | rs72822411 | *ACTR2, SPRED2* | Intergenic | 0.036 | 0.006 | 6.9 | 0.036 | 0.006 | 6.1E-09 | 3.1 | 2.2E-01 |
| 2 | 112,656,652 | 112,155,241 | 113,263,025 | G | A | rs7604403 | *MERTK* | Intronic | 0.034 | 0.006 | 7.2 | 0.034 | 0.006 | 3.5E-09 | 1.5 | 4.7E-01 |
| 2 | 144,158,418 | 143,651,523 | 144,698,216 | T | C | rs12469628 | *ARHGAP15* | Intronic | -0.040 | 0.006 | 9.3 | -0.041 | 0.006 | 1.9E-11 | 1.2 | 5.5E-01 |
| 2 | 163,110,536 | 162,610,535 | 163,610,536 | A | G | rs2111485 | *FAP, IFIH1* | Intergenic | 0.029 | 0.005 | 6.2 | 0.029 | 0.005 | 3.7E-08 | 0.9 | 6.5E-01 |
| 2 | 230,009,317 | 229,480,420 | 230,515,954 | C | T | rs7566501 | *PID1* | Intronic | -0.035 | 0.005 | 10.5 | -0.035 | 0.005 | 1.9E-12 | 7.2 | 2.7E-02 |
| 3 | 69,820,782 | 69,302,901 | 70,409,425 | T | C | rs12714757 | *MITF* | Intronic | -0.036 | 0.006 | 8.4 | -0.036 | 0.006 | 1.5E-10 | 2.2 | 3.3E-01 |
| 4 | 56,316,979 | 55,816,978 | 56,816,979 | T | C | rs11133381 | *CLOCK* | Intronic | -0.028 | 0.005 | 6.0 | -0.027 | 0.005 | 4.8E-08 | 2.1 | 3.6E-01 |
| 4 | 73,420,634 | 72,919,494 | 73,984,862 | G | A | rs13105983 | *ADAMTS3* | Intronic | -0.029 | 0.005 | 7.3 | -0.029 | 0.005 | 2.8E-09 | 3.4 | 1.8E-01 |
| 4 | 146,809,998 | 146,259,606 | 147,321,725 | T | G | rs10006310 | *ZNF827* | Intronic | -0.029 | 0.005 | 7.2 | -0.029 | 0.005 | 3.2E-09 | 1.0 | 6.0E-01 |
| 4 | 186,692,853 | 186,190,488 | 187,193,134 | T | C | rs7680806 | *SORBS2* | Intronic | 0.037 | 0.007 | 6.4 | 0.037 | 0.007 | 3.0E-08 | 4.9 | 8.4E-02 |
| 5 | 4,023,730 | 3,506,098 | 4,600,989 | C | T | rs548581 | *IRX1, LINC02114* | Intergenic | 0.022 | 0.005 | 9.2 | 0.022 | 0.005 | 1.6E-05 | 31.0 | 1.8E-07 |
| 5 | 74,619,132 | 74,119,131 | 75,119,132 | C | T | rs13354746 | *ANKRD31, HMGCR* | Intergenic | 0.031 | 0.006 | 6.4 | 0.032 | 0.006 | 2.5E-08 | 1.5 | 4.7E-01 |
| 6 | 90,314,917 | 89,814,916 | 90,829,895 | A | G | rs9351209 | *ANKRD6* | Intronic | -0.029 | 0.005 | 6.4 | -0.029 | 0.005 | 1.8E-08 | 3.4 | 1.8E-01 |
| 6 | 149,714,790 | 149,153,791 | 150,244,924 | G | T | rs2744427 | *TAB2* | Intronic | -0.036 | 0.007 | 6.3 | -0.036 | 0.007 | 2.4E-08 | 3.0 | 2.3E-01 |
| 7 | 35,286,471 | 34,769,624 | 35,802,478 | C | A | rs4723406 | *TBX20* | Intronic | -0.033 | 0.005 | 8.8 | -0.033 | 0.005 | 7.9E-11 | 0.7 | 7.0E-01 |
| 7 | 56,122,058 | 55,622,057 | 56,622,058 | A | T | rs6593297 | *CCT6A* | Intronic | -0.031 | 0.005 | 7.0 | -0.031 | 0.005 | 4.0E-09 | 0.3 | 8.5E-01 |
| 8 | 25,064,984 | 24,561,806 | 25,564,984 | G | A | rs9693598 | *DOCK5* | Intronic | -0.044 | 0.008 | 6.6 | -0.044 | 0.008 | 1.6E-08 | 0.1 | 9.4E-01 |
| 8 | 95,260,225 | 94,750,557 | 95,781,052 | A | G | rs2623168 | *CDH17, GEM* | Intergenic | 0.047 | 0.008 | 7.8 | 0.047 | 0.008 | 1.1E-09 | 6.7 | 3.5E-02 |
| 8 | 102,832,405 | 102,313,394 | 103,395,016 | C | T | rs13255004 | *NCALD* | Intronic | 0.034 | 0.006 | 6.7 | 0.034 | 0.006 | 1.1E-08 | 3.8 | 1.5E-01 |
| 9 | 107,586,238 | 107,085,015 | 108,104,394 | C | A | rs35093463 | *ABCA1* | Intronic | 0.054 | 0.007 | 11.0 | 0.055 | 0.008 | 3.4E-13 | 2.3 | 3.2E-01 |
| 10 | 74,682,633 | 73,808,317 | 75,191,035 | T | G | rs11000448 | *OIT3* | Intronic | -0.053 | 0.008 | 8.8 | -0.052 | 0.008 | 3.9E-11 | 2.4 | 3.0E-01 |
| 10 | 102,075,479 | 101,575,478 | 102,575,479 | G | A | rs603424 | *PKD2L1* | Intronic | 0.040 | 0.007 | 6.9 | 0.040 | 0.007 | 5.8E-09 | 0.0 | 9.9E-01 |
| 11 | 224,845 | 0 | 738,058 | G | C | rs73392700 | *SIRT3* | Intronic | 0.064 | 0.010 | 9.2 | 0.064 | 0.010 | 3.3E-11 | 6.5 | 4.0E-02 |
| 11 | 32,441,377 | 31,875,668 | 32,941,377 | C | T | rs7115190 | *WT1* | Intronic | 0.032 | 0.005 | 7.5 | 0.032 | 0.005 | 1.6E-09 | 1.0 | 6.1E-01 |
| 11 | 61,277,698 | 60,777,697 | 61,778,918 | A | G | rs11230728 | *LRRC10B* | 3'UTR | -0.036 | 0.006 | 6.3 | -0.036 | 0.006 | 3.6E-08 | 1.5 | 4.8E-01 |
| 11 | 118,950,217 | 118,428,252 | 119,450,217 | C | G | rs1307145 | *VPS11* | Intronic | 0.031 | 0.005 | 6.8 | 0.031 | 0.005 | 7.2E-09 | 2.5 | 2.8E-01 |
| 11 | 120,198,093 | 119,698,092 | 120,847,715 | G | A | rs1893261 | *TMEM136* | Synonymous SNV | 0.028 | 0.005 | 6.3 | 0.028 | 0.005 | 3.1E-08 | 2.4 | 3.0E-01 |
| 12 | 10,876,573 | 10,376,572 | 11,392,139 | C | A | rs2607903 | *YBX3* | Upstream | -0.040 | 0.007 | 7.2 | -0.040 | 0.007 | 4.8E-09 | 9.3 | 9.7E-03 |
| 13 | 98,859,335 | 98,357,228 | 99,366,535 | G | A | rs9556903 | *FARP1* | Intronic | -0.044 | 0.007 | 8.8 | -0.044 | 0.007 | 7.9E-11 | 6.2 | 4.5E-02 |
| 14 | 103,900,481 | 103,349,714 | 104,484,616 | A | G | rs35224956 | *MARK3* | Intronic | 0.031 | 0.005 | 7.2 | 0.031 | 0.005 | 3.2E-09 | 0.7 | 7.2E-01 |
| 15 | 74,223,716 | 73,721,297 | 74,782,343 | G | C | rs28522673 | *LOXL1* | Intronic | 0.044 | 0.006 | 9.4 | 0.044 | 0.007 | 1.8E-11 | 4.5 | 1.1E-01 |
| 15 | 81,377,717 | 80,834,756 | 81,898,791 | C | A | rs1879454 | *TLNDR1, CFAP161* | Intergenic | -0.036 | 0.006 | 8.1 | -0.036 | 0.006 | 4.5E-10 | 3.7 | 1.6E-01 |
| 16 | 1,584,618 | 1,084,617 | 2,084,866 | T | C | rs2076438 | *IFT140, TMEM204* | Intronic | 0.028 | 0.005 | 6.2 | 0.028 | 0.005 | 3.1E-08 | 0.9 | 6.4E-01 |
| 16 | 15,917,838 | 15,412,982 | 16,417,838 | C | G | rs216158 | *MYH11* | Intronic | -0.029 | 0.005 | 6.1 | -0.029 | 0.005 | 3.9E-08 | 0.1 | 9.6E-01 |
| 16 | 86,699,163 | 86,198,030 | 87,216,058 | A | G | rs12444314 | *FOXL1, LINC02189* | Intergenic | 0.032 | 0.006 | 7.2 | 0.032 | 0.005 | 3.6E-09 | 0.2 | 9.1E-01 |
| 17 | 49,308,707 | 48,808,706 | 49,808,707 | C | T | rs4794213 | *MBTD1* | Intronic | -0.028 | 0.005 | 6.1 | -0.028 | 0.005 | 3.6E-08 | 0.3 | 8.7E-01 |
| 17 | 66,469,400 | 65,953,304 | 66,970,892 | T | G | rs2952286 | *PRKAR1A* | Intronic | 0.035 | 0.006 | 7.7 | 0.035 | 0.006 | 5.3E-10 | 3.6 | 1.6E-01 |
| 18 | 19,998,810 | 19,498,809 | 20,584,756 | G | C | rs948386 | *CTAGE1* | Upstream | 0.036 | 0.005 | 10.8 | 0.036 | 0.005 | 8.1E-13 | 5.3 | 7.2E-02 |
| 20 | 62,709,274 | 62,190,682 | 63,025,520 | G | A | rs12625329 | *RGS19* | Intronic | -0.031 | 0.005 | 7.4 | -0.031 | 0.005 | 2.1E-09 | 2.8 | 2.5E-01 |
| 22 | 30,669,883 | 30,167,276 | 31,169,883 | G | A | rs6006426 | *OSM, CASTOR1* | Intergenic | 0.030 | 0.005 | 8.0 | 0.031 | 0.005 | 4.1E-10 | 5.7 | 5.7E-02 |

Summary statistics of novel loci that reached genome-wide significance in the transethnic meta-analysis (121,234 cases and 527,824 controls). CHR, Chromosome; POS, position(hg19); REF, reference allele; ALT, alternate allele; rsID, reference SNP cluster ID; SE, standard error; BF, Bayes factor; Q, Cochran's Q value; P<sub>Q</sub>, P value for Chochran's Q test; SNV, single nucleotide variant; ncRNA, non-coding RNA.

### Transethnic fine mapping of CAD-associated loci

For the fine mapping of the CAD-associated loci, we constructed credible sets for the genome-wide significant loci detected in the transethnic meta-analysis. To assess the contribution of the transethnic meta-analysis to the sizes of the credible sets, we compared the numbers of variants included in the 99% credible sets derived from the transethnic meta-analysis (BBJ, C4D, and UKBB) and the meta-analysis of the two previous European studies (C4D and UKBB, Supplementary Table 10). The sizes of the 99% credible sets in the previously established loci derived from the transethnic meta-analysis were significantly decreased compared to those from the European only analysis [Median number of variants (1st– 3rd quantile value) = 13.5 (5 – 33.5) and 19 (6 – 55.5), respectively, *P* = 8.7 × 10^−9^, paired Wilcoxon rank-sum test, Supplementary Fig. 8]. We found 28 and 55 lead variants with a posterior probability of association (PPA) greater than 80% and 50%, respectively (Supplementary Table 11), including four coding variants (rs11556924 in *ZC3HC1*, rs11601507 in *TRIM5*, rs3741380 in *EHBP1L1*, and rs1169288 in *HNF1A*). A previous study implicated three of these variants (rs11556924, rs11601507, and rs3741380) as a causal variant for each locus^9^. rs1169288, a non-synonymous variant in *HNF1A* (Ile27Leu), which encodes a crucial transcription factor highly expressed in the liver and intestine^33^, was also previously implicated in cholesterol homeostasis and glucose tolerance^32,34^, and was predicted to be highly protein damaging (CADD PHREAD score of 23.4)^35^.

### Shared allelic effects in the transethnic meta-analysis, and derivation of the transethnic PRS

To assess the heterogeneity of allelic effects on CAD development among the studies, we next compared the alternate allele frequency (AAF) and β_CAD_ between the Japanese and two European studies (Supplementary Fig. 9). We observed considerably different allele frequencies between the Japanese and European studies for the 176 lead variants detected in the transethnic meta-analysis. Nevertheless, we found a significant positive correlation and concordant allelic effects of these variants (Spearman’s ρ_BBJ-C4D_ = 0.81, *P*_BBJ-C4D_ = 6.2 × 10^−42^; ρ_BBJ-UKBB_ = 0.79, *P*_BBJ-UKBB_ = 8.0 × 10^−39^. directional consistencies were 95.5% for BBJ vs. C4D and 94.9% for BBJ vs. UKBB).

For further exploration of allelic effect consistency, we compared the allelic effect directions of the variants with various significances (Supplementary Fig. 10). We observed strong correlations of allelic effect direction in the variants with genome-wide significance (concordant rate of allelic effect was 95.9% for variants with *P*_fixed-effect_ < 5 × 10^−8^, 100% for *P*_random-effect_ < 5 × 10^−8^) as mentioned above. Moreover, we found significant concordance of allelic effect direction in variants even with nominal significance (82.0% for *P*_fixed-effect_ < 0.05, 93.7% for *P*_random-effect_ < 0.05). In addition, we noted that the random-effect model consistently gives better estimates for variants with shared allelic effect across these P-value thresholds.

The transethnic meta-analysis substantially increased the number of significant associations and suggested a shared allelic effect among ethnicities. These results indicate that transethnic meta-analysis could improve the performance of a PRS. To determine the best PRS under such circumstances, we derived the PRS exhaustively from all combinations of summary statistics, reference LD structure, and parameters for derivation. As a result, the random-effects transethnic meta-analysis showed the best performance [Nagelkerke’s R^2^ = 0.0776, the area under the receiver-operator curve (AUC) = 0.664, OR = 8.30, Supplementary Fig. 11, Supplementary Table 12], in the independent Japanese case-control validation cohort (1,827 cases, 9,172 controls). This transethnic CAD-PRS outperformed the previously derived CAD-PRS from a European study^16^ (Nagelkerke’s R^2^ = 0.0479, AUC = 0.628, OR = 3.70, *P* = 1.3 × 10^−6^ vs. transethnic CAD-PRS) or the PRS derived from the current Japanese GWAS (Nagelkerke’s R^2^ = 0.0498, AUC = 0.634, OR = 4.24, *P* = 7.1 × 10^−7^, vs. transethnic CAD-PRS).

### Associations of CAD-PRS with clinical risk factors

To determine the clinical pathways explaining the involvement of the CAD-PRS in CAD pathophysiology, we assessed the correlation between the CAD-PRS and various phenotypes as denoted above. Among the 34 variables tested, significant correlations were observed between 13 clinical factors and CAD-PRS after multiple-testing correction (Fig. 4a, Supplementary Fig. 12, Supplementary Fig. 13, Supplementary Table 13). Most of the traditional CAD risk factors, including blood pressure and lipid or diabetic measurements, showed significant correlations with the CAD-PRS. SBP showed the most significant positive correlation [Spearman’s ρ (95% CI) = 0.052 (0.042 – 0.061), *P* = 1.9 × 10^−24^], and HDLC showed the third most significant, negative correlation as expected from its clinical consequence [ρ (95% CI) = −0.045 (−0.057; −0.033), *P* = 2.9 × 10^−13^]. The CAD-PRS was also significantly correlated with inflammatory markers [white blood cell count (WBC), ρ (95% CI) = 0.018 (0.008 – 0.027), *P* = 3.2 × 10^−4^], alcohol drinking [β (95% CI) = −0.048 (−0.069; −0.027), *P* = 6.8 × 10^−6^], and cigarette smoking [β (95% CI) = 0.040 (0.016 – 0.065), P = 1.3 × 10^−3^].

**Fig. 4.**
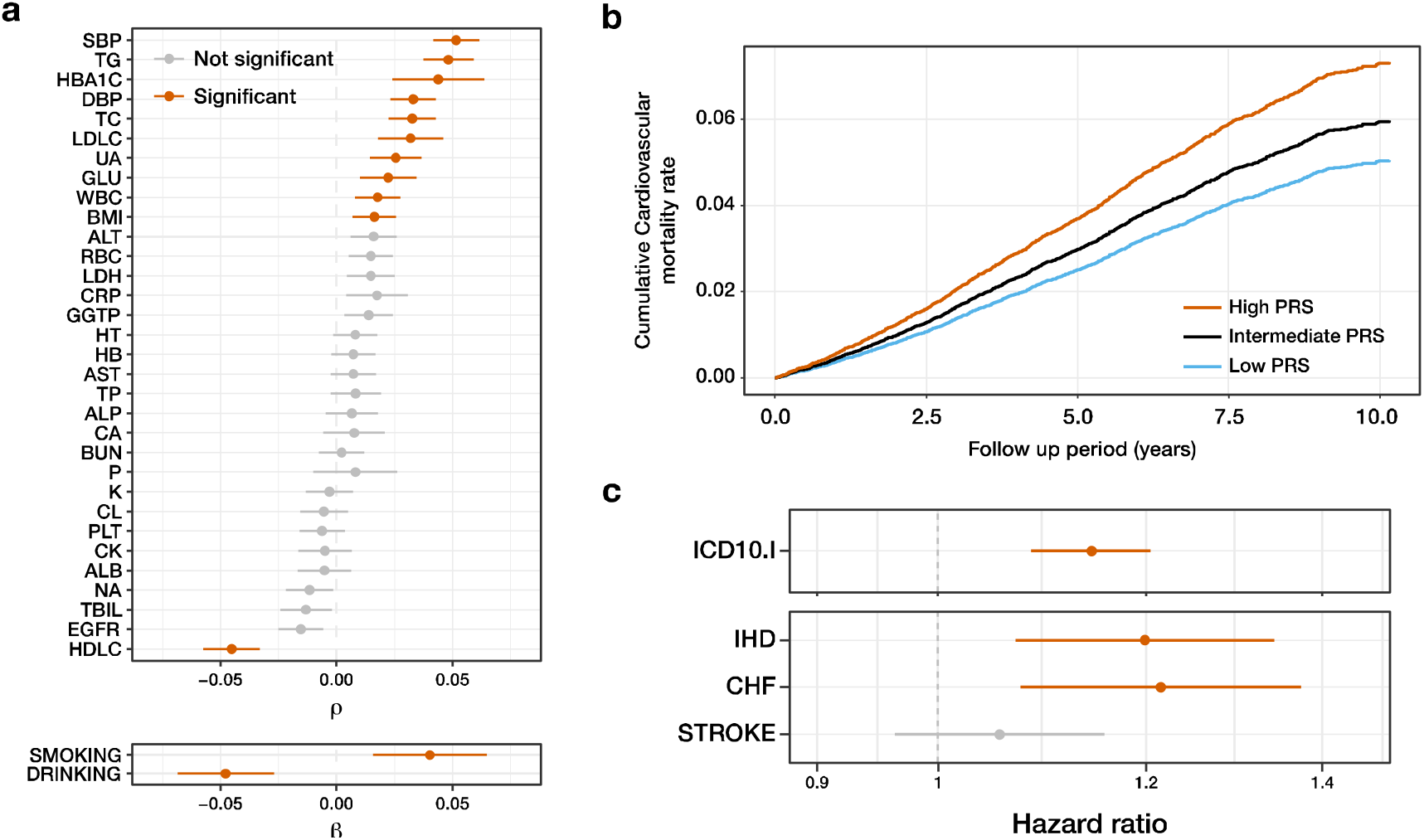
Correlation between transethnic CAD-PRS and clinical indices. **a**, Upper panel; Each point indicates Spearman’s correlation coefficient between CAD-PRS and clinical indices. Error bars indicates 95% confidence intervals. Lower panel; Each point indicates a beta coefficient for 1SD increase in CAD-PRS estimated by the logistic regression model. Significant correlations or associations are shown in orange (*P* < 0.05/34). **b**, Adjusted curves for mortality from ICD10.I diseases estimated by Cox’s proportional hazard model are shown. Individuals are stratified into high PRS (top 20 percentile, orange), low PRS (bottom 20 percentile, blue), and intermediate PRS (others, black). **c**, Each point indicates a hazard ratio of 1SD increase in CAD-PRS for mortality from ICD10.I subtypes. Error bars represents the 95% confidence interval. PRS, polygenic risk score; ICD10, International Statistical Classification of Diseases and Related Health Problems 10th Revision; IHD, ischemic heart disease; CHF, congestive heart failure; SD, standard deviation. Abbreviations of other phenotypes are defined in Supplementary Table 17.

### CAD-PRS and cardiovascular mortality

Considering the strong correlation between the CAD-PRS and CAD risk factors (e.g., blood pressure, serum lipid profiles), we hypothesized that CAD-PRS reflects the severity of the genetic background for CAD development, and as a result, the individuals who have a high CAD-PRS will have a poor prognosis. To test this hypothesis, we assessed the impact of the CAD-PRS on mortality in long-term follow-up data. As illustrated in Fig. 4b, individuals with high CAD-PRS showed significantly increased mortality from diseases classified as circulatory system-related [International Statistical Classification of Diseases (ICD)10-I, HR for increasing the CAD-PRS by 1 standard deviation (95% CI) = 1.14 (1.09 – 1.20), *P* = 4.4 × 10^−7^, Supplementary Fig. 14, Supplementary Table 14]. Of note, this association was highly specific to circulatory diseases, and other causes of death were not associated with the CAD-PRS. We found significant association between CAD-PRS and all-cause mortality [HR (95% CI) = 1.05 (1.02 – 1.07), *P* = 3.4 × 10^−4^], but, when we excluded ICD10-I disease from all-cause mortality, the significance was diminished [HR (95% CI) = 1.02 (0.99 – 1.05), *P* = 0.183]. Furthermore, we divided individuals who died from ICD10-I into three sub-categories: ischemic heart disease (ICD-10 I21-I25), congestive heart failure (ICD-10 I50), and stroke (ICD-10 I60-I69). Two of them showed significant associations with CAD-PRS (i.e., HR (95% CI) = 1.20 (1.07 – 1.34), *P* = 1.6 × 10^−3^ for ischemic heart disease, HR (95% CI) = 1.22 (1.08 – 1.37), *P* = 1.8 × 10^−3^ for congestive heart failure, HR (95% CI) = 1.06 (0.96 – 1.16), *P* = 0.25 for stroke, Fig. 4c, Supplementary Table 14).

### Genetic basis of CAD-PRS-associated traits

Finally, to assess the related loci of these CAD-PRS associated traits, we performed association analyses for the 176 lead variants detected in the transethnic meta-analysis and the 13 CAD-PRS-associated traits in the BBJ dataset. Ninety-seven variant-phenotype pairs showed Bonferroni-corrected significance (*P* < 0.05/2,288, Supplementary Fig. 15, Supplementary Table 15). Serum TC level was the most frequently associated trait (18 associations) and the directions of allelic effect (CAD-risk and TC-increasing) were completely concordant (18/18). To further characterize the relationships between the CAD-associated loci and pleiotropic effects on CAD risk factors, we performed unsupervised clustering for the Z-value matrix of these variants, revealing distinct functional clusters (Supplementary Fig. 16). The loci with positive impacts on serum TC or LDLC levels segregated in Cluster 1. This cluster contains well-characterized loci associated with serum lipid profiles, exemplified by *PCSK9, APOB*, *HMGCR*, and *LDLR*. Cluster 2 harbours the loci associated with glycaemic traits. The loci in Cluster 3 positively impact on WBCs; the 9p21 (*CDKN2B-AS1*) locus is included in this cluster. The loci in cluster 4 positively impact serum triglyceride levels and negatively impact HDLC levels; this cluster also contains *APOA5* and *LPL* loci, which are crucial genes for lipoprotein metabolism. Cluster 5 includes lead variants that affect BMI. Cluster 6, the second largest cluster, contains loci associated with blood pressure.

## Discussion

We performed a large-scale CAD GWAS in the Japanese population in combination with whole-genome sequencing, transethnic meta-analysis, and analyses of various types of biobank-based datasets, including clinical phenotypes and long-term follow-up data. In addition to discovering 46 novel loci, we confirmed the clinical relevance of the CAD-PRS to survival prediction and its association with various CAD-associated phenotypes.

The use of a population- and disease-specific reference haplotypes improved the imputation quality and enabled assessments of very rare variants with a high impact^36–38^. In the current study, we found significant associations between a loss-of-function variant in *LDLR* and a high OR for CAD. This variant was not even found in the large-scale sequencing database (gnomAD) that included a substantial number of individuals of East Asian ancestry. In addition, such disease-relevant rare variants identified in this study are highly population specific (Supplementary table 8). These findings warrant further effort for sequencing of diseased individuals in diverse populations to explore disease mechanisms and molecular targets. Although the contribution of these rare and very rare variants to the total disease heritability is small, accurate information on the effects of these variants is essential to provide better medical advice or genetic counselling for the carriers. Therefore, continuous efforts should be devoted to research on the behaviour and the impact of such rare variants in the population.

In addition to these population-specific rare variants, we found 176 genome-wide associated loci for CAD including 43 novel loci in the transethnic meta-analysis. The allelic effects of these lead variants showed almost the same directionality between our Japanese GWAS and previously reported European GWAS. This result coincides with previous findings that allelic effects of complex diseases or traits are shared between individuals of East Asian and European descent^19,20^. Moreover, we observed that a large proportion of variants with nominal significance showed directional consistency. This shared allelic effect enabled us to derive a CAD-PRS from a transethnic meta-analysis for Japanese, which outperformed those derived from either Japanese or European GWAS results. Recently, it was reported that a PRS derived from a specific ethnic group could not achieve the same performance with other ethnic groups^39^. Before the implementation of PRS in clinical practice or public health policies, it is essential to develop PRS that are impartial and applicable to diverse populations. Therefore, it is crucial to develop a reliable method to share GWAS results between different ethnic groups effectively. Our results indicate that a transethnic meta-analysis could assist in the extrapolation of existing data that was obtained with a different ancestry.

In the phenome-wide analysis, we found significant dose-dependent relationships between the CAD-PRS and 13 CAD-relevant traits. This result suggests that the CAD-PRS not only distinguishes case-control status as a binary trait but also reflects the severity of the genetic background of CAD. Supporting this hypothesis, survival analysis revealed tight relationships between the CAD-PRS and mortality in long-term follow-up data. These observations support the feasibility of polygenic risk stratification in non-European populations, especially utilizing the data from European populations, and this warrants further follow-up study.

In conclusion, our large-scale genetic analysis allowed us to revise the genetic architecture of CAD and revealed 46 previously unreported loci associated with CAD susceptibility. The transethnic analysis also significantly improved the performance of the CAD-PRS compared to that of population-specific ones. Moreover, biobank-based phenome-wide analysis and survival analysis revealed the behaviour of CAD-PRS in clinical settings through the correlation with clinical risk factors and survival implications. These data provide a fundamental resource for future research, and the clinical implementation of CAD genetics for precision medicine.

### URLs

BioBank Japan, https://biobankjp.org; Nagahama cohort, http://zeroji-cohort.com; JPHC, https://epi.ncc.go.jp; J-MICC, http://www.jmicc.com; CARDIoGRAMplusC4D, http://www.cardiogramplusc4d.org; Cardiovascular Disease Knowledge portal; http://www.broadcvdi.org; UK Biobank, http://www.ukbiobank.ac.uk; HapMap project, http://hapmap.ncbi.nlm.nih.gov; 1000 Genomes Project, http://www.1000genomes.org; gnomAD, https://gnomad.broadinstitute.org; ClinVar, https://www.ncbi.nlm.nih.gov/clinvar; LDSC, https://github.com/bulik/ldsc/; Eagle, https://data.broadinstitute.org/alkesgroup/Eagle; Minimac3, https://genome.sph.umich.edu/wiki/Minimac3; PLINK, https://www.cog-genomics.org/plink/1.9; METASOFT, http://genetics.cs.ucla.edu/meta; ANNOVAR, http://annovar.openbioinformatics.org; LocusZoom, http://locuszoom.sph.umich.edu; NBDC Human Database, https://humandbs.biosciencedbc.jp.

## Online Methods

### Subjects

Biobank Japan (BBJ)^40,41^ is a hospital-based Japanese national biobank project including data of approximately 200,000 patients enrolled between 2003 and 2007. Participants were recruited at 12 medical institutes throughout Japan (Osaka Medical Center for Cancer and Cardiovascular Diseases, the Cancer Institute Hospital of Japanese Foundation for Cancer Research, Juntendo University, Tokyo Metropolitan Geriatric Hospital, Nippon Medical School, Nihon University School of Medicine, Iwate Medical University, Tokushukai Hospitals, Shiga University of Medical Science, Fukujuji Hospital, National Hospital Organization Osaka National Hospital, and Iizuka Hospital). The Nagahama study is a community-based cohort study, conducted in Shiga, Japan. Participants were recruited from the general population aged 30 – 74 years in Nagahama city from 2008 to 2010. The Japan Public Health Center-based Prospective Study (JPHC)^42^ is an ongoing community-based prospective study conducted in 11 public health center areas nationwide since 1990. The JPHC enrolled residents aged 40 – 69 years. Japan Multi-Institutional Collaborative Cohort (J-MICC) Study, community dwellers aged 35 – 69 years were recruited between 2005 and 2013 in 13 study areas throughout Japan. The Osaka Acute Coronary Insufficiency Study (OACIS) is a hospital-based registry in which patients with acute myocardial infarction were enrolled at Osaka University and 24 collaborating hospitals in the Osaka-Hyogo area from 1998 to 2014. In the Informed consent was obtained from all participants in each study. Our study was approved by the relevant ethical committees at each facility.

### Whole-genome sequencing, quality control, and construction of reference panels

We sequenced 1,782 samples from patients with early-onset CAD (cases) and 3,148 controls. Whole-genome sequencing was performed on the HiSeqX5 platform aiming at a 15 × depth, using 2 × 150-bp paired-end reads. All case samples were obtained from the BBJ cohort (n = 1,782), and control samples were obtained from the BBJ (n = 1,007) or Nagahama (n = 2,141) cohort. Sequenced data were processed using Picard and aligned to the hs37d5 reference genome in the BBJ cohort and to hg19 in the Nagahama cohort using the Burrows-Wheeler algorithm. Genotypes of the samples were called individually in each centre using the HaplotypeCaller according to Genome Analysis Toolkit best practice for germline SNPs and indels. Per-sample GVCF genotype data were merged and jointly called using GenotypeGVCFs. We defined exclusion filters for genotypes as follows: (1) filtered depth (DP) < 5 and (2) quality of the assigned genotype (GQ) < 20. We set these genotypes as missing and excluded variants with call rates < 90% before variant quality score recalibration (VQSR). After VQSR filtering, sample quality control was performed by excluding samples with excess heterozygosity, excess singletons, and closely related samples estimated based on identity by states (PIHAT > 0.2). Principal component analysis restricted samples in the Japanese mainland cluster. After excluding these samples, 1,781 CAD cases and 2,636 controls remained. Then variant quality control was performed excluding variants (1) with more than 5% missing data, (2) with a Hardy Weinberg equilibrium P-value < 1 × 10^−6^, (3) with an allele frequency difference in control samples between data processing centres (BBJ and Nagahama cohort, Fisher’s exact test P < 1 × 10^−6^), (4) in the low complexity region, and (5) overlapping with insertions or deletions. After these procedures, we performed case control association analysis using genotypes in whole genome sequence data, and confirmed that the variant quality was well controlled (Supplementary Fig. 17). To construct the reference panel, we excluded singletons from the quality-controlled whole-genome sequencing data, and then haplotype phasing was performed using Eagle (v2.4.1)^43^. Phased VCFs were transformed into m3vcf format using minimac3 (v2.0.1)^44^. For comparison purposes, we obtained genotypes from the 1000 Genomes Project phase 3 (version 5). We then constructed the reference panel under the same pipeline for all subjects (n = 2,504) and for East Asian subjects (n = 504) separately.

### Haplotype phasing and imputation of case-control samples

The subjects included in the GWAS were genotyped using the HumanOmniExpressExome platform (Illumina) or in combination with HumanOmniExpress and HumanExomeBeadChip (Illumina). For variant quality control, we excluded variants with (1) call rates < 99%, (2) Hardy Weinberg Equilibrium *P*-values < 1.0 × 10^−6^, and (3) heterozygous counts less than five. After exclusion of these variants, we performed pre-phasing using Eagle. Phased haplotypes were imputed to the reference panels by minimac3^44^. For evaluation of the imputation quality, we created a phased genotype dataset comprised only of the OmniExpress13 array, imputed up to the 1KG_EAS_, 1KG_ALL_, and BBJ_CAD_ panels. After imputation, we compared the imputed dosage and genotypes directly determined by the exome array. Correlations were assessed by Pearson’s correlation coefficients. For all downstream analyses, we excluded variants with R^2^ < 0.3. Genotyped or imputed variants were annotated by ANNOVAR (Build 2017 Jul 7^45^), or the ClinVar database downloaded 4 February 2019.

### Phenotype

CAD was defined as a composite of stable angina, unstable angina, and myocardial infarction, which were determined by a physician upon study inclusion. The demographic features of the case-control cohort are provided in Supplementary Table 16. The quantitative trait data were obtained from medical records. Quantitative traits were normalized and adjusted as described below. Before normalization, we excluded samples from patients younger than 18 years and using the phenotype-specific criteria provided in Supplementary Table 17. We then corrected the effect of medication as follows. For individuals taking cholesterol-lowering drugs, TC and LDLC levels were corrected by dividing by 0.7 as previously reported^46,47^. For individuals taking antihypertensive drugs, 15 mmHg was added to the SBP and 10 mmHg to the diastolic blood pressure readings^48^. We next constructed a linear model for these phenotypes with sex, age, age^2^, the top 10 principal components and disease status. For WBC and C-reactive protein (CRP), smoking status was also introduced to the model. Using these models, we computed the residuals of the phenotype for each individual. The residuals were normalized by inverse-rank normalization and used as continuous variables^49^. Samples with excess heterozygosity, excess missing genotypes, a non-Japanese outlier identified by principal component analysis, and closely related samples estimated based on identity by states (PIHAT > 0.2) were excluded from the case-control or quantitative phenotype association studies.

### GWAS

The case-control association analysis was performed by logistic regression, implemented in PLINK2^50^. Sex, age, age^2^, and the top 10 principal components were included in the model as covariates. We tested 19,707,525 variants with a minor allele frequency (MAF) ≥ 0.02% in case-control population, because our reference panel contained almost ten thousand haplotypes and variants with minor allele counts ≥ 2 (0.02%). The genome-wide significance threshold was set at P < 5 × 10^−8^ for variants with MAF ≥ 1%, and *P* < 3.93 × 10^−9^ (0.05/12,710,563) for those with MAF < 1% (number of variants with MAF < 1% = 12,710,563). To define a locus, we created a set of genomic ranges adding 500 kb bilaterally for all variants with genome-wide significance, and then merged overlapping ranges. In the MHC region (chromosome 6: 25,000,000–35,000,000 bp), we added 1 Mb to the signal bilaterally. Previously reported loci were also created in the same manner based on the curated top variants. Significant loci without overlap and a lead variant not in LD (<0.10) in both the East Asian or European population with previously reported loci were considered novel. LD was estimated using genotypes of the 1KG dataset. For quantitative traits, we performed linear regression analysis implemented in PLINK2 for normalized phenotypes as described above.

### LD score regression and heritability estimation

We performed LD score regression with ldsc software to estimate bias from population stratification and explained heritability^24^. We used the 1KG EAS population as the reference LD panel and included only variants present in HapMap3 SNPs. For the calculation of liability scale heritability, we assumed that the CAD prevalence in the Japanese population is 0.8% based on a Japanese government report published in 2009 (https://www.mhlw.go.jp/).

### Stepwise conditional analysis

To identify statistically independent signals in the loci, we performed sample-level stepwise conditional analysis for each genome-wide significant locus defined as denoted above. In the first step, we added the dosage of the lead variant to the covariates and performed logistic regression for all variants in the locus. If we found a locus-wide significant association (P < 1 × 10^−5^), we additionally introduced the dosage of the most significant variant to the covariates. This procedure was repeated until none of the variants showed locus-wide significance.

### Meta-analysis

We obtained summary statistics for the European CAD-GWAS from the website of the CARDIoGRAM plus C4D consortiums (http://www.cardiogramplusc4d.org/data-downloads/)^7^ and online supplementation of a previous report (https://data.mendeley.com/datasets/2zdd47c94h/1)^9^. We aligned the β and allele frequency to the alternate allele of hg19 and then merged only SNPs with an MAF ≥ 1% in all the summary statistics. The transethnic meta-analysis was performed using MANTRA(v2)^30^ and genome-wide significance was set at log_10_ BF > 6 according to a previous simulation result^31^. We excluded log_10_ BF for heterogeneity > 6. For PRS derivation, β and P-values were calculated by fixed- and random-effects meta-analyses using METASOFT (v2)^51^.

### Estimation of allelic concordance

To obtain LD-independent variant sets at various significance levels, we performed *P*-value thresholding for each *P*-value threshold from the summary statistics obtained from the fixed-effect transethnic meta-analysis and random-effect transethnic meta-analyses. For these variant sets, we calculated the Pearson’s correlation coefficient of the β between the European study and Japanese studies. The 1KG European population was used as the LD reference and the LD threshold was fixed at 0.8.

### Credible set analysis

To construct sets of variants that likely include causal variants in each significant locus identified by the transethnic meta-analysis, we performed a credible set analysis. For each genome-wide associated locus, we calculated the PPA (π)^52^ for all variants as follows:

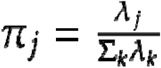

Here, λ denotes the BF for variant j, and k denotes all of the variants included in the locus. We listed the variants in order of decreasing PPA, and then constructed the 99% credible set including variants from the top PPA until the cumulative PPA reached 0.99. For comparison, we performed the European meta-analysis using the C4D and the UKBB datasets (referred to as the European analysis).

### PRS

We derived the PRS by the *P*-value thresholding (P/T) method using summary statistics from the 1) the Japanese GWAS with MAF > 0.01, 2) the transethnic meta-analysis of the fixed-effect model, and 3) the mixed-effect model. We used the P-value thresholds 1.0, 0.5, 0.1, 0.05, 0.01, 1× 10^−3^, 5 × 10^−4^, 5 × 10^−6^, 5 × 10^−8^, and R^2^ thresholds 0.2, 0.4, 0.6, and 0.8. Pruning and thresholding procedure was performed by PLINK software version 1.90b3.37. Individual risk scores were computed with derived weights in independent case-control cohorts (case = 1,827, control = 9,172) by multiplying the genotype dosage and corresponding weight using PLINK2 software. To assess the performance of the PRS, we calculated the Nagelkerke’s R^2^, the AUC values and OR using a logistic regression model adjusted by age and sex. OR indicates the ratio of disease prevalence in the top decile to that in the bottom decile. For a comparison of AUC values, we performed DeLong’s test implemented in the pROC package for R. The previously derived PRS from the European study^16^ was downloaded from Cardiovascular Disease Knowledge Portal (http://www.broadcvdi.org).

### Survival analysis

For survival analysis, we obtained survival follow-up data with the cause of death under the ICD-10 code for 132,737 individuals from the BBJ project. Data collection and feasibility were reported previously^40,41^. Briefly, survival status was collected from medical records or resident card. Then we obtained vital statistics from the Statistics and Information Department of Ministry of Health, Labor and Welfare in Japan, and identified the cause of death according to ICD-10. The follow-up rate was 97% and the median follow-up period was 7.7 years. We divided the cause of death based on ICD-10 classifications and excluded categories with fewer than 100 events. HRs and associated P-values were calculated for genotype dosage or PRS by the proportional hazard model adjusting for sex, age, age^2^, top 10 principal components, and disease status. Analyses were performed with the R package survival, and survival curves were estimated using the R package survminer with modification.

### PRS associated phenotypes

For clinical evaluation of the CAD-PRS, we randomly split and withheld 1/3 of the control samples. Using the remaining control samples and all case samples, we re-performed the Japanese GWAS, transethnic meta-analysis, and PRS derivation (Supplementary Fig. 1c). Using the derived model, we calculated the PRS for the withheld control samples, and assessed relationships between the CAD-PRS and clinical indices including 32 numerical clinical traits and two binary lifestyle traits (i.e., cigarette smoking and alcohol drinking). For the numerical traits, we calculated Spearman’s correlation coefficients, associated confidence intervals, and P-values. For the binomial traits, we performed logistic regression adjusted by sex, age, age^2^, top ten principal components, and disease status. We set the significance threshold at P < 0.05/34.

### Data availability

Summary statistics of the Japanese GWAS will be publically available in National Bioscience Database Center (research ID hum0014, https://humandbs.biosciencedbc.jp/).

## Acknowledgments

We appreciate the staff of BBJ for their excellent assistance in collecting samples and clinical information. We also thank the Nagahama study, the Japan Public Health Centre-based prospective Study (JPHC) study and Japan Multi-Institutional Collaborative Cohort (J-MICC) Study, and Osaka Acute Coronary Insufficiency Study (OACIS) study for their invaluable contributions to the study. We are grateful to the CARDIoGRAMplusC4D investigators, P.v.d Harst, and N Verweij for making their data publically available. We thank A.P. Morris for providing us with MANTRA software with valuable advice.

## Author Contributions

S.K., K.I., C.T., M.K., and Y.K. conceived and designed the study. C.K., J.S., K.H., and F.M. collected, managed and genotyped the Nagahama cohort. K.M, Y. Murakami, and M.K. collected and managed the BBJ sample. M.I, T.Y, N.S, and S.T collected and managed the JPHC study. T.K., H. Ikezaki, N.T., K.T., K.A., K.K., M.N., and K.W. collected and managed the J-MICC study. S.S., Yasuhiko Sakata, H.S., M. Hori, Yasushi Sakata collected and managed the OACIS study. C.T., Y. Momozawa, A.T., M.K., and Y.K., performed genotyping. S.K., K.I., C.T., and Y.K. performed statistical analysis. S.K., K.I., C.T., M.A., M. Horikoshi, H. Matsunaga, H. Ieki., K.O., Y.O. contributed to data processing, analysis and interpretation. S.N., H. Morita, H. Akazawa., H. Aburatani, and I.K. supervised the study. S.K. and K.I. wrote the manuscript and many authors also provided valuable edits.

