## Supplementary Figures for "Population-specific and transethnic genome-wide analyses reveal distinct and shared genetic risks of coronary artery disease"

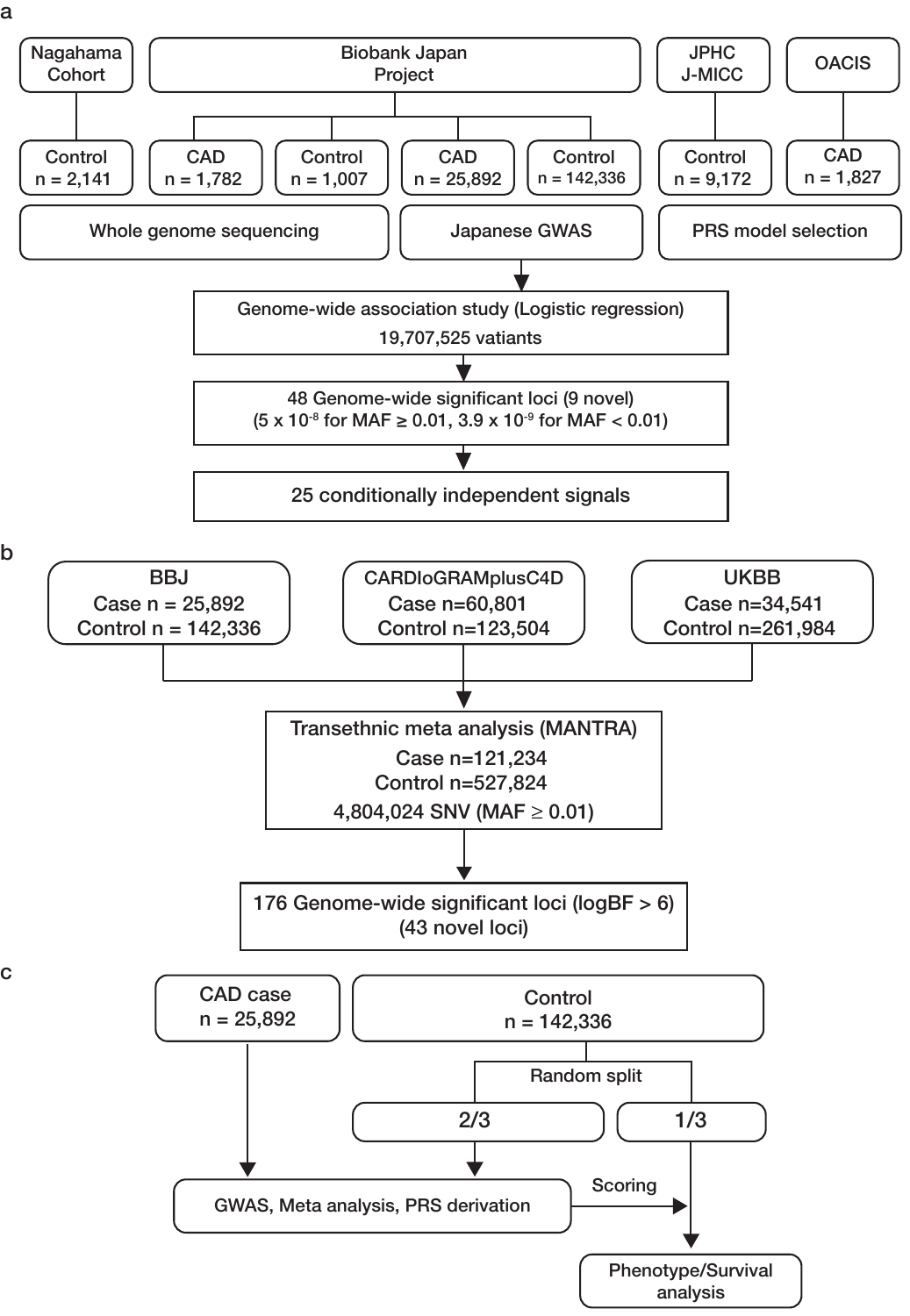

**Supplementary Fig. 1 | Overview of the study. a**, Overview of whole-genome sequencing, Japanese GWAS, and PRS model selection. **b**, Overview of transethnic meta-analysis. **c**, Overview of PRS phenotype analysis JPHC, Japan Public Health Center-based prospective Study; J-MICC, Japan Multi-Institutional Collaborative Cohort; OACIS, Osaka Acute Coronary Insufficiency Study; UKBB, UK Biobank; CAD, coronary artery disease; MAF, minor allele frequency; GWAS, genome-wide association study; PRS, polygenic risk score; SNV, single nucleotide variant; BF Bayes factor.

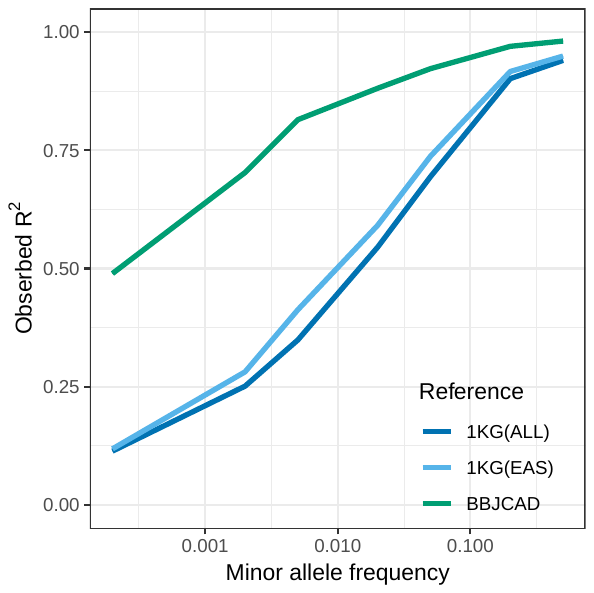

**Supplementary Fig. 2 | Imputation accuracy of BBJCAD panel and 1KG panels.** The mean-observed R^2^ for each MAF bin is plotted. Observed R^2^ indicates Fisher’s correlation coefficient between imputed dosage and genotypes determined by the genotyping array. 1KG, 1000 Genomes Project; EAS, East Asian; BBJ, Biobank Japan; CAD, coronary artery disease.

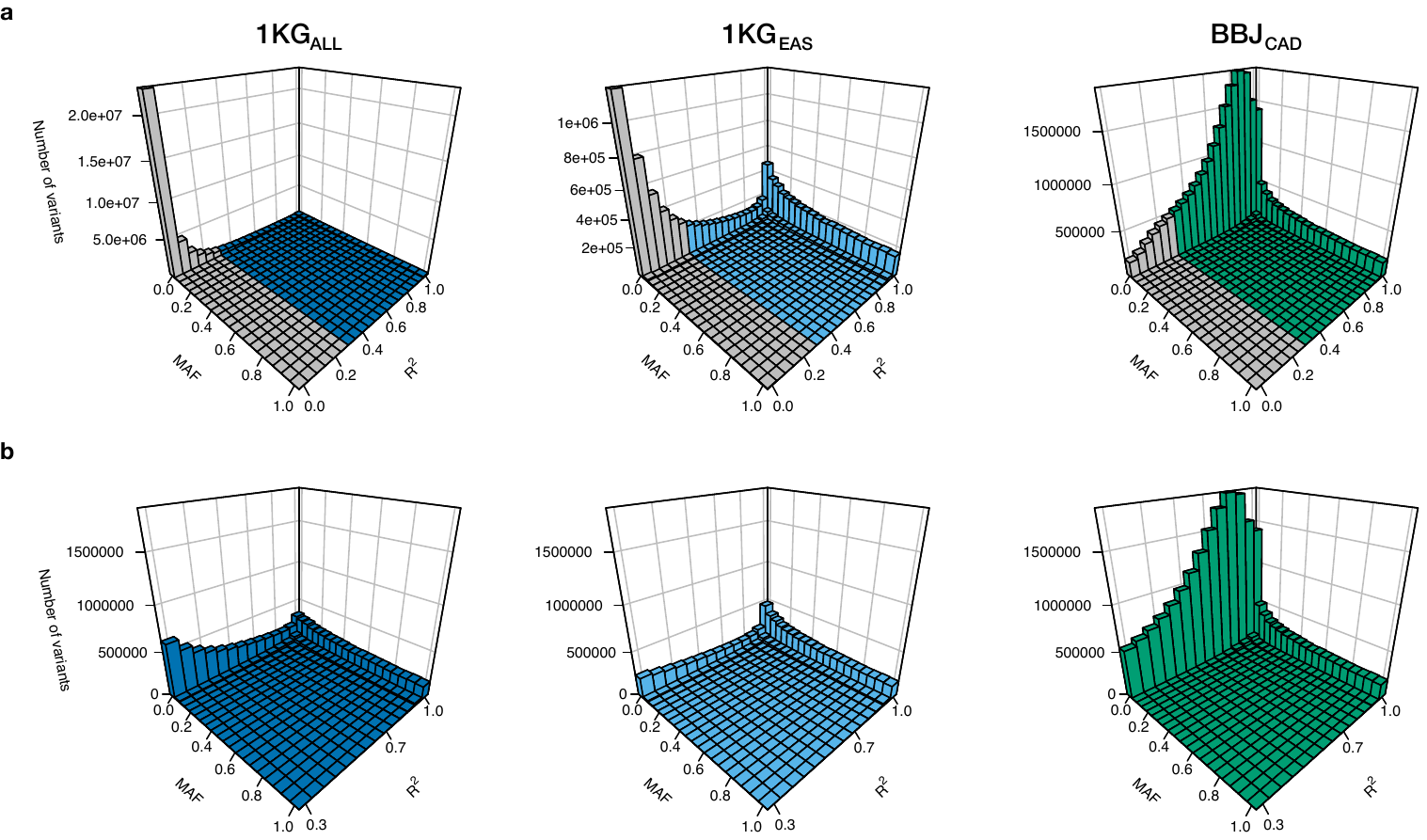

**Supplementary Fig. 3 | Spectra of minor allele frequency and imputation quality.** **a**, The distributions of all imputed variants stratified by MAF and R^2^. Z-axis indicates the number of variants **b**, The distributions of variants with imputation quality ≥ 0.3. Note that the Z axes in **a**, have different scales. MAF; minor allele frequency. 1KG, 1000 Genomes Project; EAS, East Asian; BBJ, Biobank Japan; CAD, coronary artery disease.

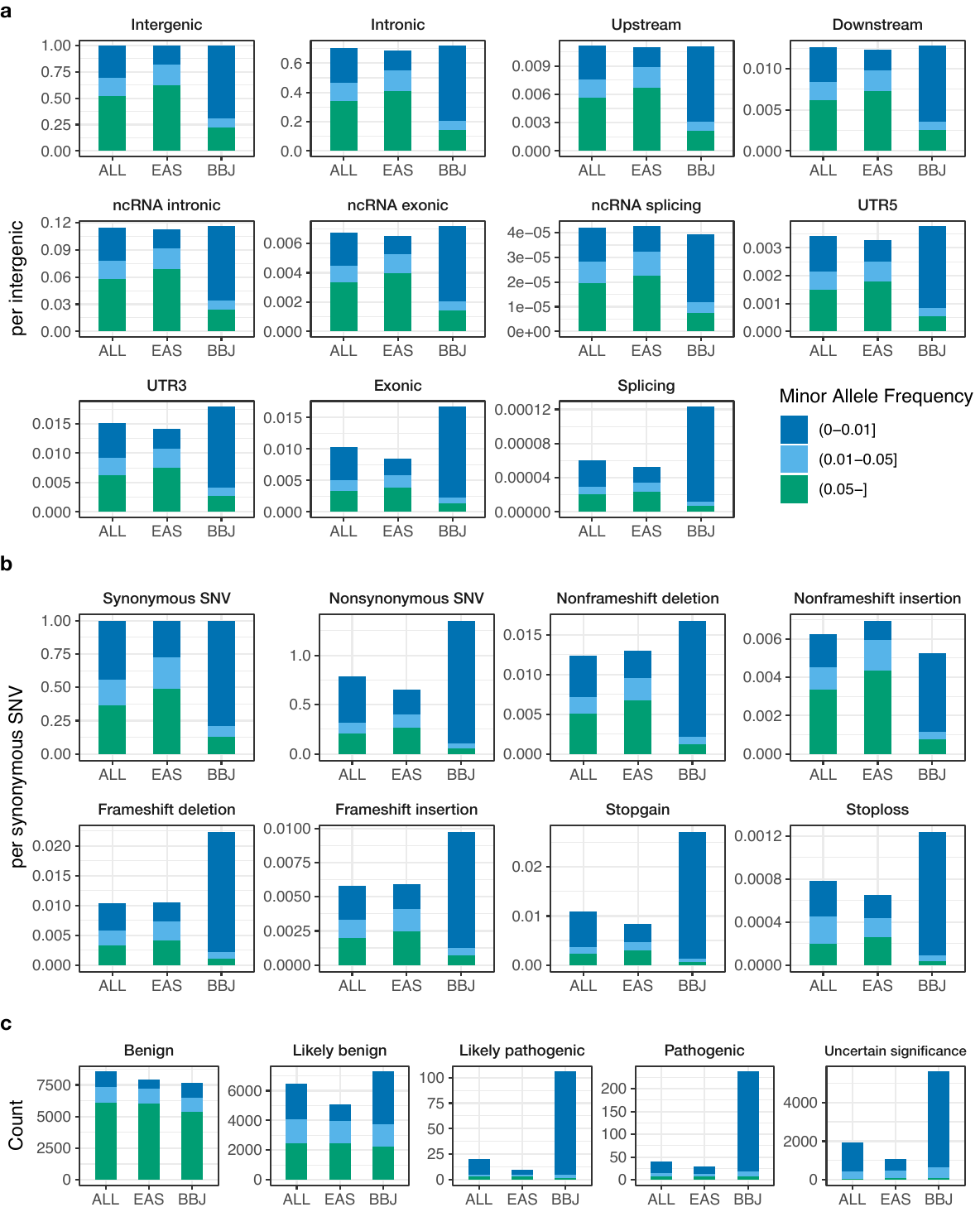

**Supplementary Fig. 4 | Distributions of the number of imputed variants across functional classes.** **a**, The number of imputed (R^2^ ≥ 0.3) variants in various functional classes. X-axis indicates the reference population. Y-axis indicates the ratio of the number of variants in each class to that of intergenic variants. The color indicates the proportion of the variants in minor allele frequency bins. **b**, The number of imputed (R^2^ ≥ 0.3) exonic variants. The ratios of the number of variants in each class to those of nonsynonymous variants are shown. **c**, The number of imputed variants registered in ClinVar database. 1KG, 1000 Genomes Project; EAS, East Asian; BBJ, Biobank Japan; CAD, coronary artery disease. ncRNA, non-coding RNA; UTR, untranslated region; SNV, single nucleotide variant.

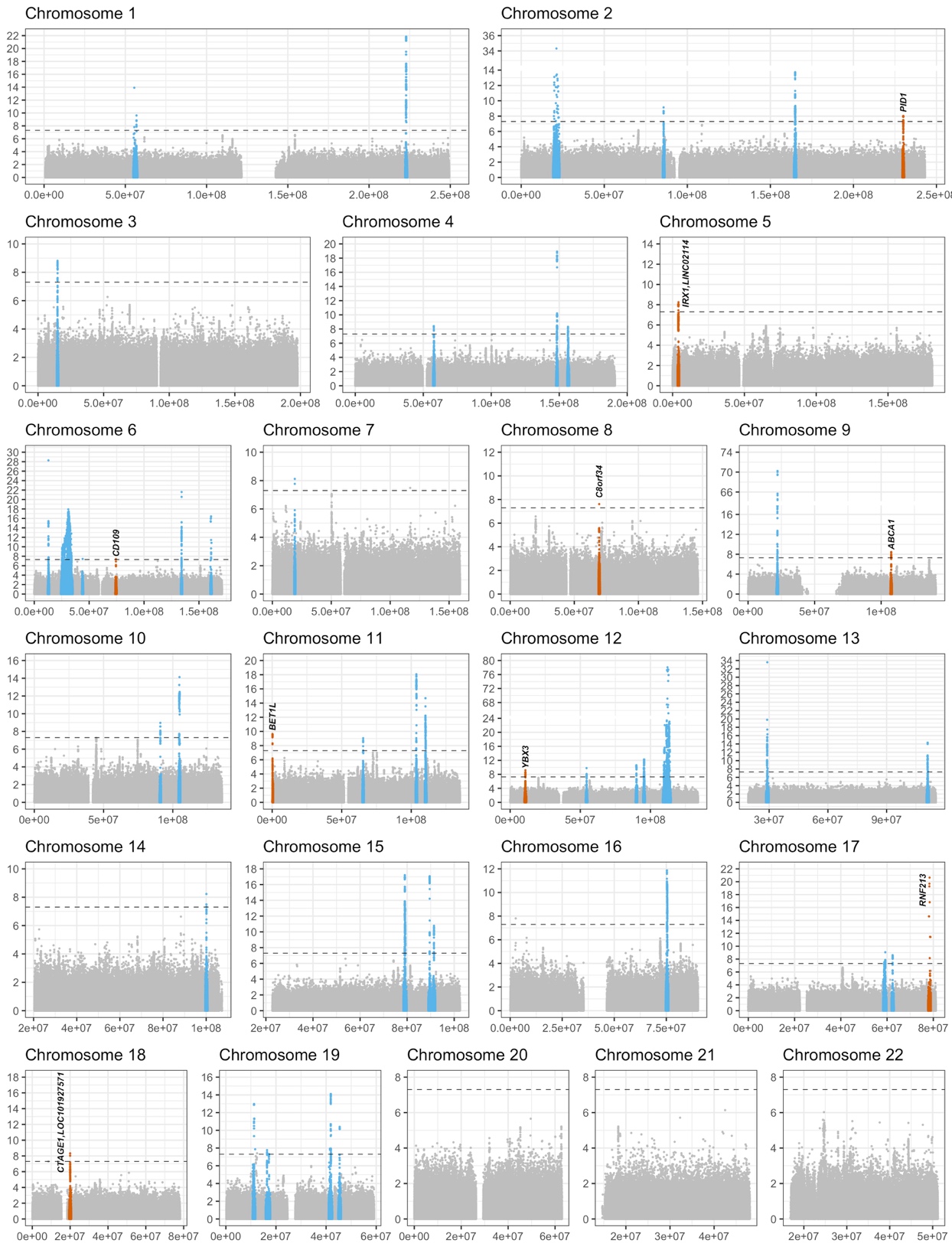

**Supplementary Fig. 5 | Manhattan plots for Japanese GWAS.** The results of the case-control association study of Japanese GWAS (CAD case 25,892, control 142,336) are shown. The negative log_10_ *P*-values on the y-axes are shown against the genomic position (hg19) on the x-axes. Variants in 9 novel and 39 previously reported loci are presented in orange and blue, respectively. Dashed lines indicate genome-wide significant thresholds (*P* = 5 × 10^-8^).

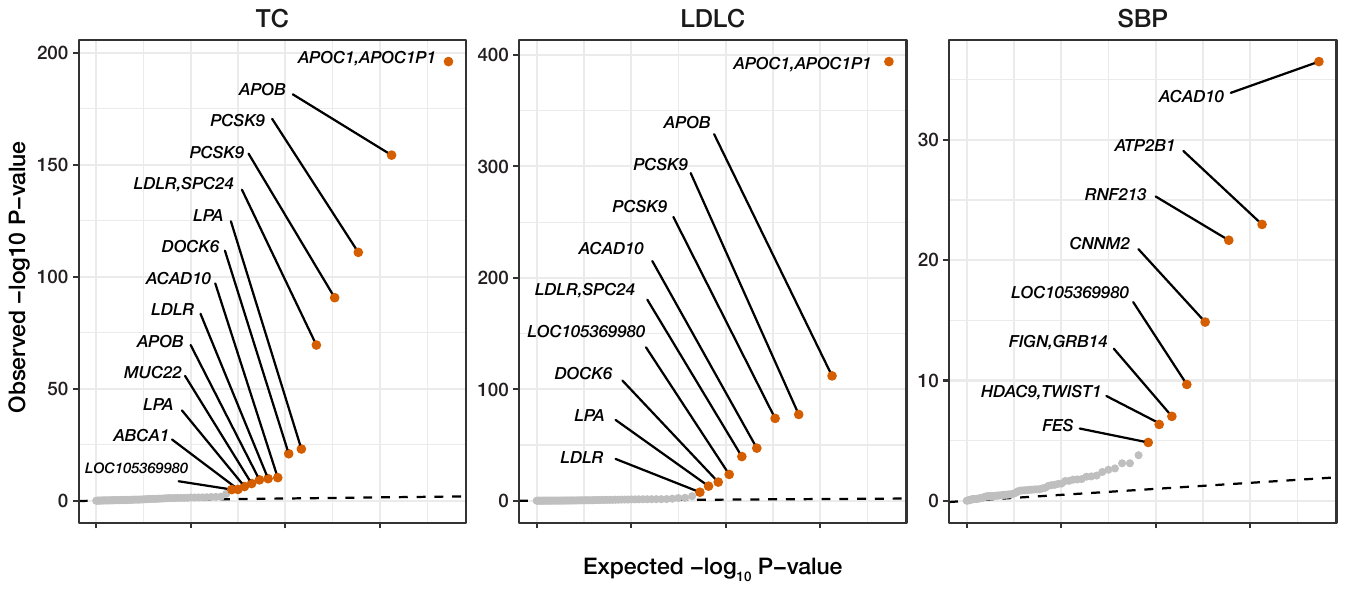

**Supplementary Fig. 6 | The pleiotropic effects of distinct signals.** Quantile-quantile plots for association analysis of 73 independent signals determined by association study for TC (n = 134,314), LDLC (n = 76,477), and SBP (n = 139,847) in the BBJ dataset. Variants showing significant associations (P < 0.05 / 2,482) for the quantitative traits are plotted in orange. Dashed lines indicate theoretical distributions. TC, total cholesterol; LDLC, low-density lipoprotein cholesterol; SBP, systolic blood pressure.

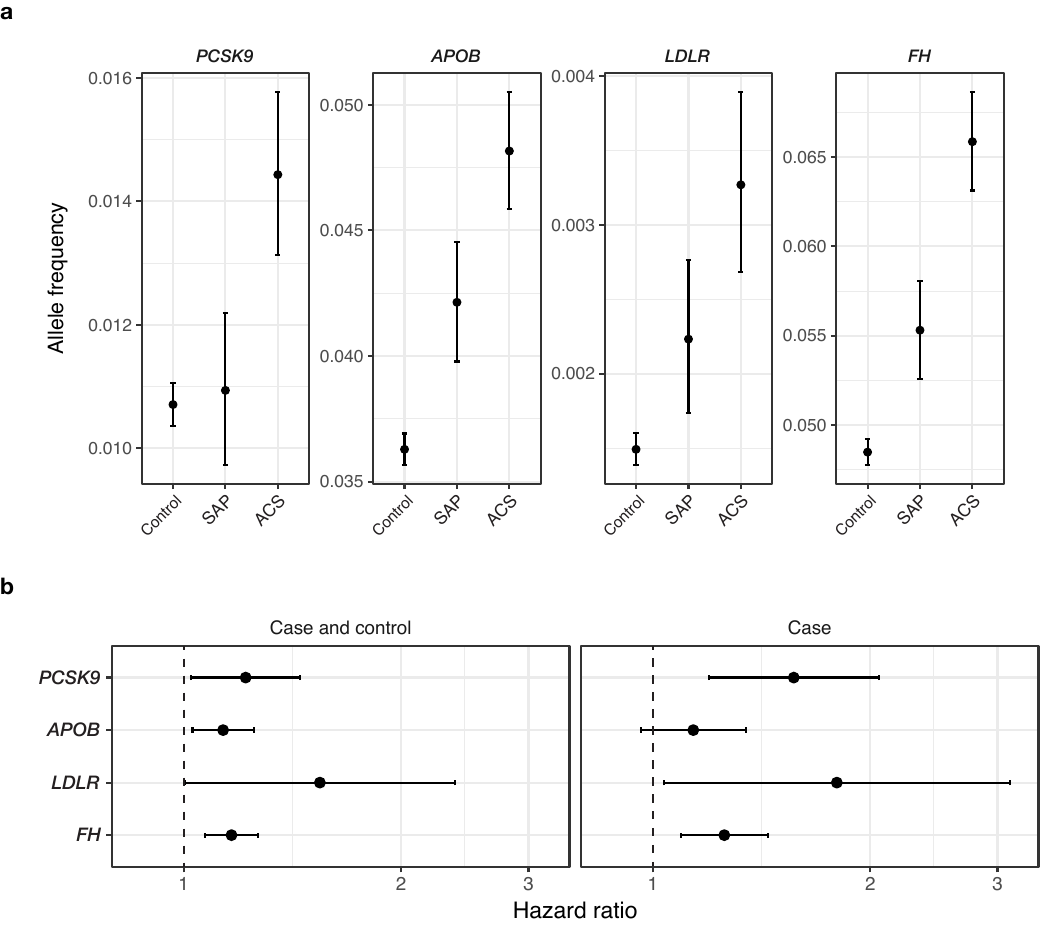

**Supplementary Fig. 7 | Summary of survival analysis and allele frequency in disease subtypes.** **a**, Each point indicates allele frequency stratified by disease status. Error bars represent 95% confidence intervals, which were estimated by 10^5^ times bootstrapping. **b**, Each point indicates the hazard ratio per allele, and each error bar indicates its 95% confidence interval. SAP, stable angina pectoris; ACS, acute coronary syndrome; FH, familial hypercholesterolemia.

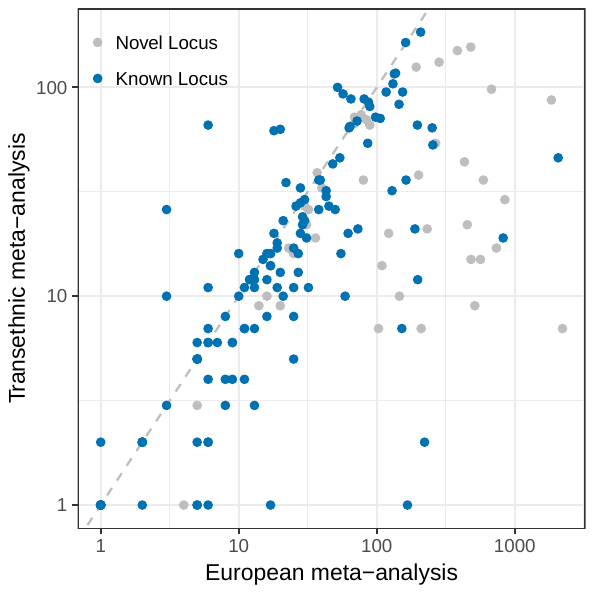

**Supplementary Fig. 8 | Distribution of the sizes of the 99% credible sets.** Each dot indicates the number of variants in the 99% credible set for 176 genome-wide associated loci in the European only meta-analysis (C4D and UKBB, x-axis) and transethnic meta-analysis (BBJ, C4D, and UKBB, y-axis). Previously established loci (n = 133) are plotted in blue and novel loci (n = 43) are plotted in grey. BBJ, Biobank Japan; C4D, Coronary ARtery DIsease Genome wide Replication and Meta-analysis plus The Coronary Artery Disease Genetics; UKBB, UK Biobank.

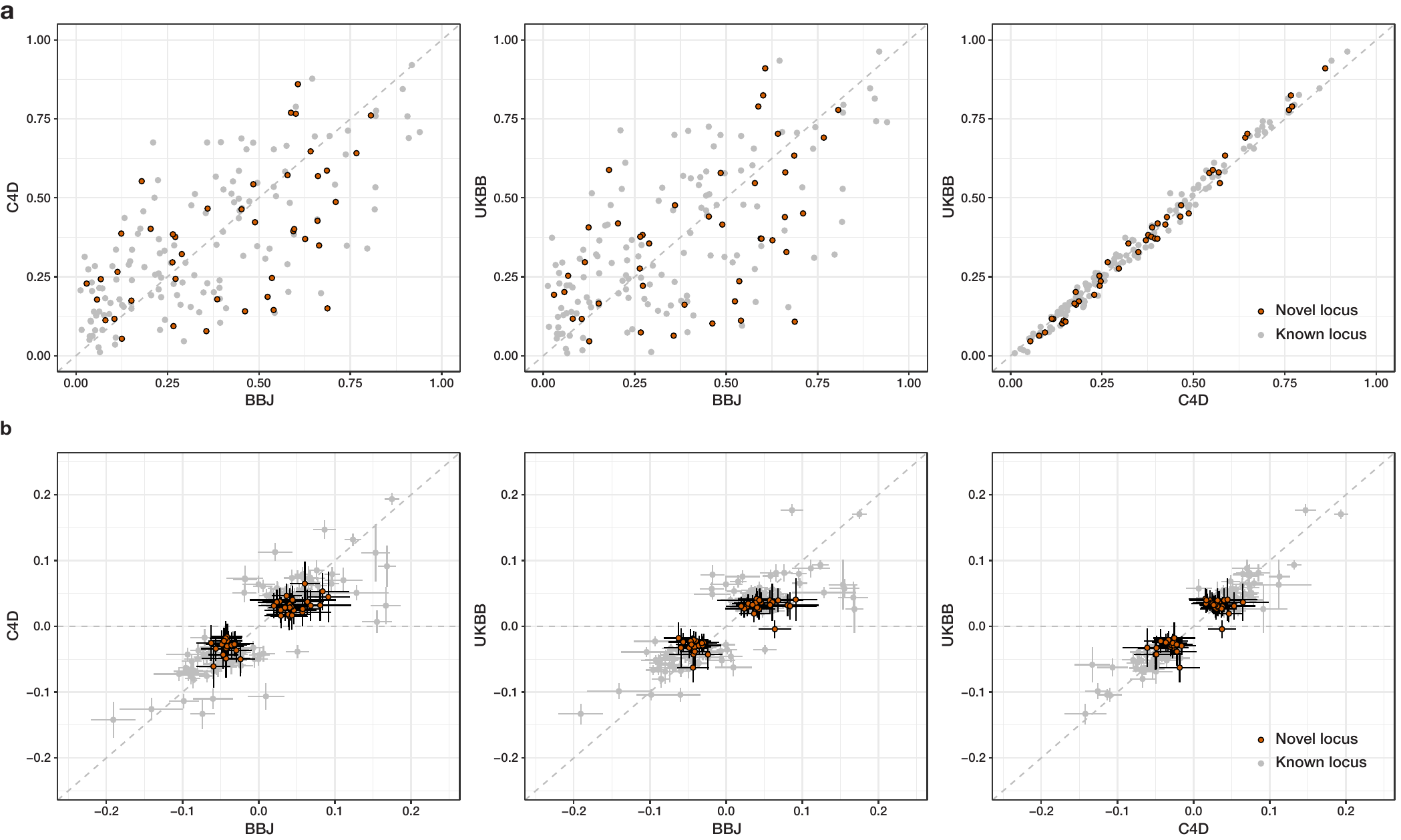

**Supplementary Fig. 9 | Transethnic comparison of allele frequencies and allelic effects.** **a**, AAF, and **b**, the estimated effect sizes of the 176 lead variants in each study are shown. Alternate alleles were aligned to the reference genome (hg19). Each error bars in **b**, indicates the 95% confidence interval of the beta estimate in each study. Grey dots indicate previously reported loci and orange dots indicate newly identified loci in this study. AAF, alternate allele frequency; BBJ, Biobank Japan; C4D, Coronary ARtery DIsease Genome wide Replication and Meta-analysis plus The Coronary Artery Disease Genetics; UKBB, UK Biobank.

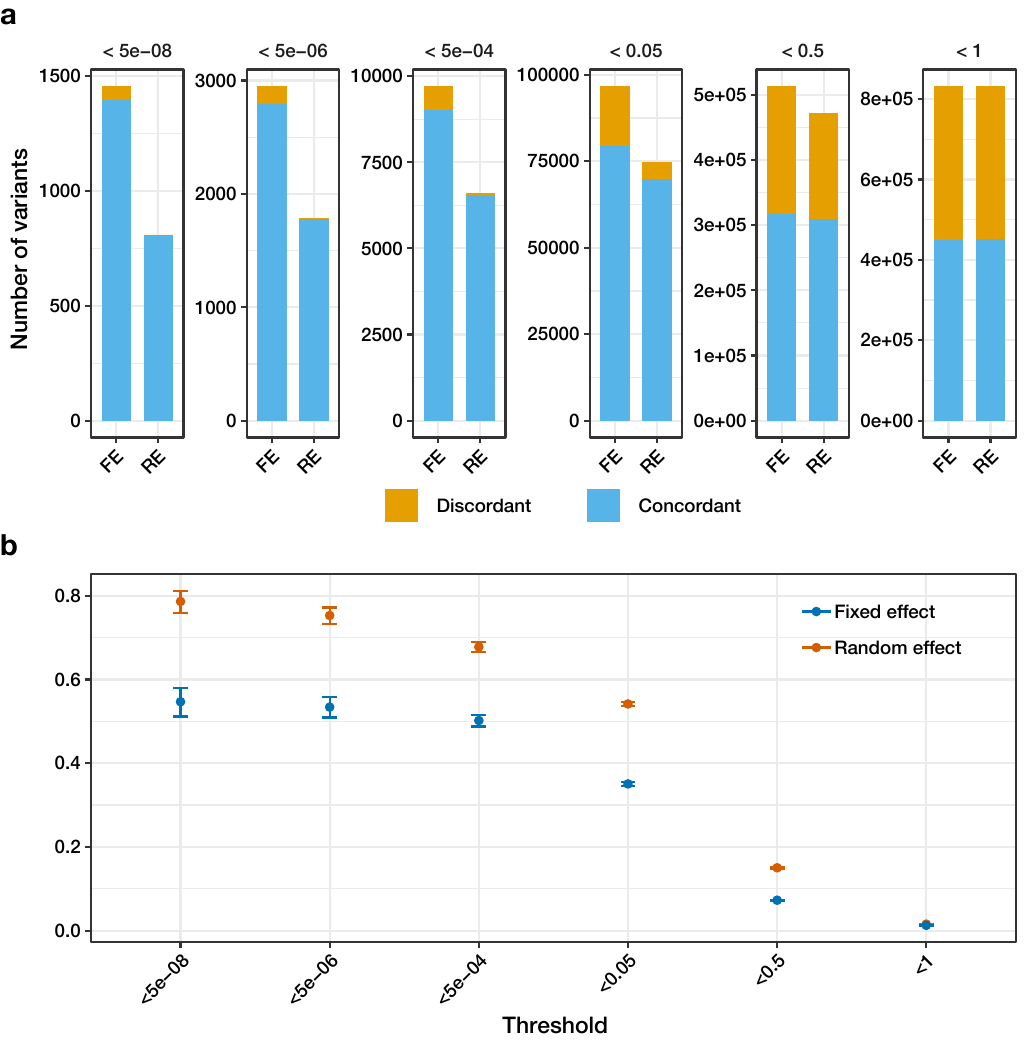

**Supplementary Fig. 10 | Consistency of allelic effect.** **a**, Allelic effect concordance of LD-independent variants between Japanese and European studies. LD-independent variants were determined by *P*-value thresholding using the fixed-effect meta-analysis or random-effect meta-analysis result with a fixed R^2^ threshold (0.8). **b**, Each point indicates the correlation coefficient of beta values of the variant set determined as in **a**. LD, Linkage disequilibrium; FE fixed-effect; RE random-effect.

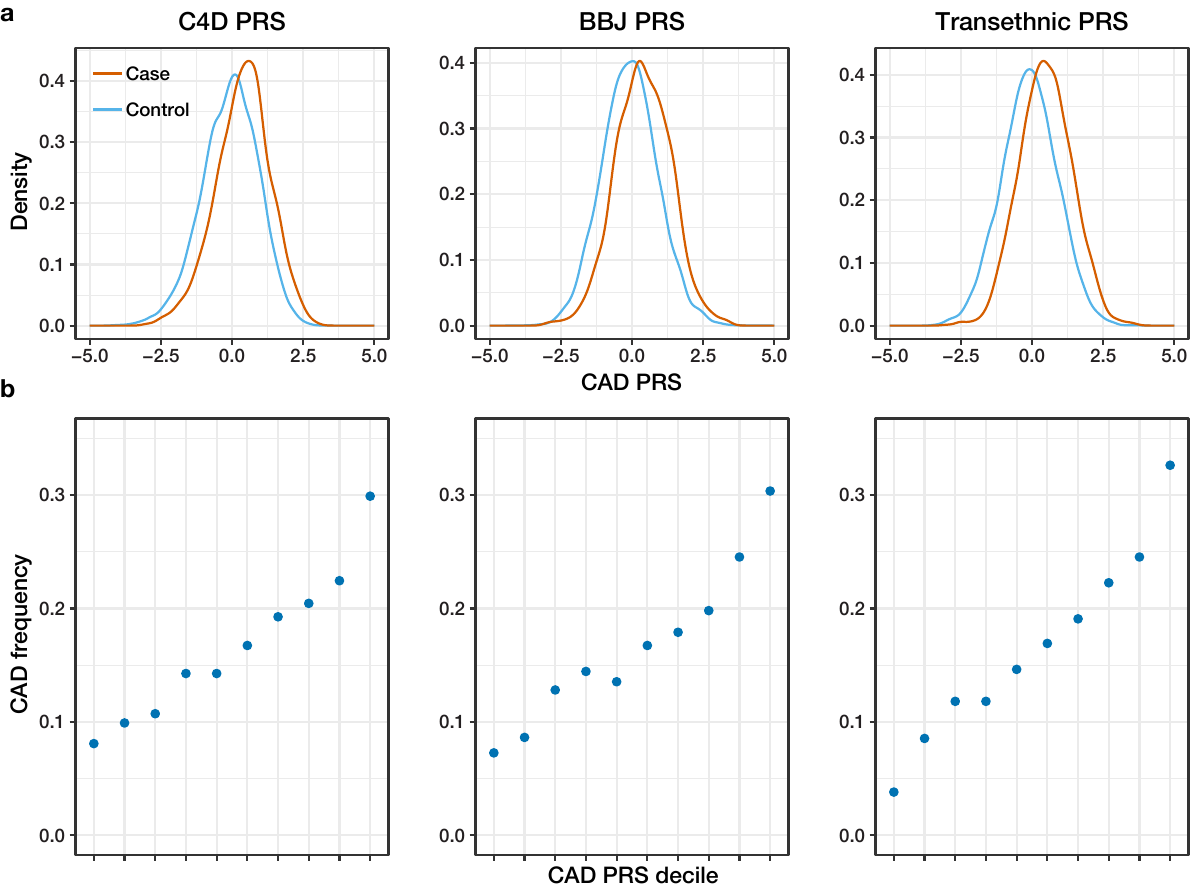

**Supplementary Fig. 11 | Performances of CAD-PRS derived from European study, Japanese study, and transethnic meta-analysis.** **a**, The distribution of European (C4D)-, Japanese (BBJ)- and transethnic meta-analysis-derived PRS in a Japanese validation cohort (1,827 cases, 9,172 controls) are shown from left to right. X-axis indicates normalized CAD-PRS. **b**, CAD prevalence in each CAD-PRS decile. C4D PRS: Previously published weights (Khera et al. Nat Genet 2018) were used for PRS calculation. BBJ PRS: PRS weights were derived from current Japanese GWAS by the *P*-value threshold method. Transethnic PRS: PRS weights were derived from transethnic meta-analysis summary statistics (random effect). PRS, polygenic risk score; CAD, coronary artery disease; BBJ, Biobank Japan; C4D, Coronary ARtery DIsease Genome wide Replication and Meta-analysis plus The Coronary Artery Disease Genetics; GWAS, genome wide association study.

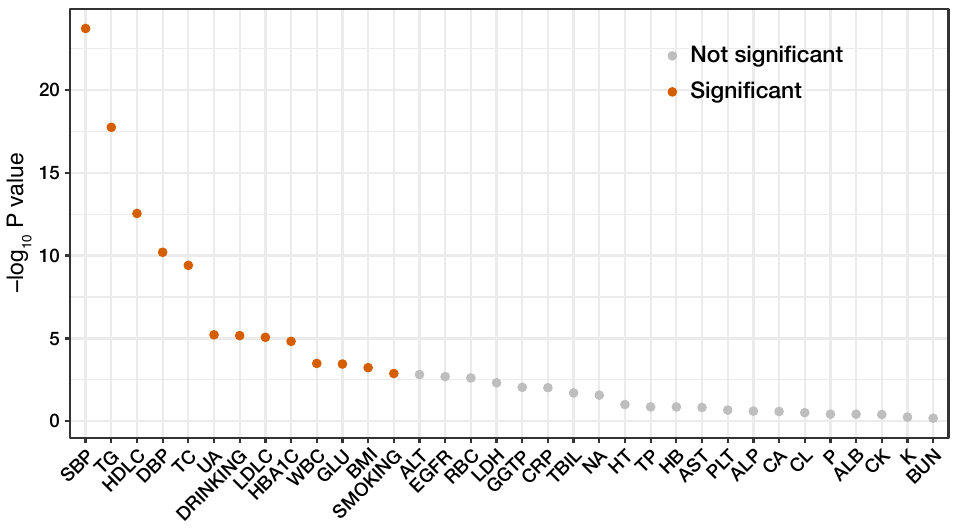

**Supplementary Fig. 12 | Significant associations between clinical traits and CAD-PRS.** Negative log_10_ *P*-values of Spearman’s correlation coefficient or negative log_10_ P-values that of beta coefficient estimated by logistic regression (for cigarette smoking and alcohol drinking behaviour) are presented. Orange dots indicate traits with Bonferroni adjusted significance (P = 0.05/34). Abbreviations are defined in Supplementary Table 17.

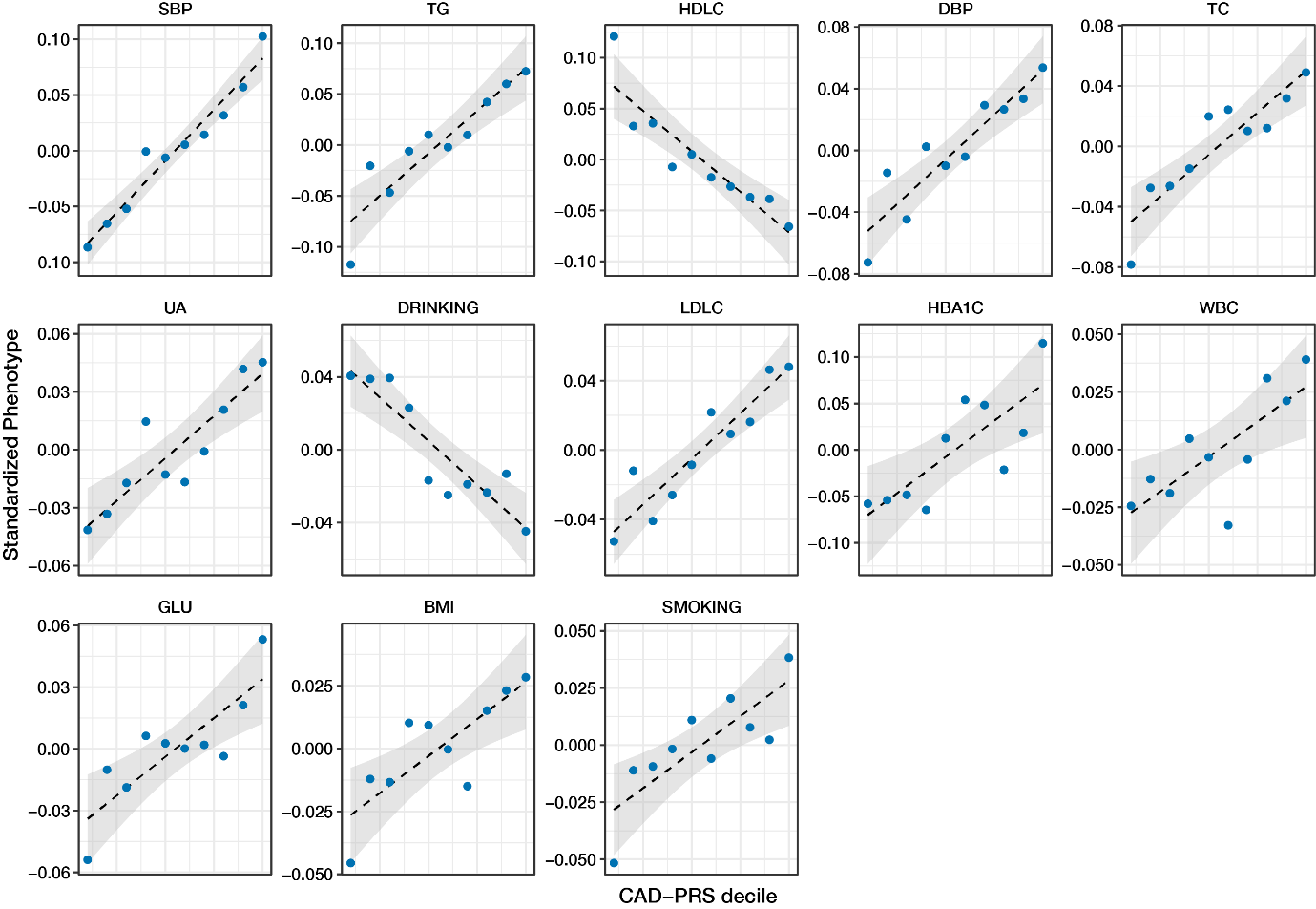

**Supplementary Fig. 13 | Dose relationships between CAD-PRS and associated traits.** Each point represents the mean value of standardized phenotypes corresponding to CAD-PRS decile. Regression lines and 95% confidence intervals are shown in dashed lines and grey areas, respectively. SBP, systolic blood pressure; TG, triglyceride; HDLC, high-density lipoprotein cholesterol; DBP diastolic blood pressure; TC, total cholesterol; UA, uric acid; LDLC, low-density lipoprotein cholesterol; HBA1C, haemoglobin A1c; WBC, white blood cell count; GLU, glucose; BMI, body mass index; CAD, coronary artery disease; PRS, polygenic risk score.

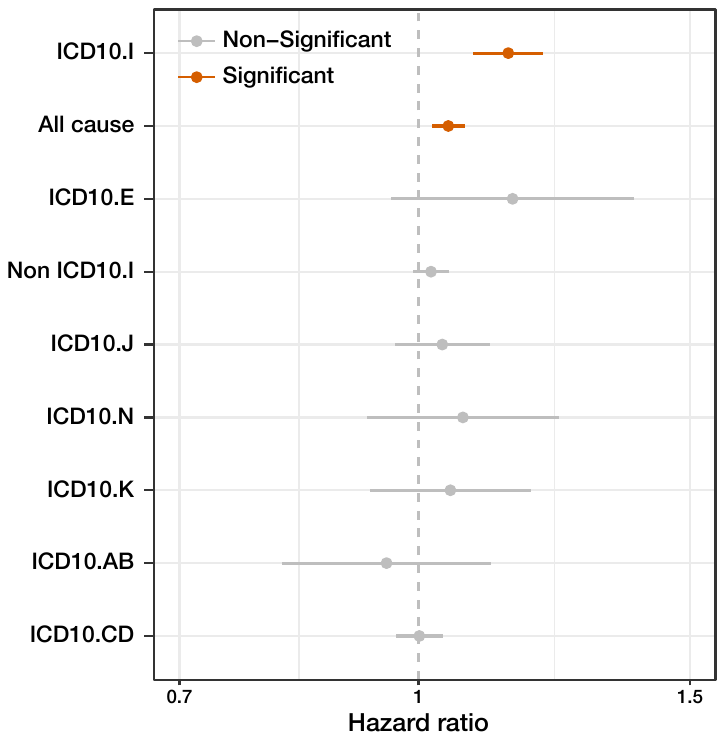

**Supplementary Fig. 14 | Summary of survival analysis.** Each dot indicates the estimated hazard ratio of per-1SD increase in CAD-PRS for each ICD10 category. Error bars indicates a 95% confidence interval. Bonferroni adjusted significant associations (P < 0.05/9) are plotted in orange. ICD10, International Statistical Classification of Diseases and Related Health Problems 10th Revision; ICD10.I, Diseases of the circulatory system; ICD10.E, Endocrine, nutritional and metabolic diseases; ICD10.J, Diseases of the respiratory system; ICD10.N, Diseases of the genitourinary system; ICD10.K, Diseases of the digestive system; ICD10.AB, Certain infectious and parasitic diseases; ICD10.CD Neoplasms.

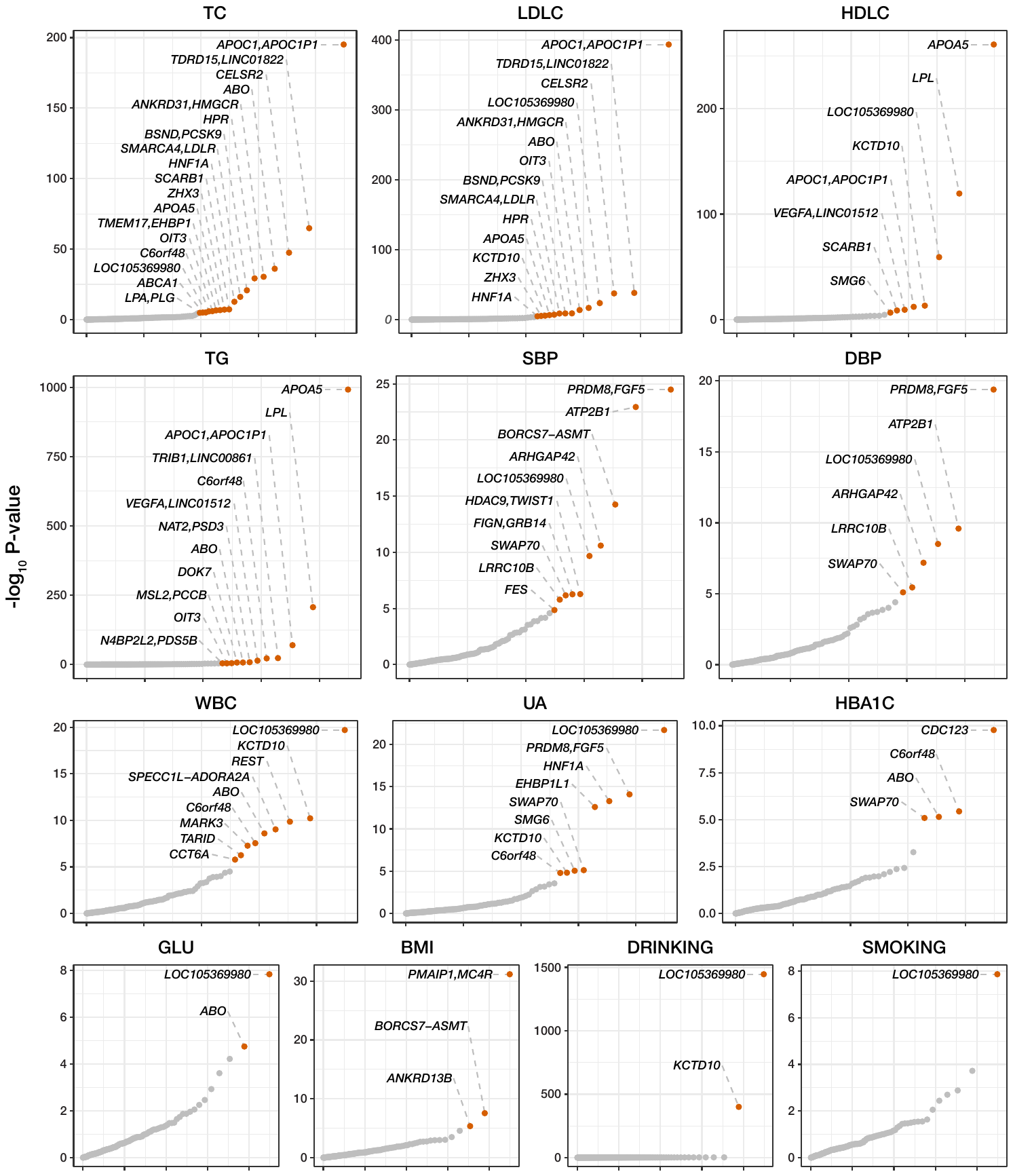

**Supplementary Fig. 15 | Pleiotropy of lead variants detected in transethnic meta-analysis.** Quantile-quantile plots of the association analysis of 176 lead variants determined by transethnic meta-analysis for 13 CAD-PRS associated traits. Y-axis indicates the negative log_10_ *P*-value. Associations with Bonferroni adjusted significance (P < 0.05/2,262) are plotted in orange. TC, total cholesterol; LDLC, low-density lipoprotein cholesterol; HDLC, high-density lipoprotein cholesterol; TG, triglyceride; SBP, systolic blood pressure; DBP, diastolic blood pressure; WBC, white blood cell count; UA, uric acid; HBA1C, haemoglobin A1c, GLU, glucose; BMI, body mass index.

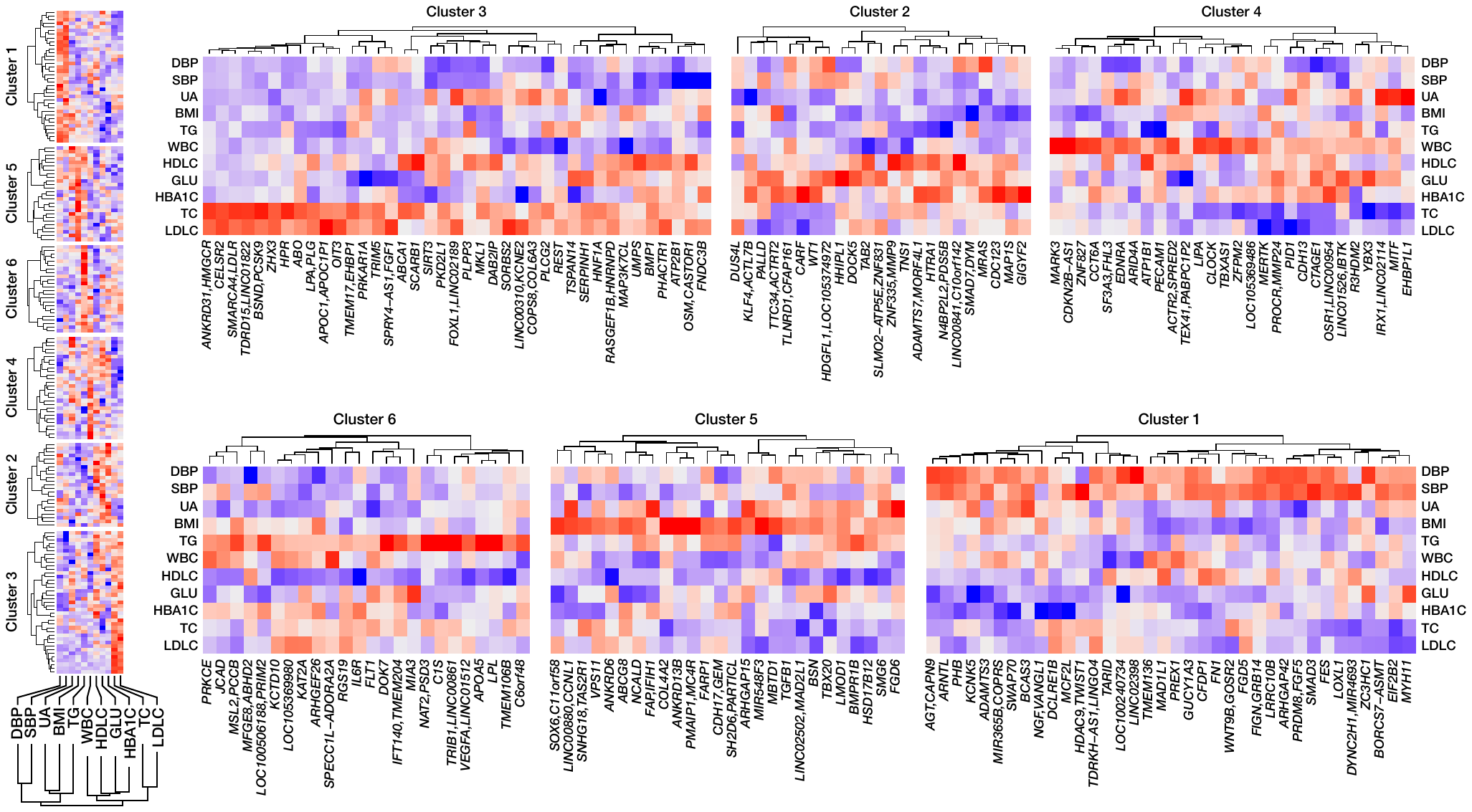

**Supplementary Fig. 16 | Unsupervised hierarchical clustering of the lead variants detected in transethnic meta-analysis.** The matrices of Z-scores of 176 lead variants for 11 CAD-PRS associated quantitative traits are presented. Red colour indicates that the CAD risk allele is associated with increased value of the phenotype. Blue colour indicates that the CAD risk allele is associated with decreased value of the phenotype. Loci are divided into 6 clusters by k-means clustering and reordered according to hierarchical clustering. DBP, diastolic blood pressure; SBP, systolic blood pressure; UA, uric acid; BMI, body mass index; TG, triglyceride; WBC, white blood cell count; HDLC, high-density lipoprotein cholesterol; GLU, glucose; HBA1C, haemoglobin A1c; TC, total cholesterol; LDLC, Low-density lipoprotein cholesterol.

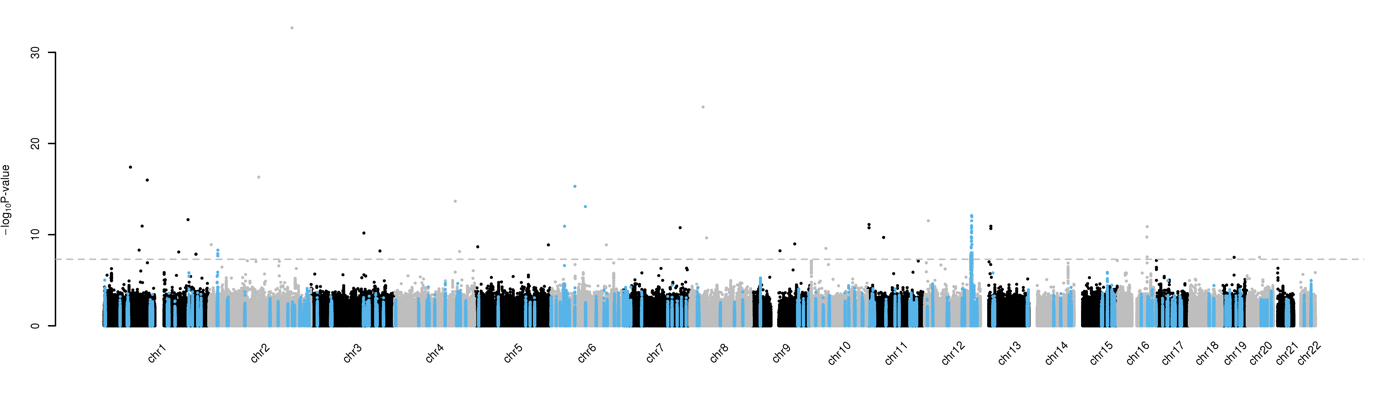

**Supplementary Fig. 17 | Case control association study using whole genome sequence data.** The results of case control association study including 1,781 CAD cases and 2,636 controls are shown. The negative log_10_ *P*-values are shown against the genomic coordinates (hg19). Variants in 167 previously reported loci are presented in blue. Dashed line indicates genome-wide significant threshold (*P* = 5 × 10^-8^).
