## Supplementary Dataset for "Population-specific and transethnic genome-wide analyses reveal distinct and shared genetic risks of coronary artery disease"

### Supplementary Data Set

1. Newly identified 9 loci in Japanese GWAS
2. Previously reported 39 loci in Japanese GWAS
3. Newly identified 43 loci in Meta analysis
4. Previously reported 133 loci in Meta analysis

### 1. Newly identified 9 loci in Japanese GWAS

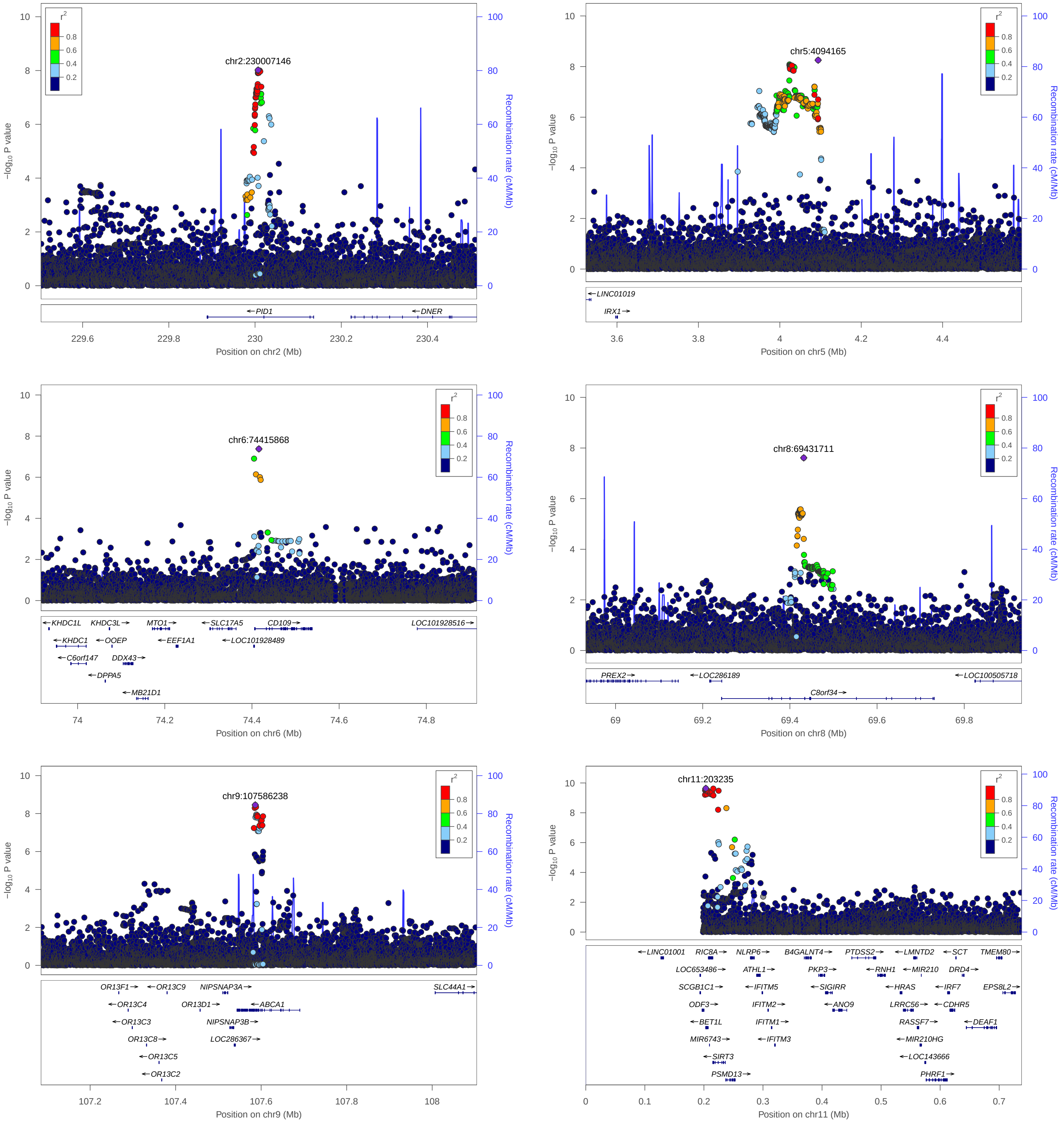

### 1. Newly identified 9 loci in Japanese GWAS

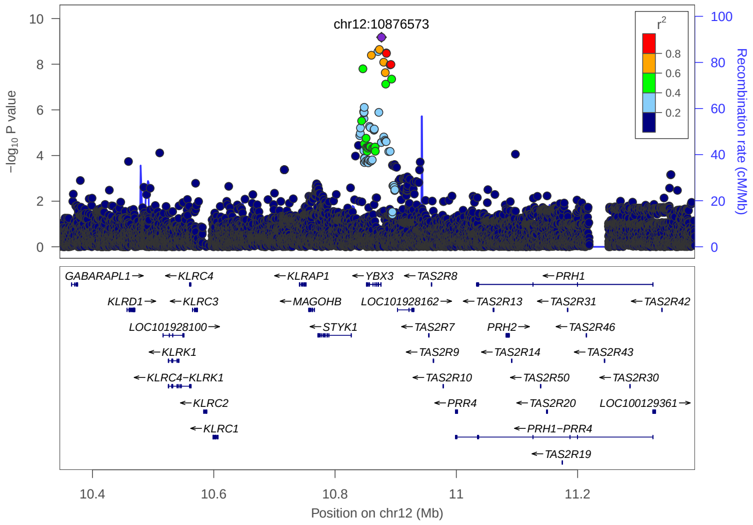


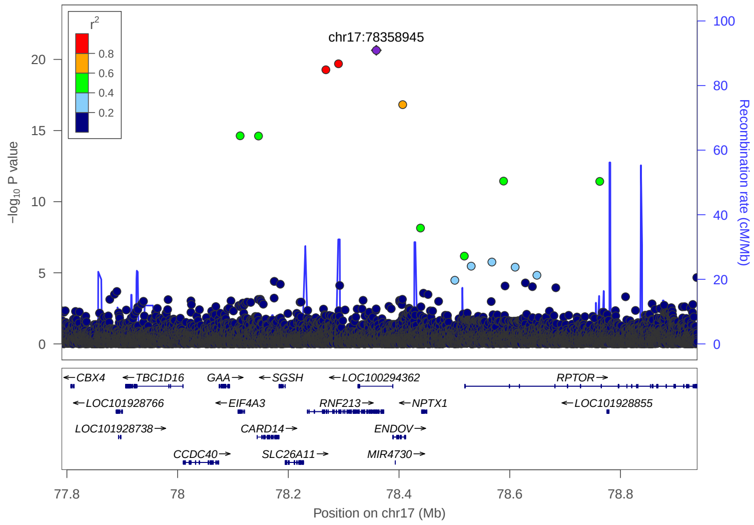


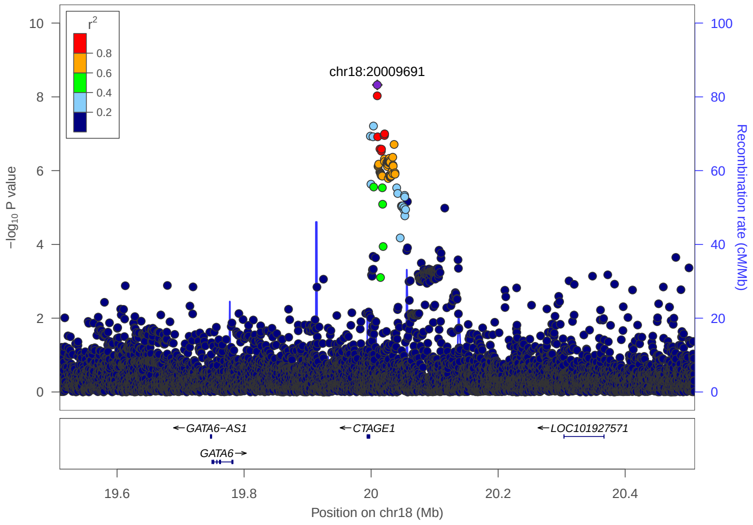

### 2. Previously reported 39 loci in Japanese GWAS

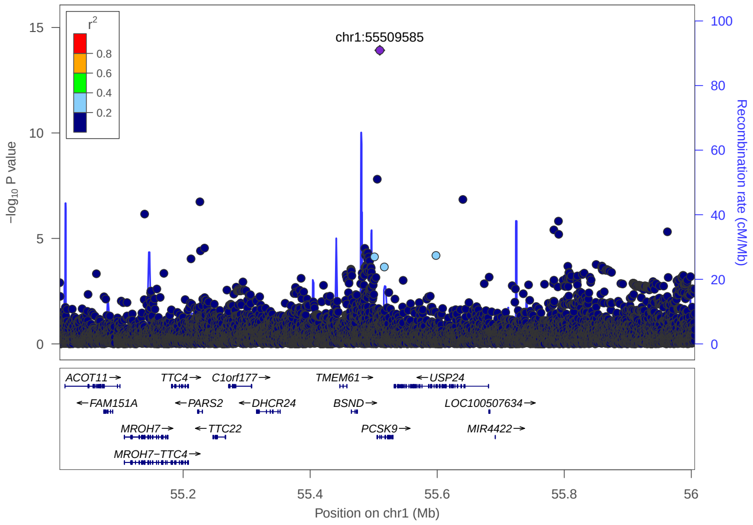


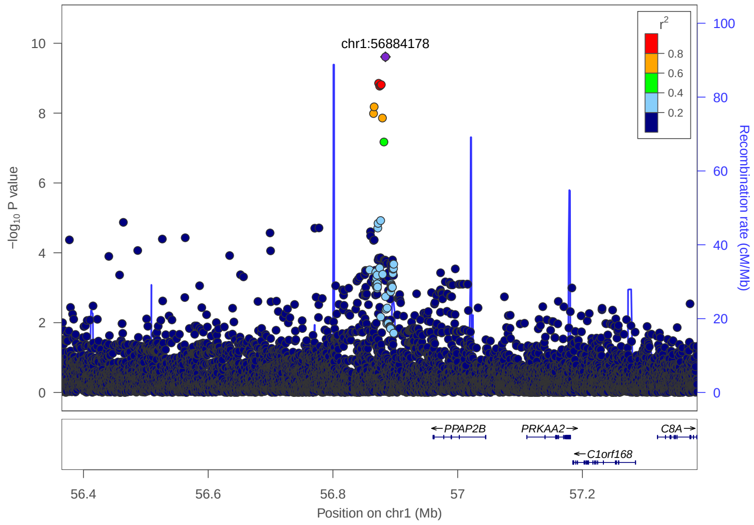


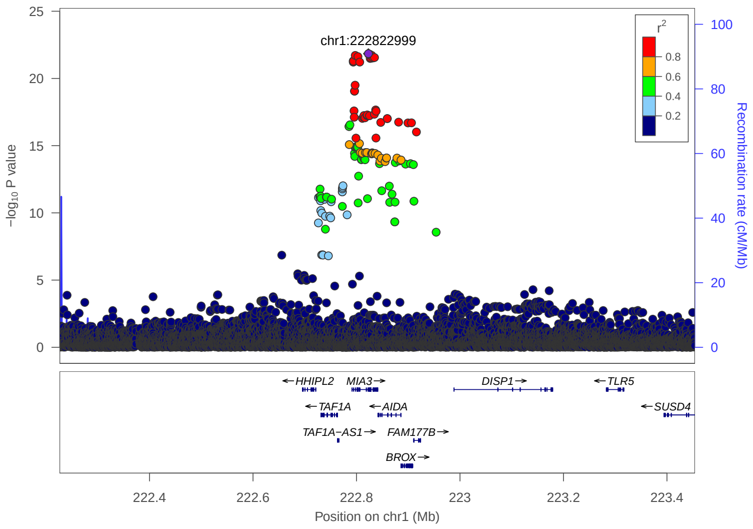


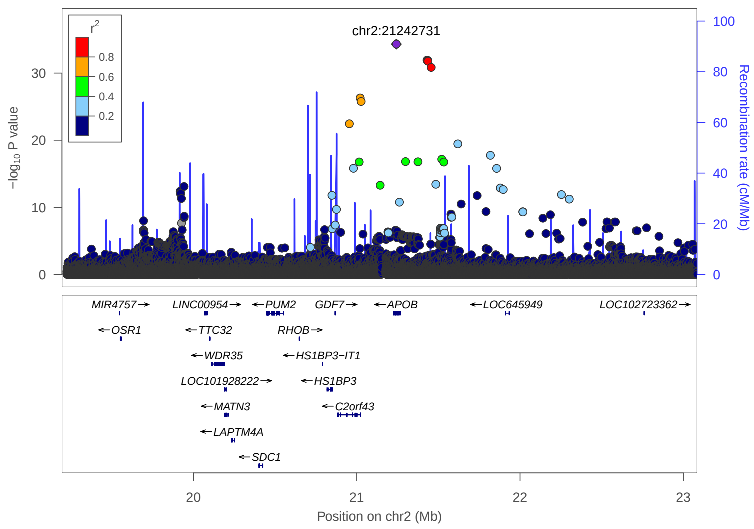


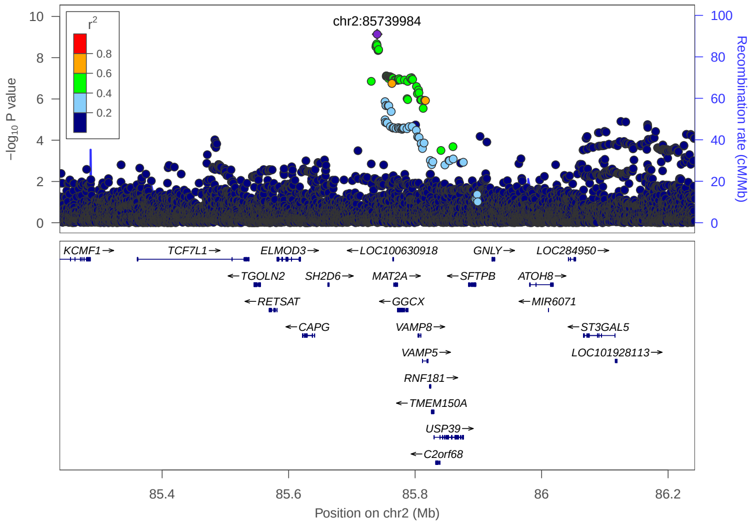


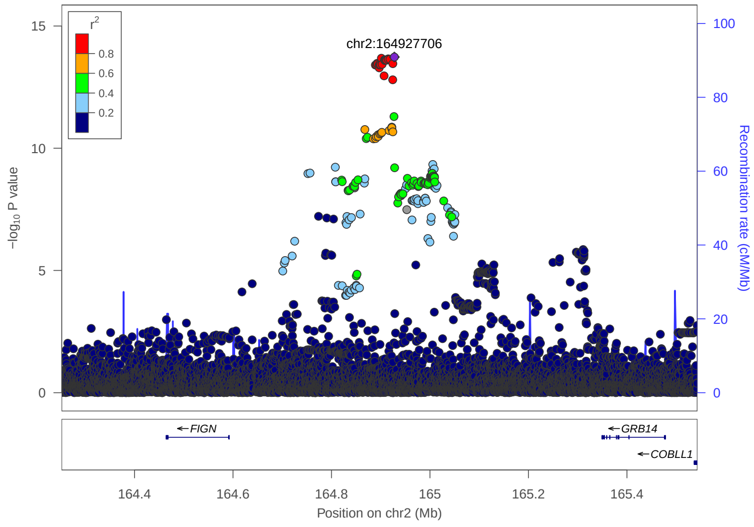

### 2. Previously reported 39 loci in Japanese GWAS

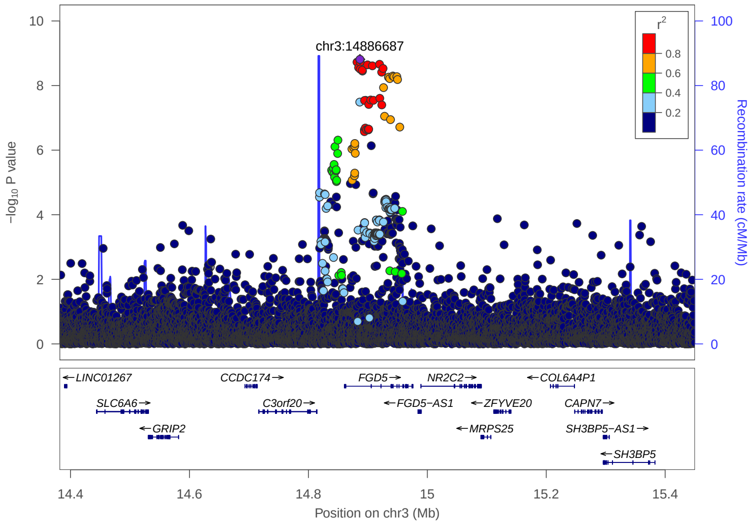


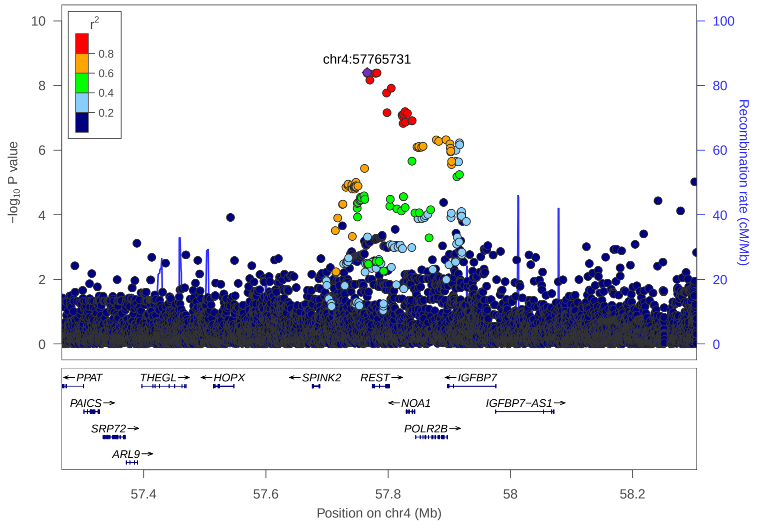


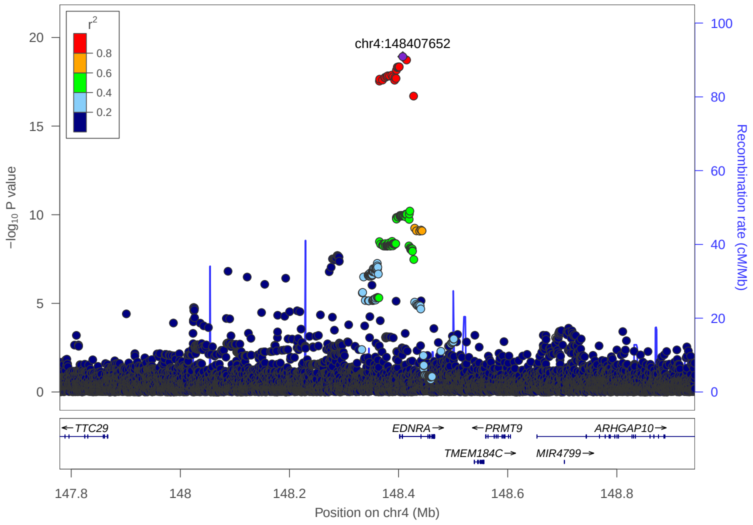


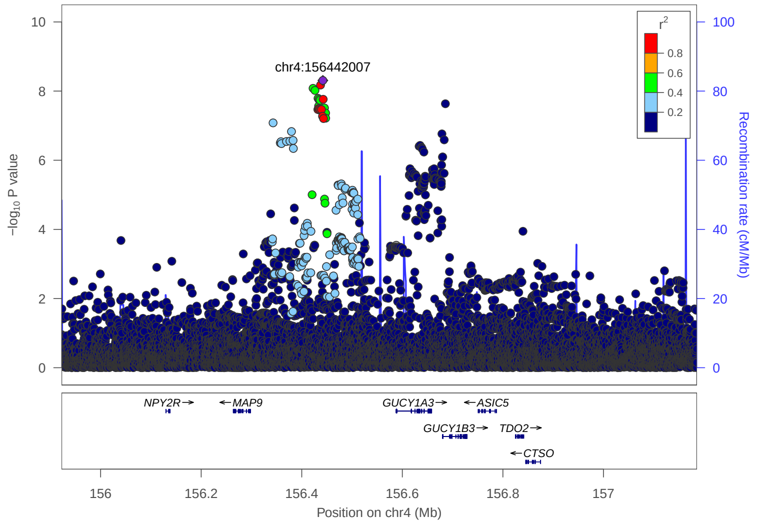


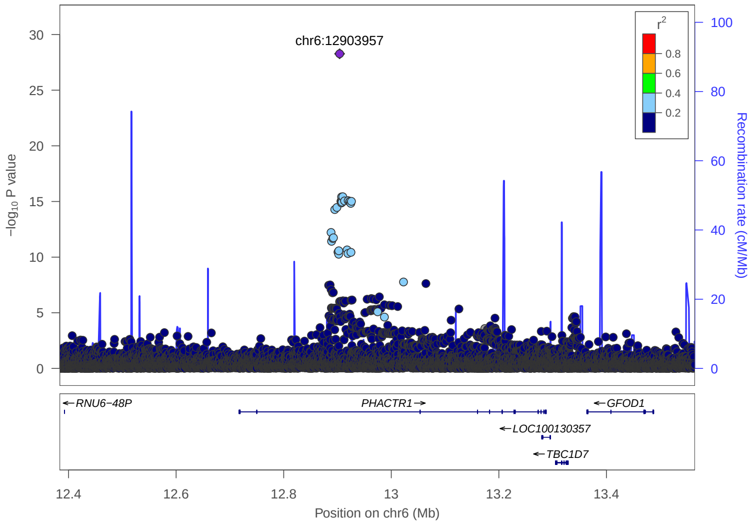


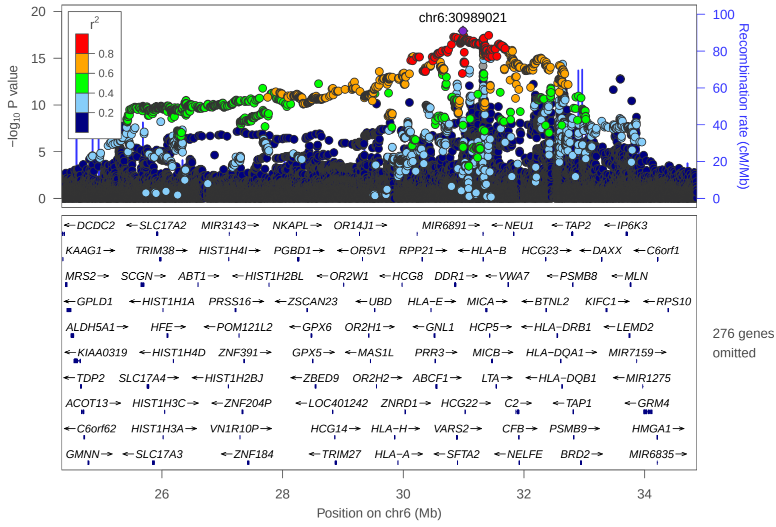

### 2. Previously reported 39 loci in Japanese GWAS

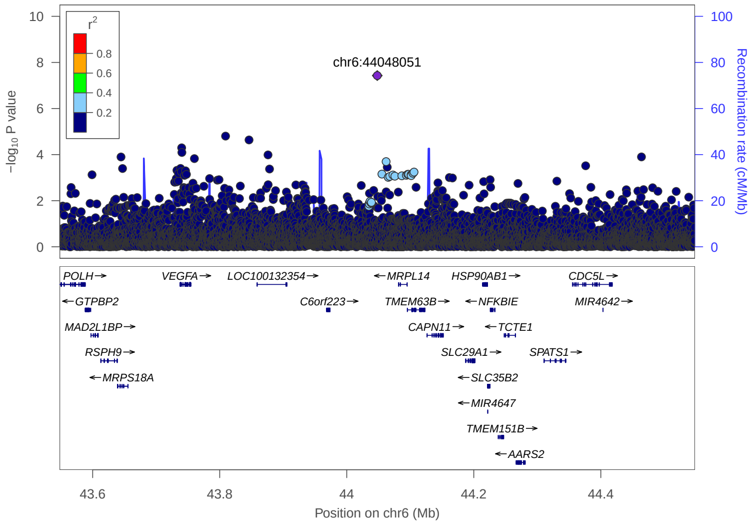


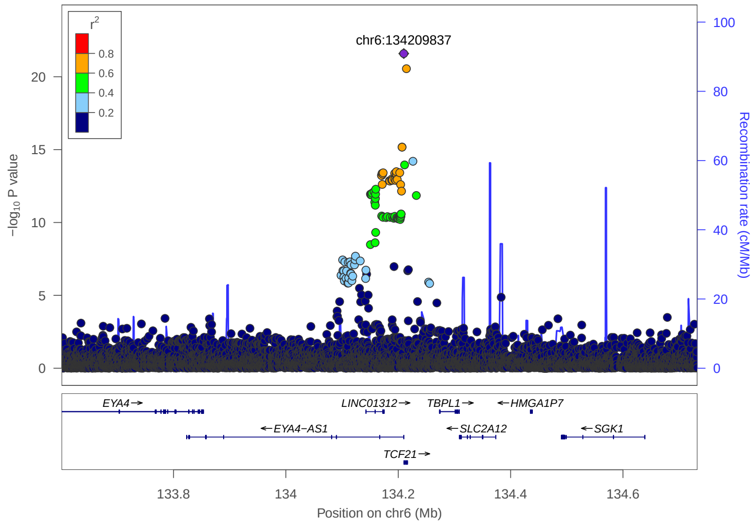


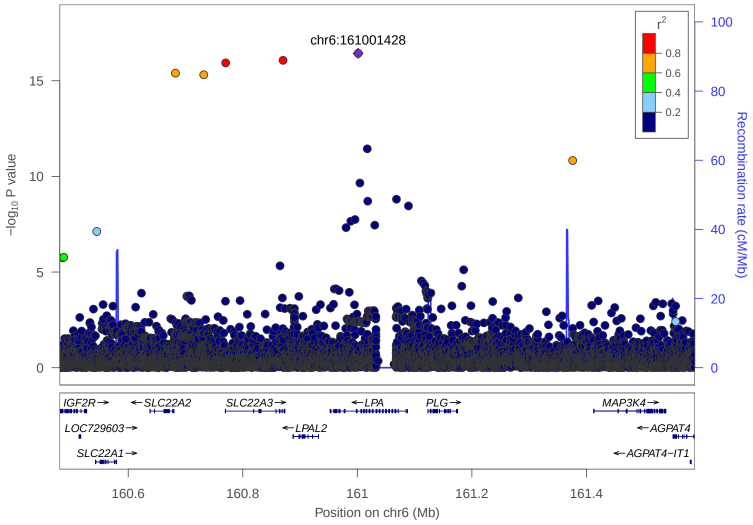


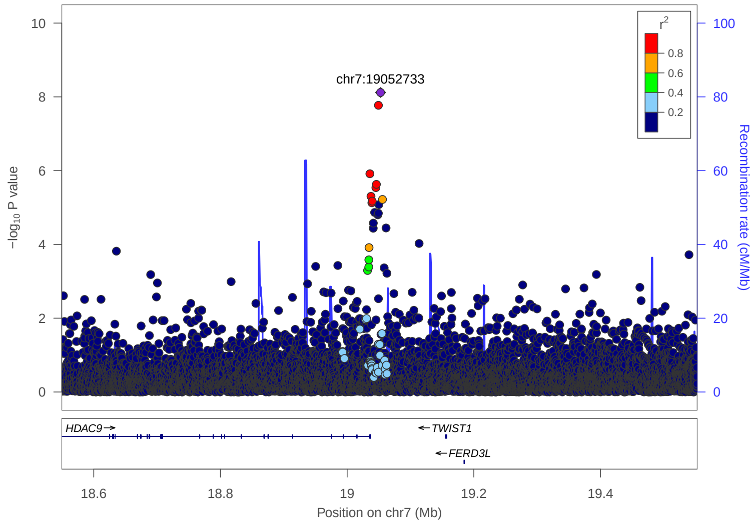


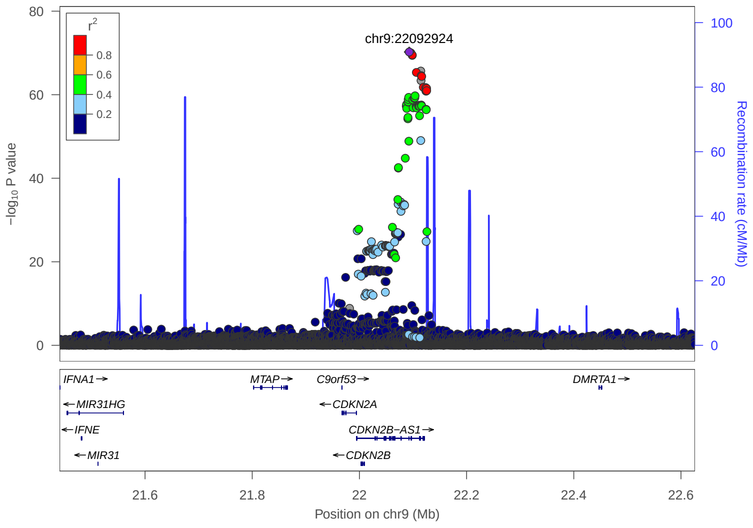


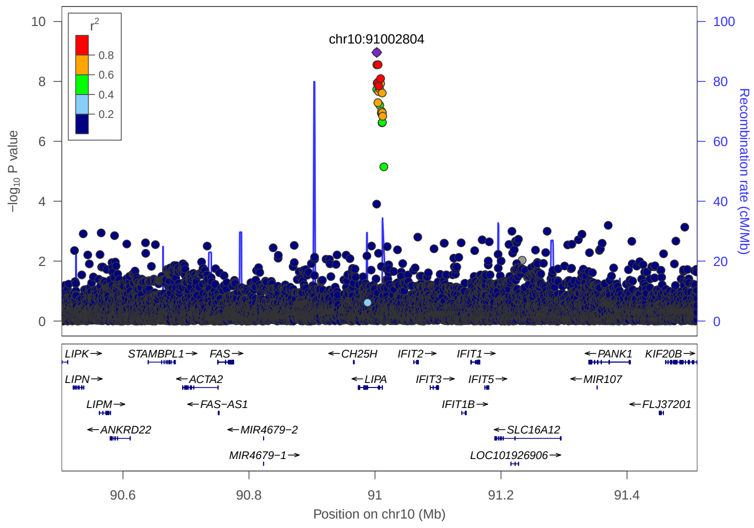

### 2. Previously reported 39 loci in Japanese GWAS

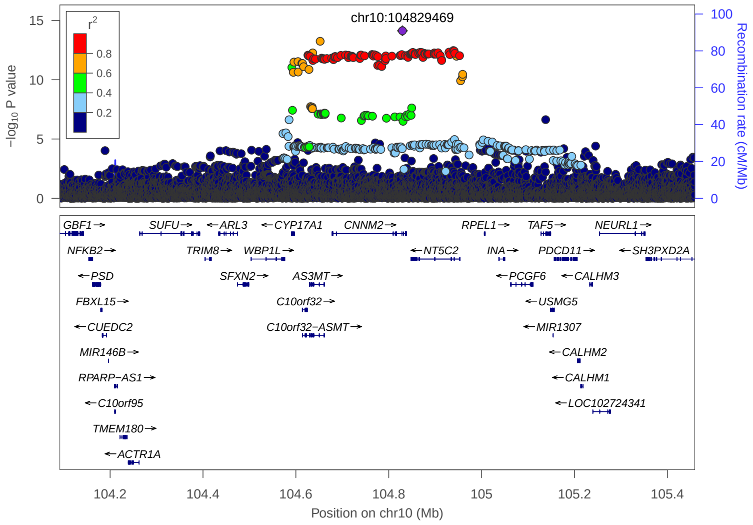


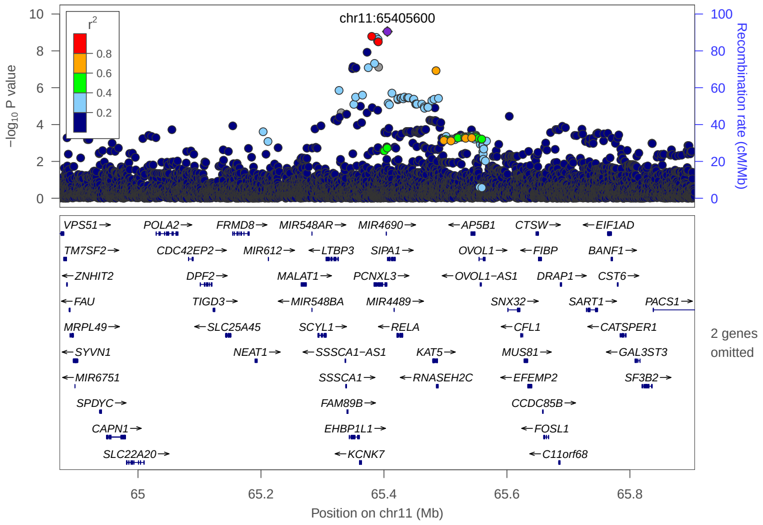


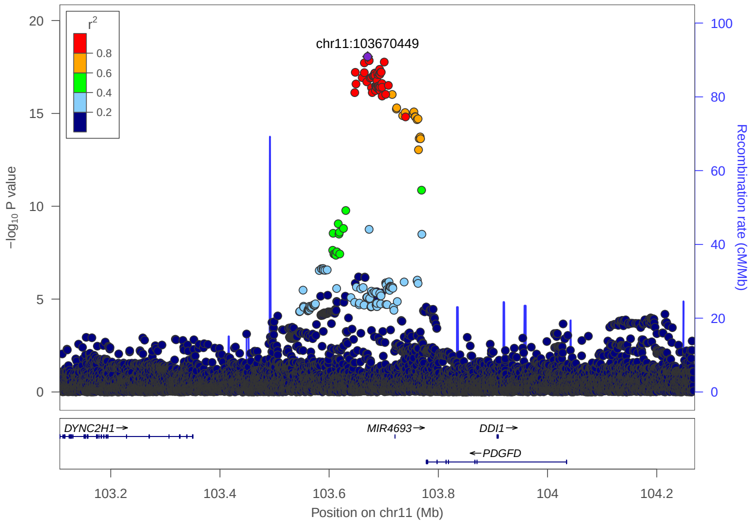


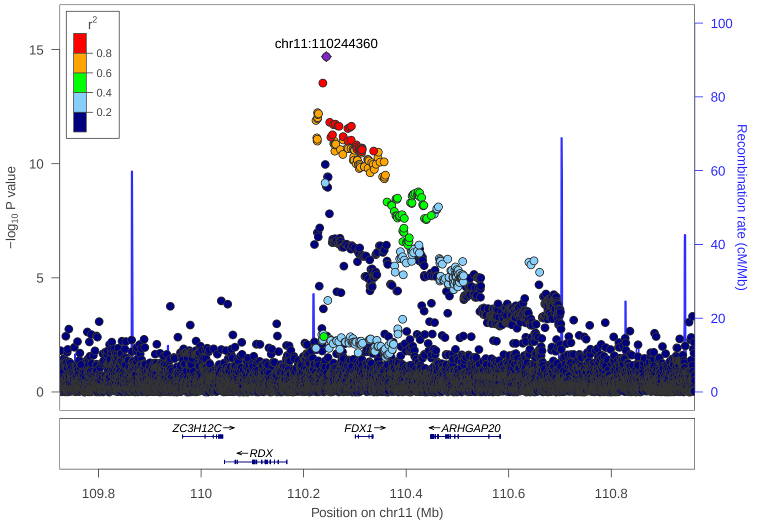


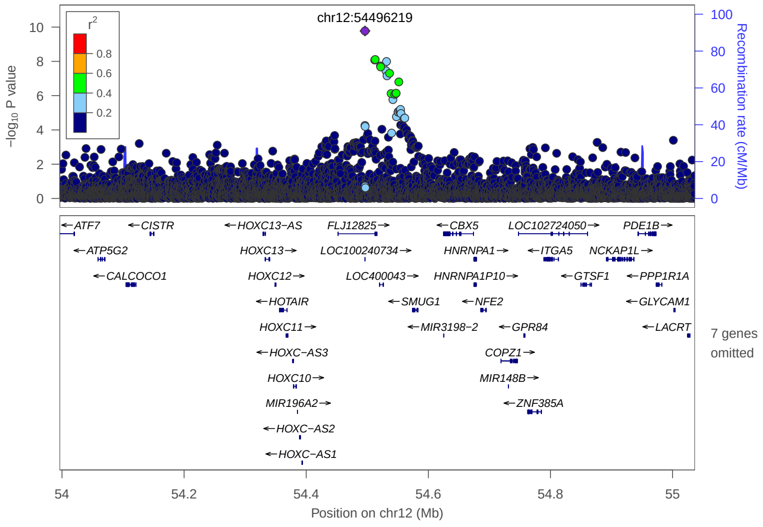


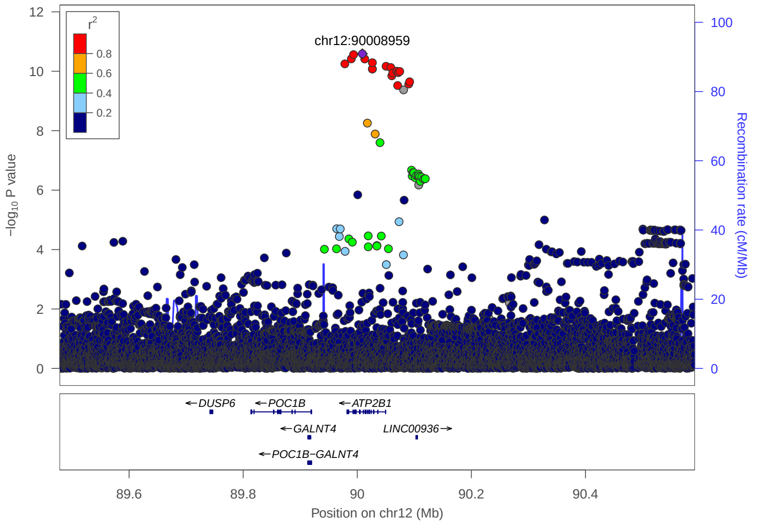

### 2. Previously reported 39 loci in Japanese GWAS

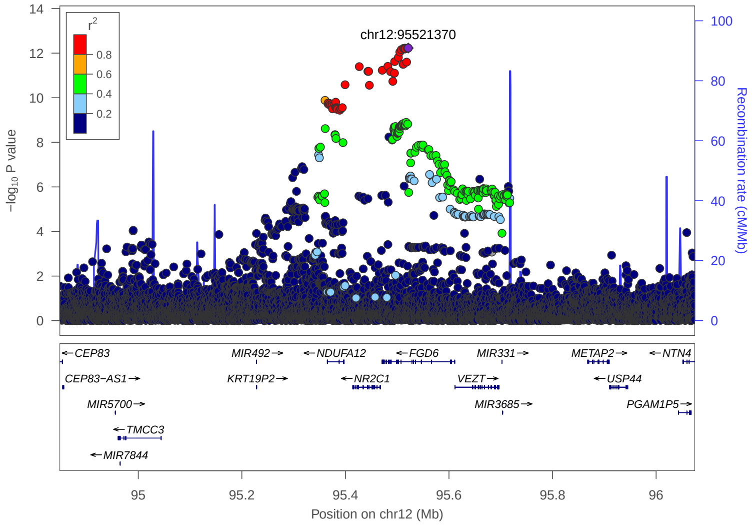


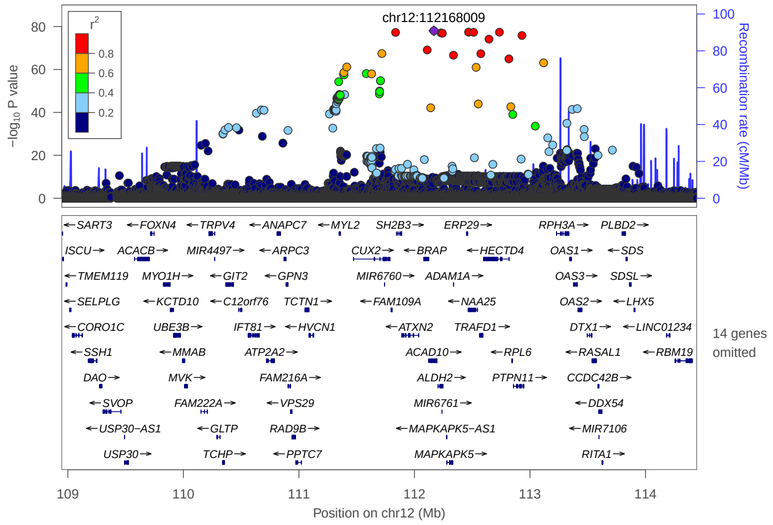


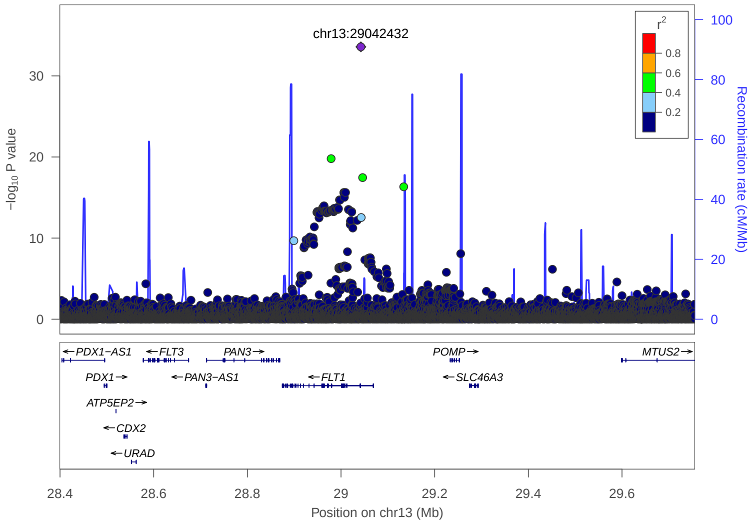


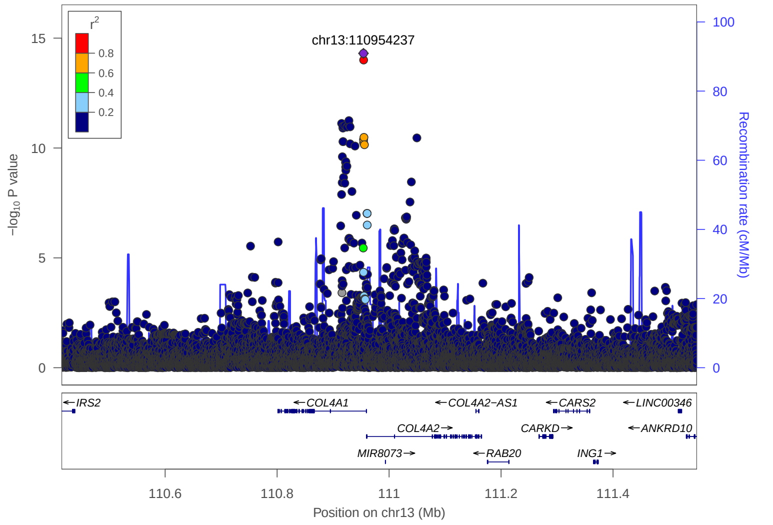


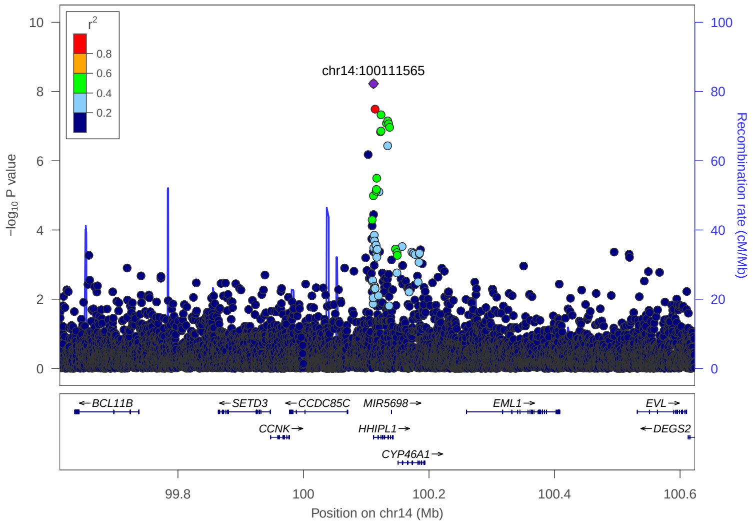


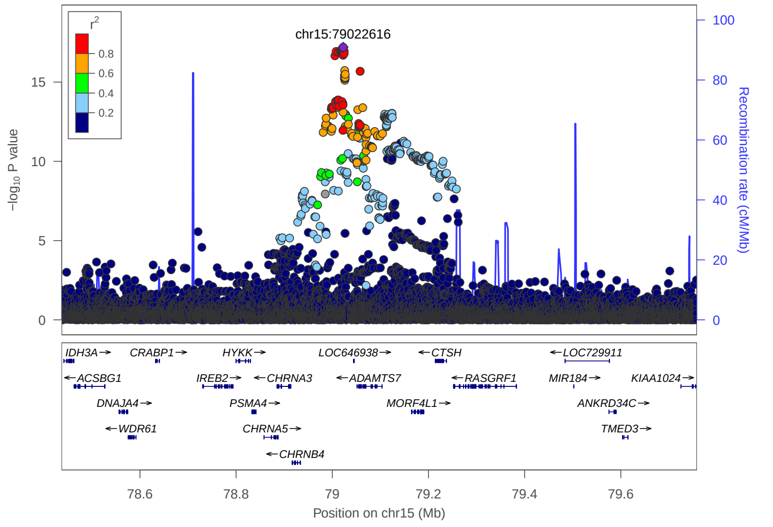

### 2. Previously reported 39 loci in Japanese GWAS

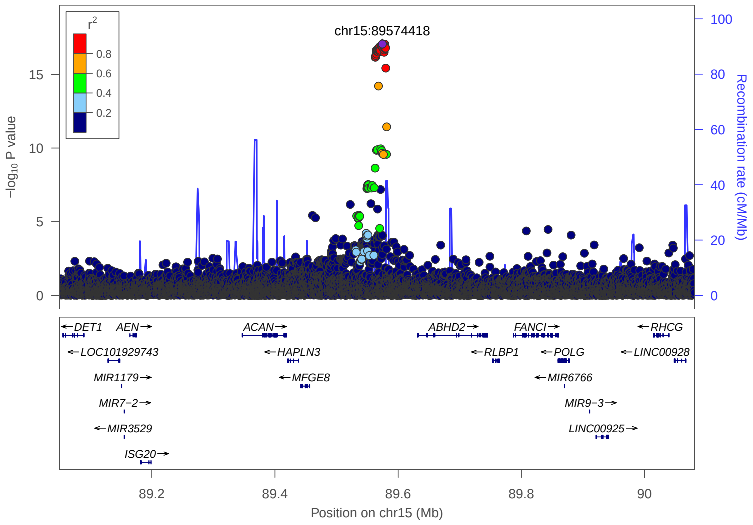


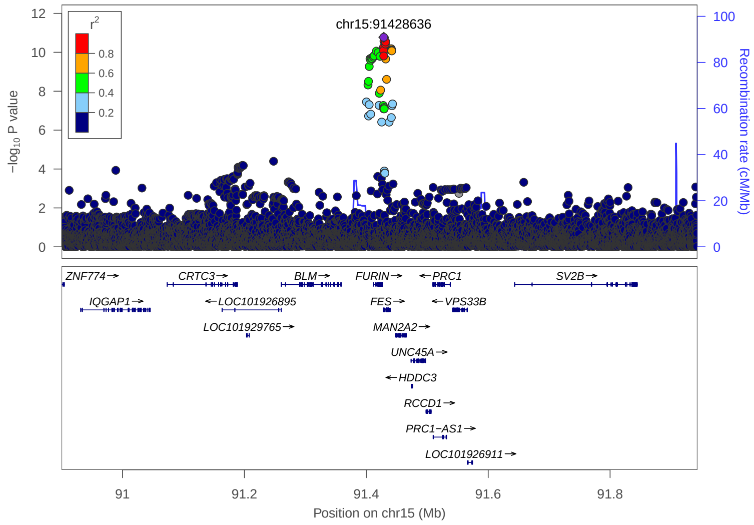


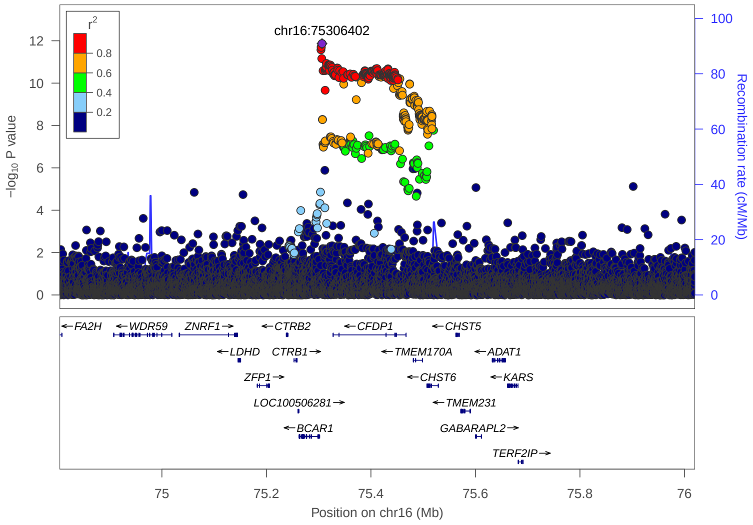


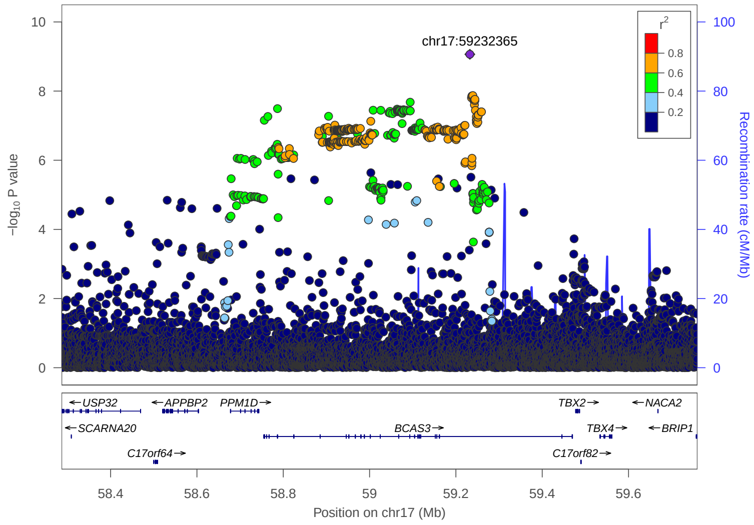


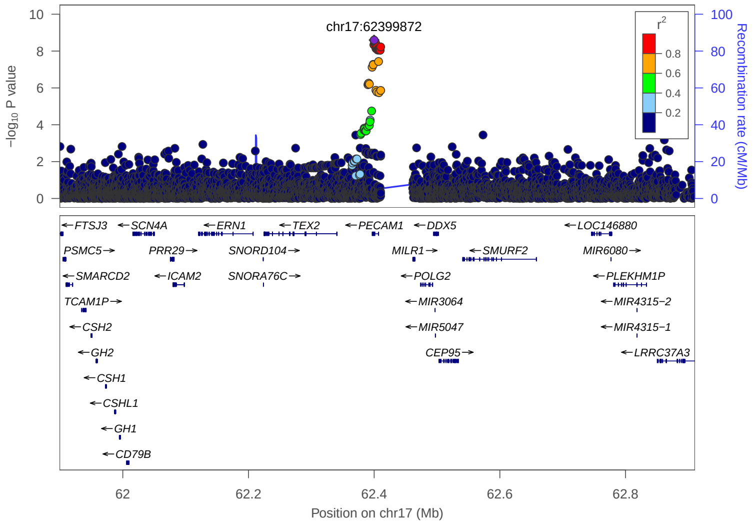


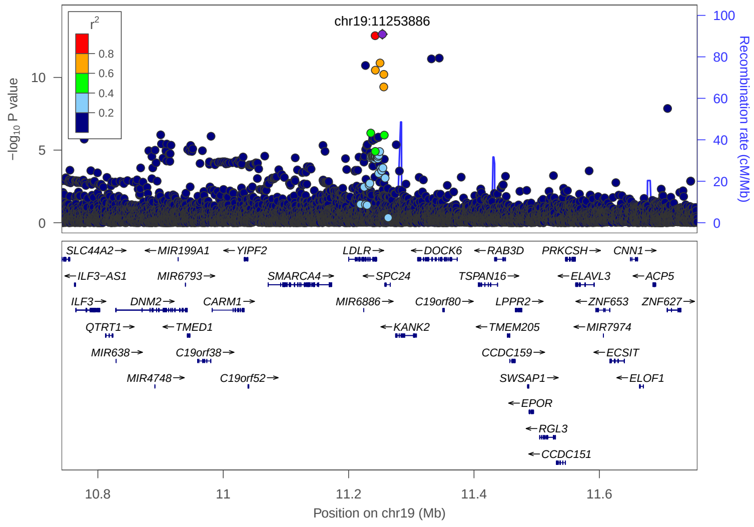

### 2. Previously reported 39 loci in Japanese GWAS

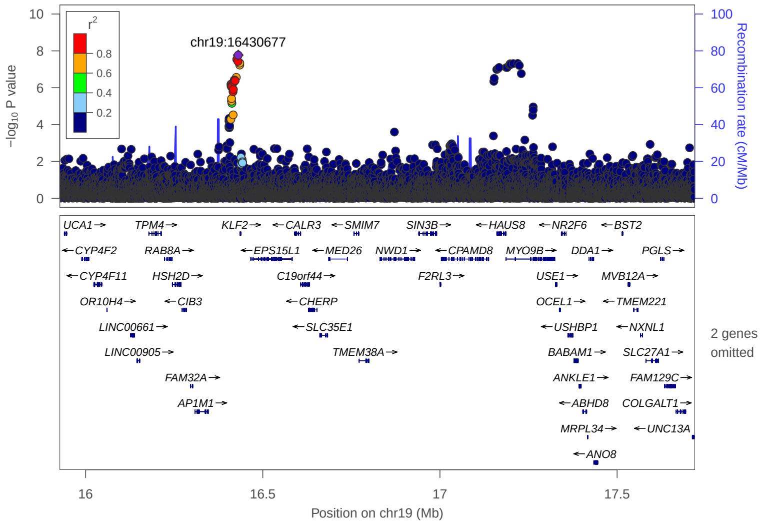


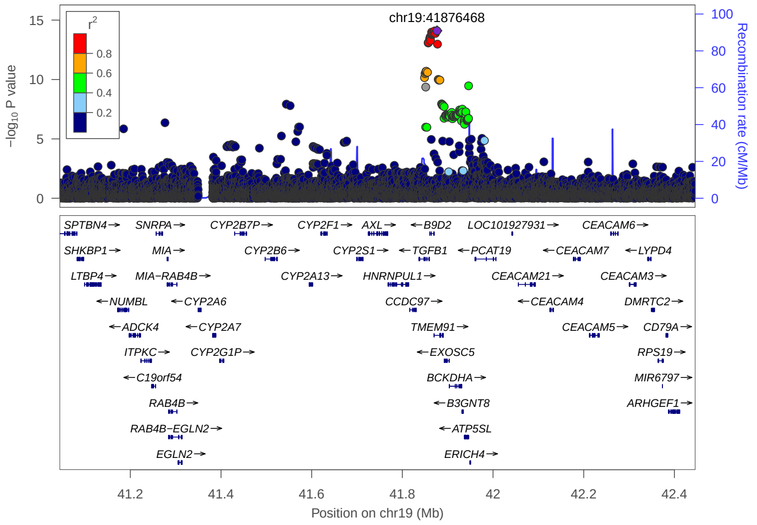


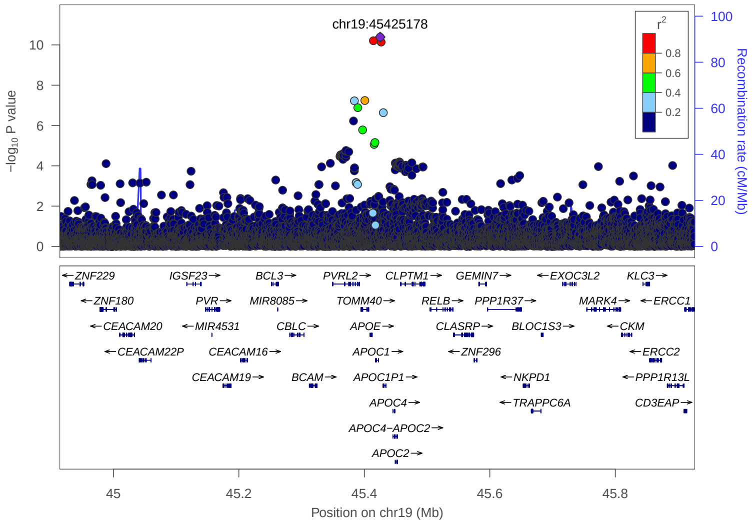

### 3. Newly identified 43 loci in Meta analysis

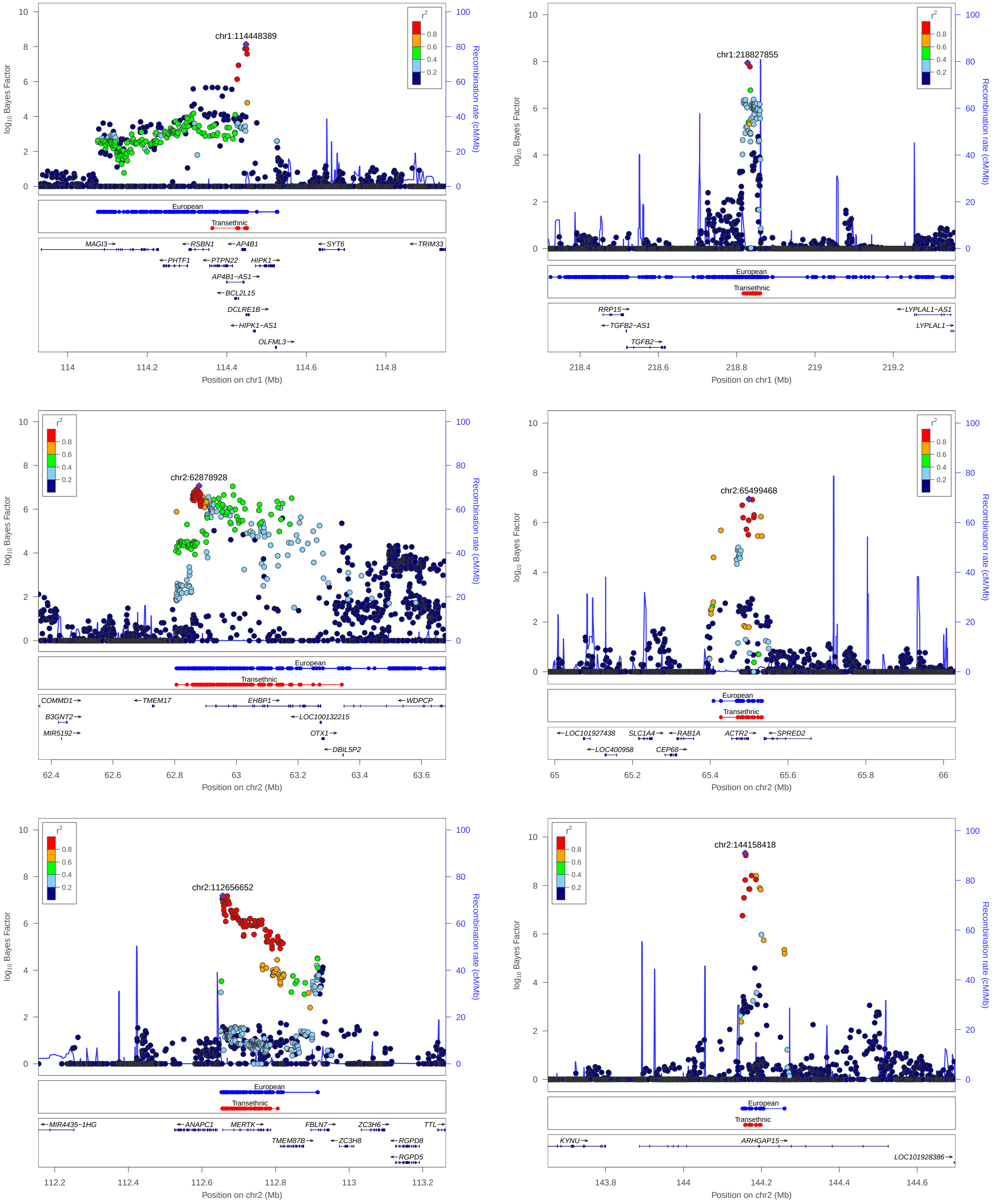


**European:** 99% credible set derived from European meta analysis

**Transethnic:** 99% credible set derived from transethnic meta analysis

### 3. Newly identified 43 loci in Meta analysis

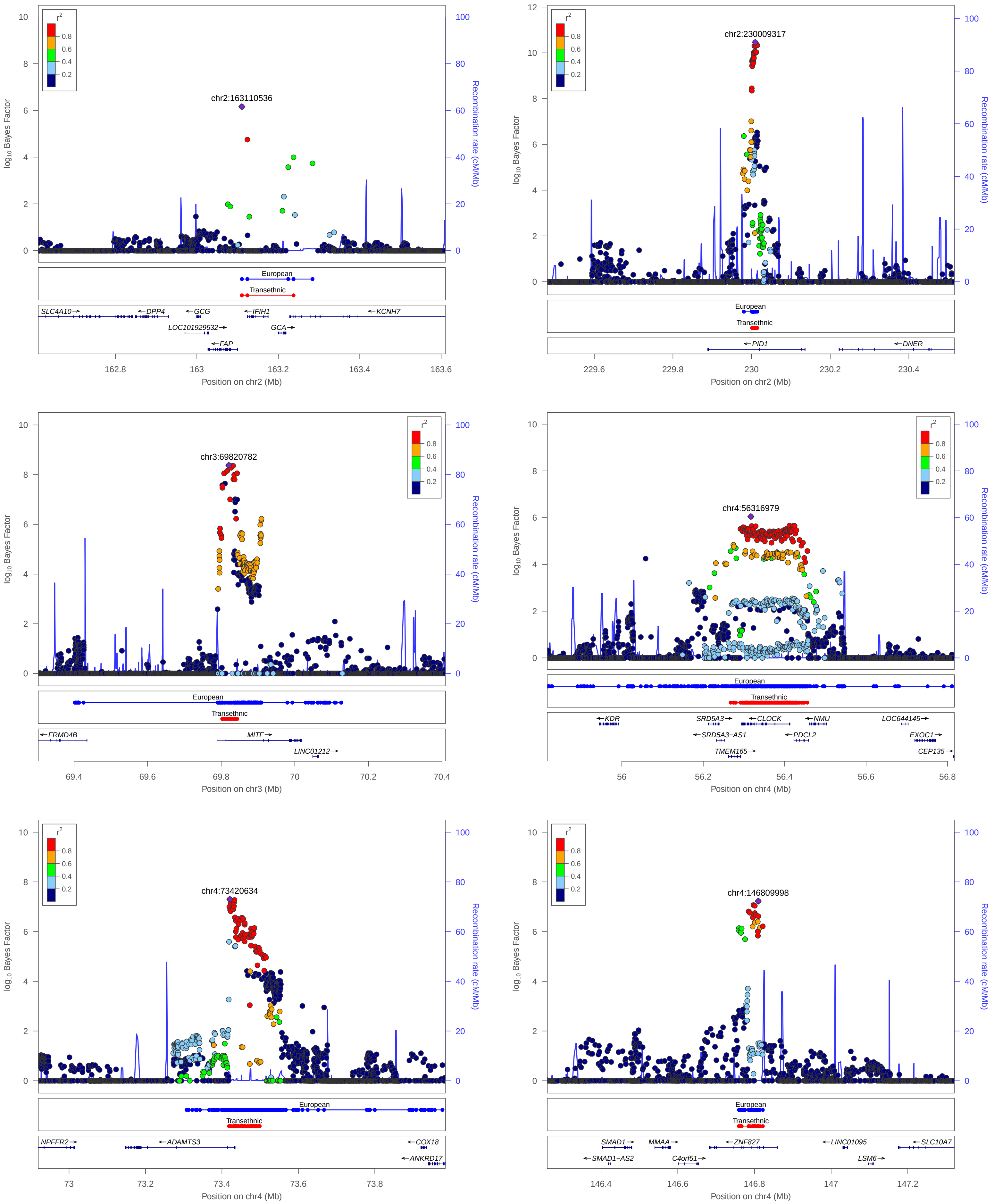


**European:** 99% credible set derived from European meta analysis

**Transethnic:** 99% credible set derived from transethnic meta analysis

### 3. Newly identified 43 loci in Meta analysis

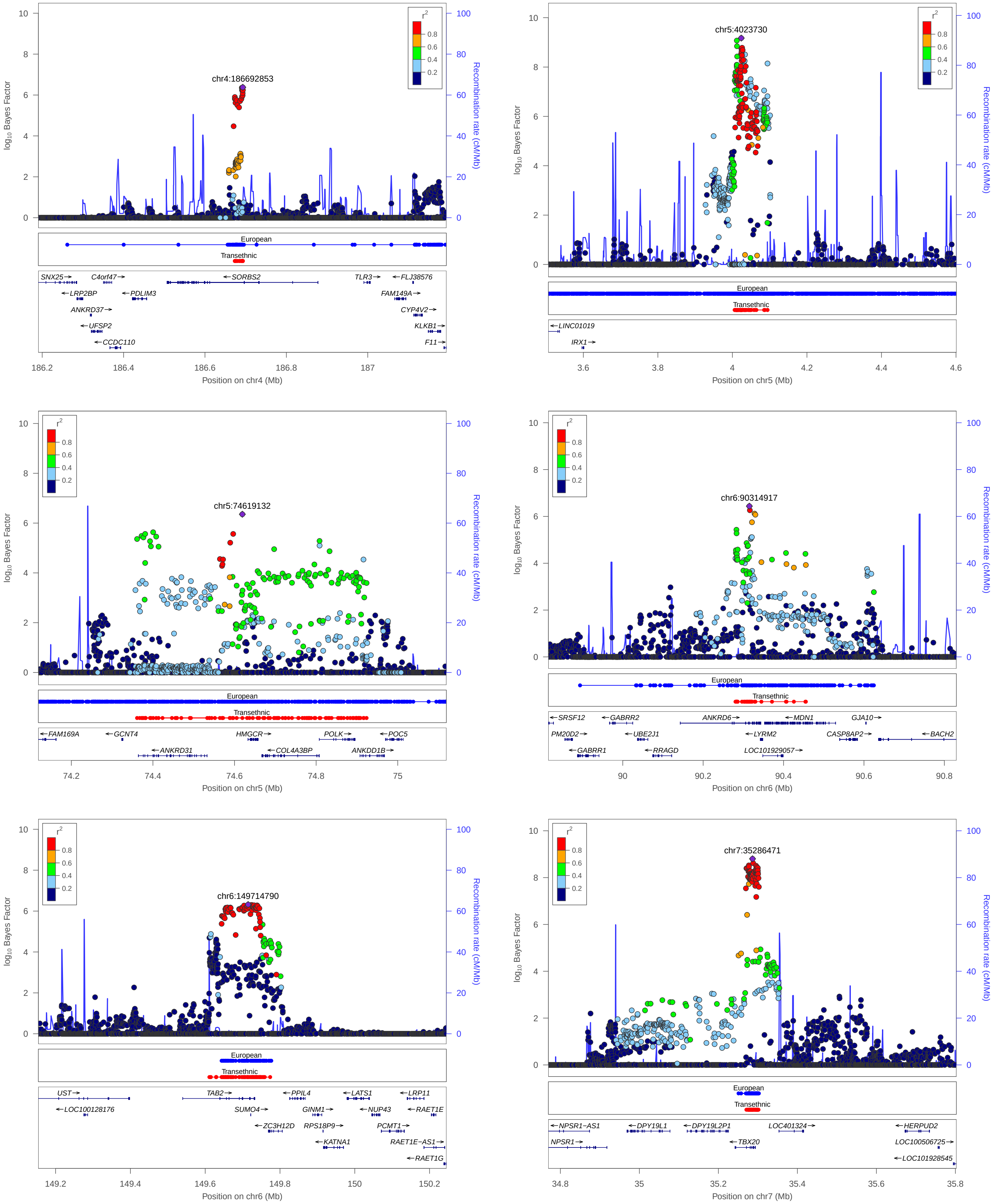


European: 99% credible set derived from European meta analysis

Transethnic: 99% credible set derived from transethnic meta analysis

### 3. Newly identified 43 loci in Meta analysis

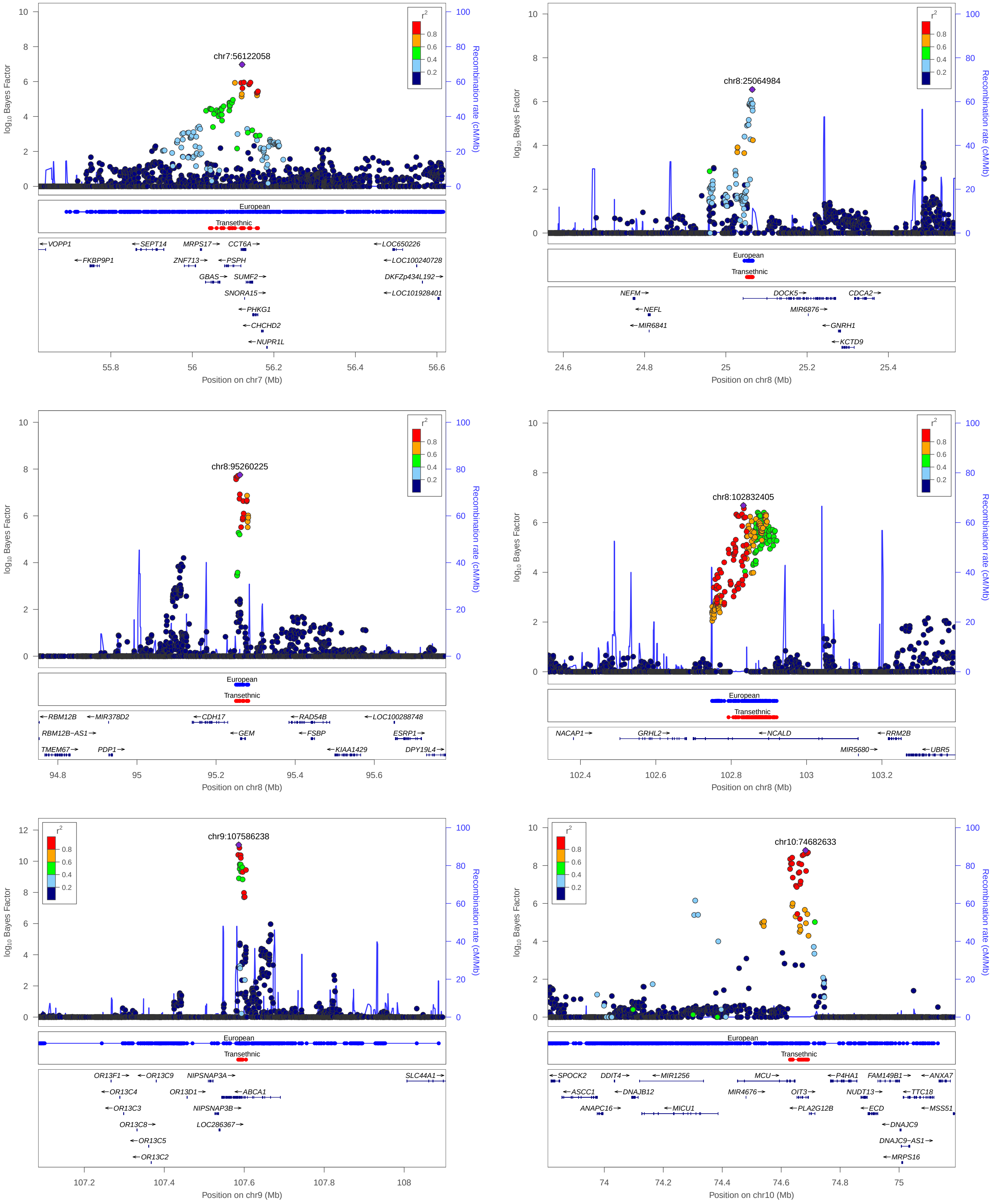


**European:** 99% credible set derived from European meta analysis

**Transethnic:** 99% credible set derived from transethnic meta analysis

### 3. Newly identified 43 loci in Meta analysis

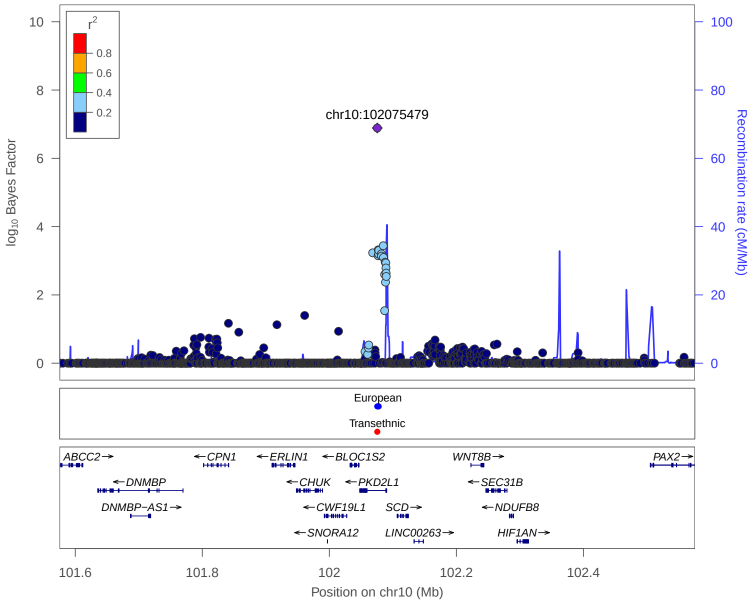


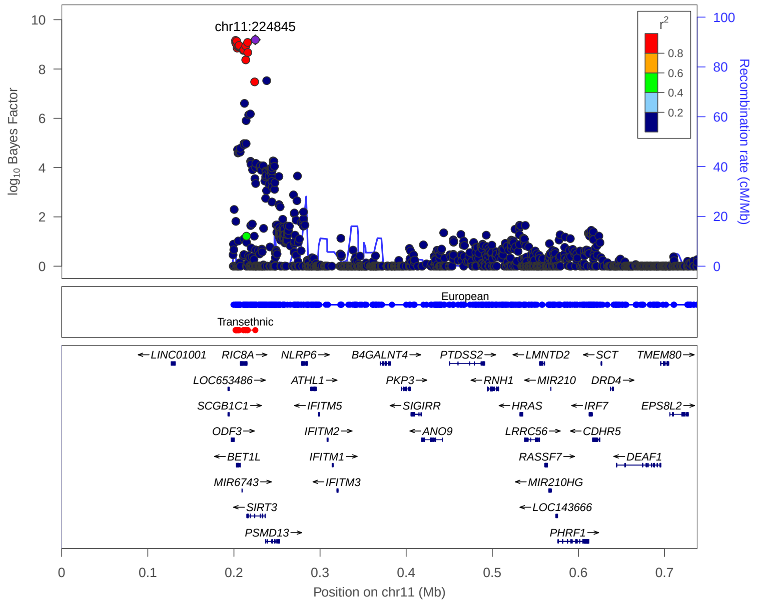


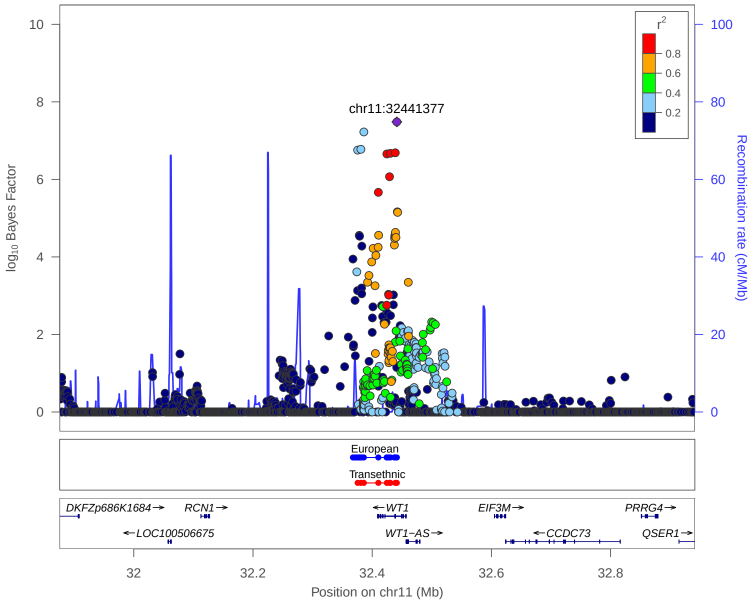


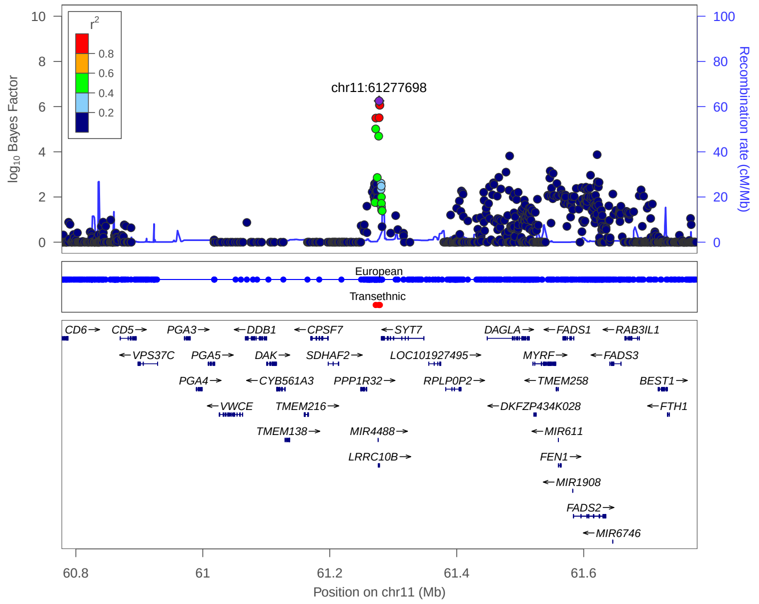


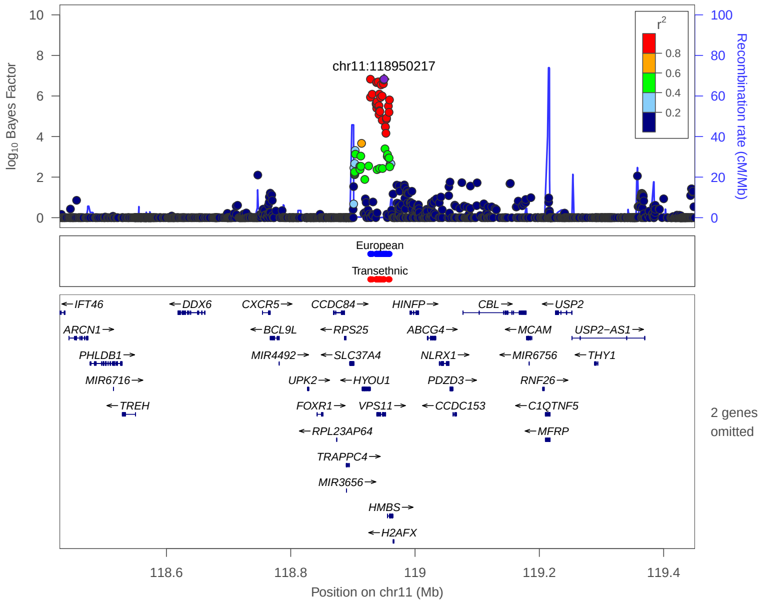


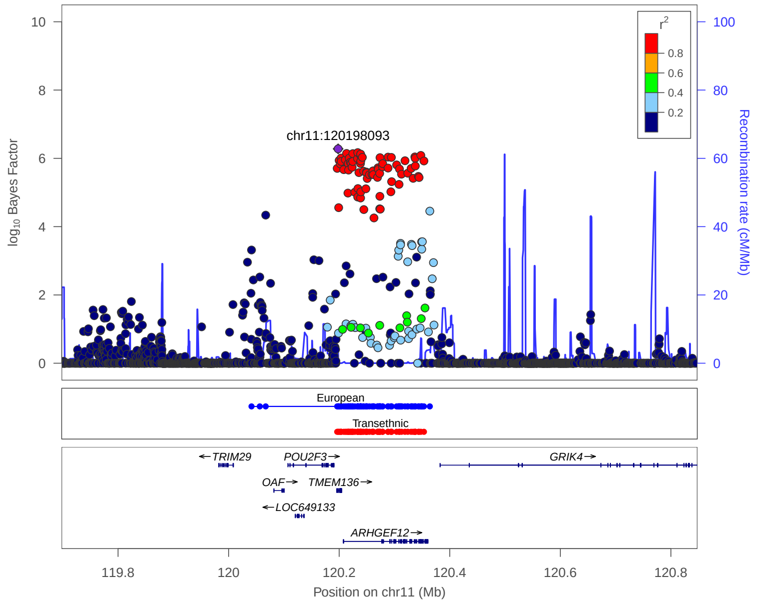


**European:** 99% credible set derived from European meta analysis

**Transethnic:** 99% credible set derived from transethnic meta analysis

### 3. Newly identified 43 loci in Meta analysis

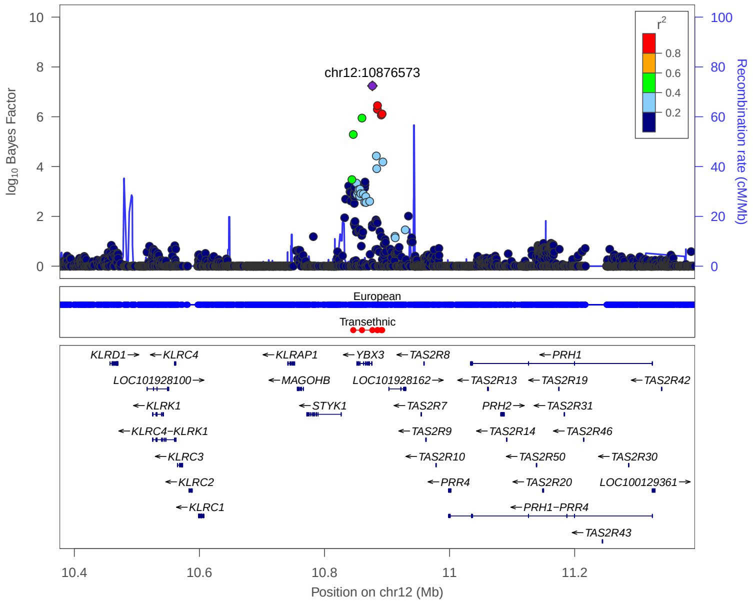


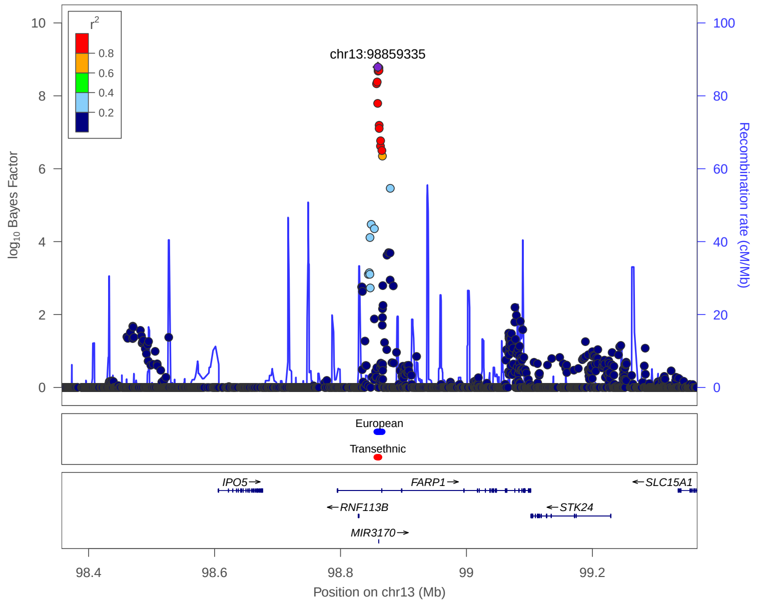


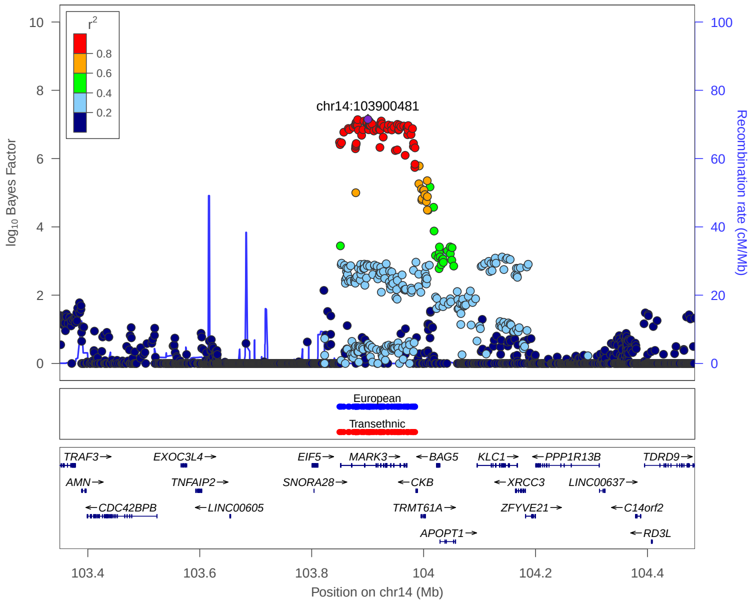


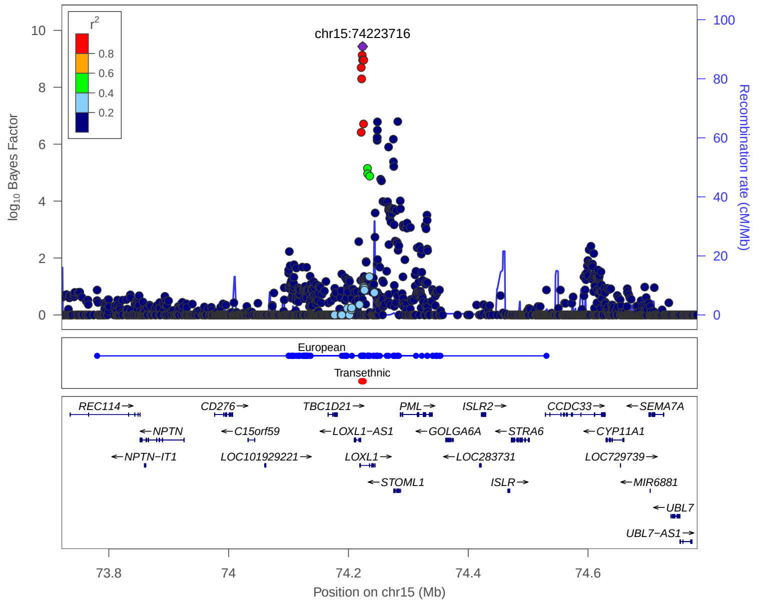


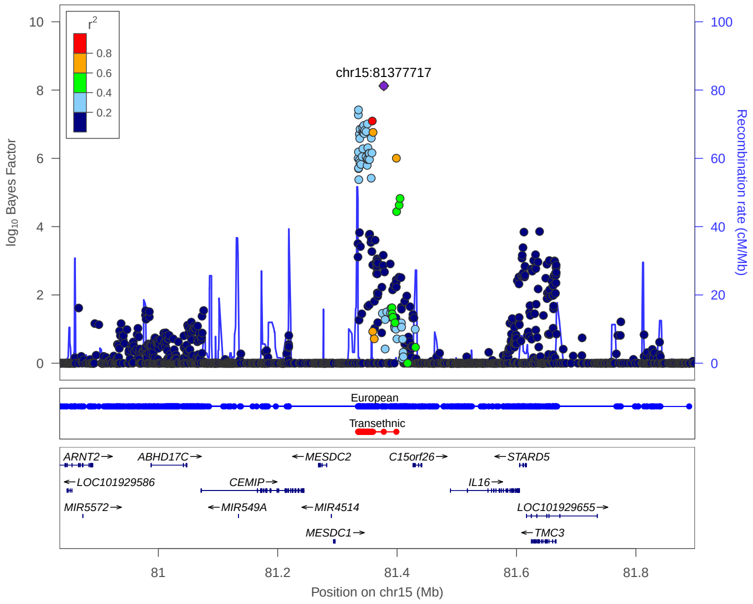


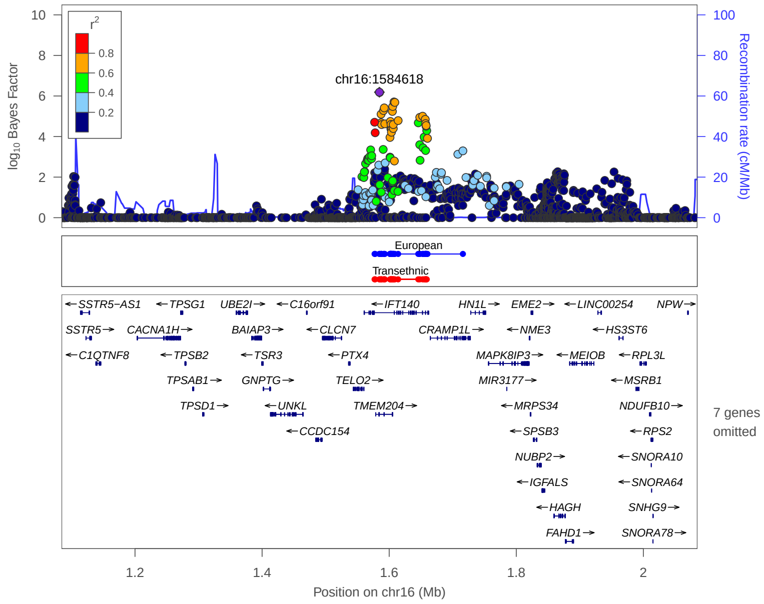


**European:** 99% credible set derived from European meta analysis

**Transethnic:** 99% credible set derived from transethnic meta analysis

### 3. Newly identified 43 loci in Meta analysis

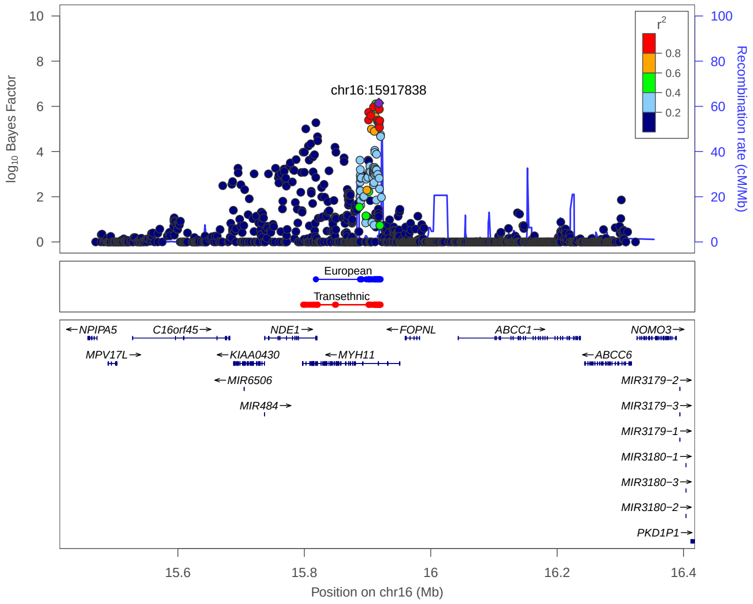


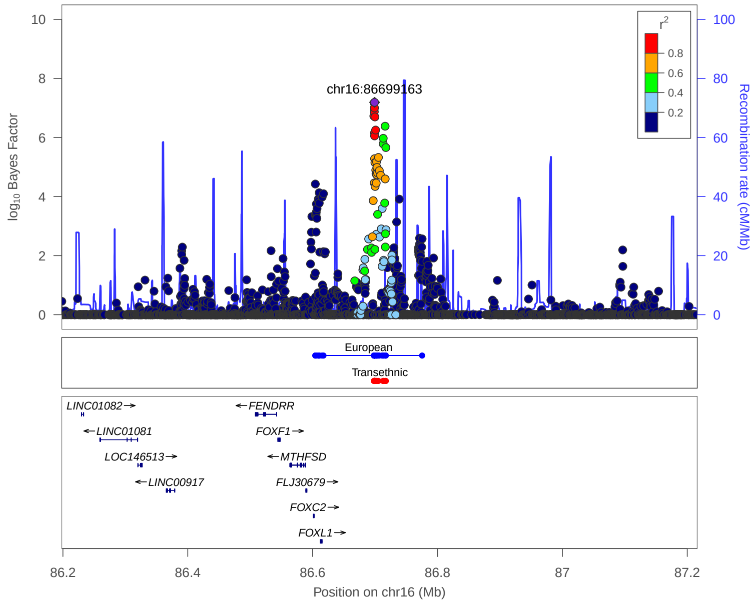


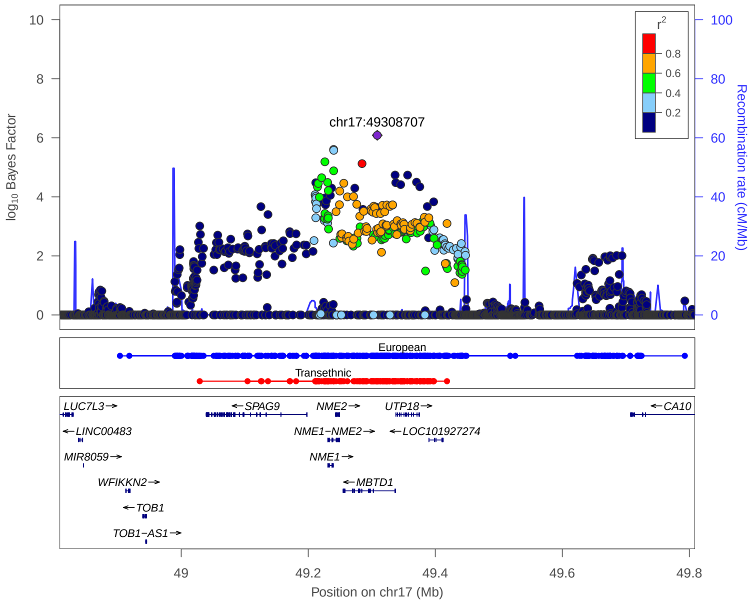


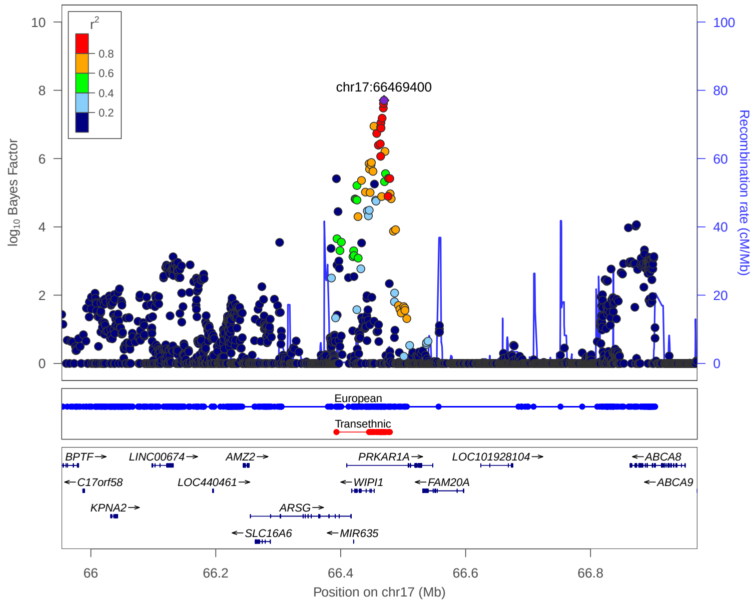


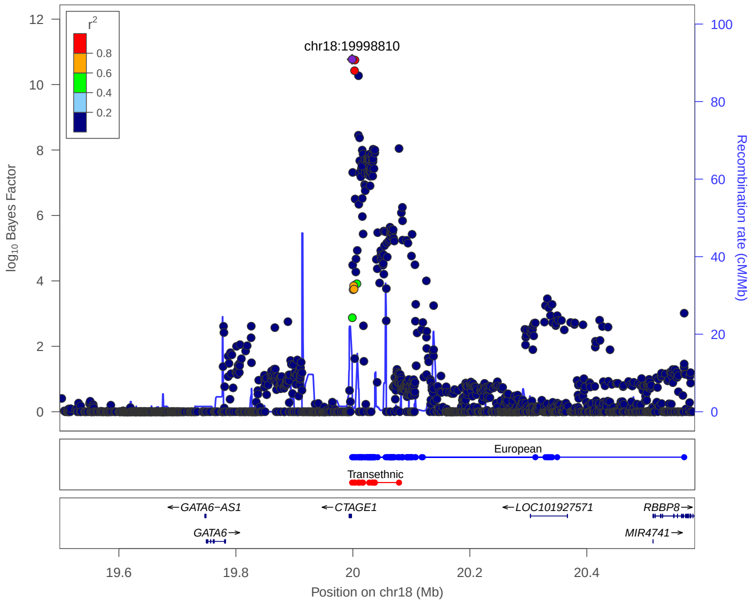


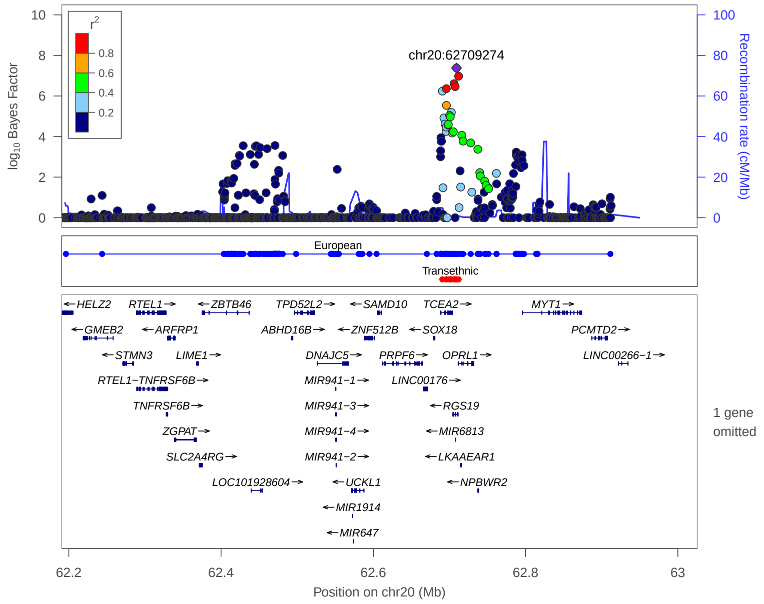


**European:** 99% credible set derived from European meta analysis

**Transethnic:** 99% credible set derived from transethnic meta analysis

### 3. Newly identified 43 loci in Meta analysis

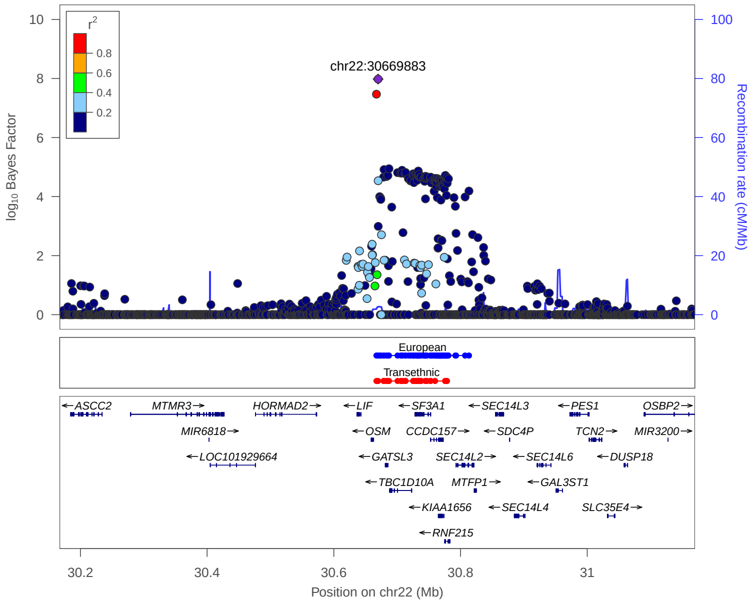


European: 99% credible set derived from European meta analysis

Transethnic: 99% credible set derived from transethnic meta analysis

### 4. Previously reported 133 loci in Meta analysis

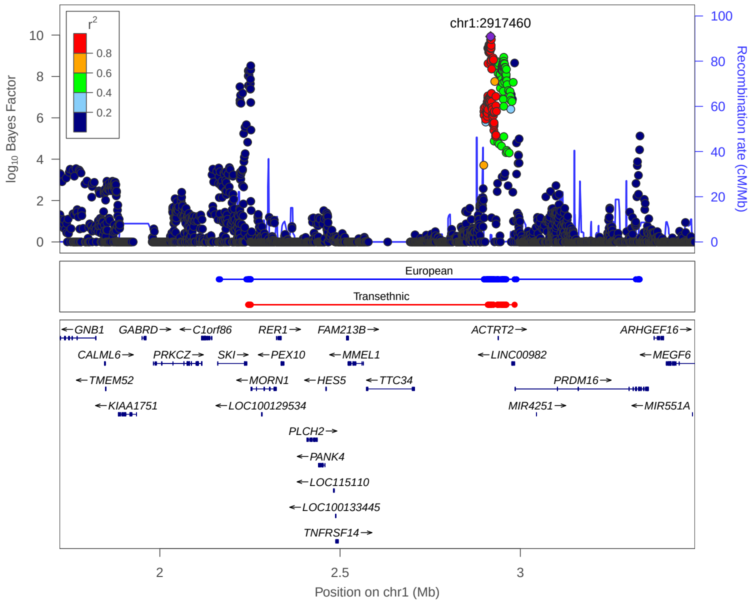


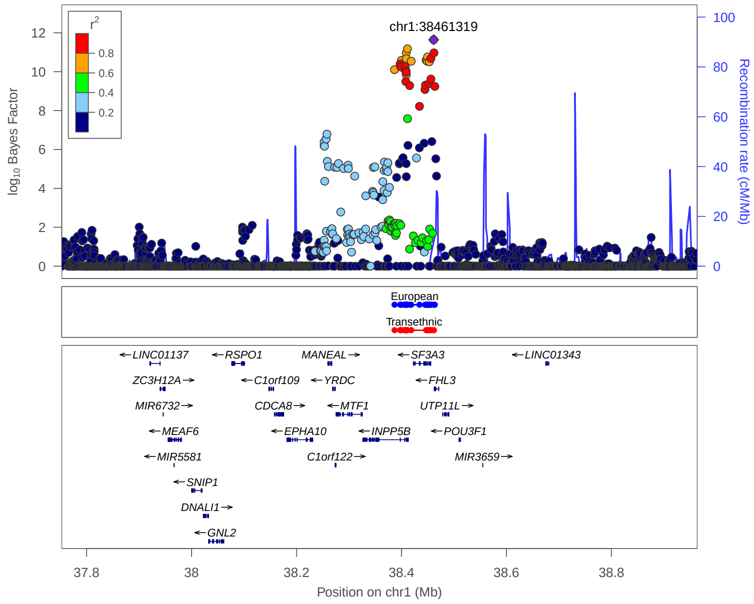


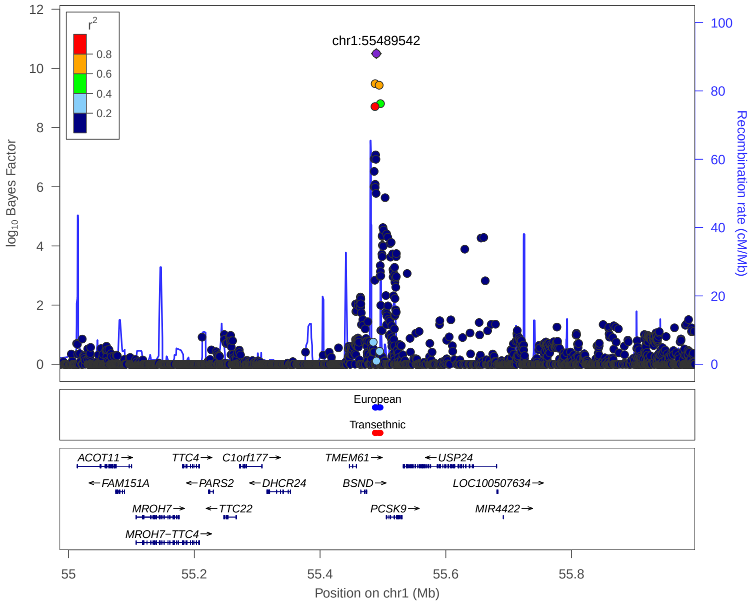


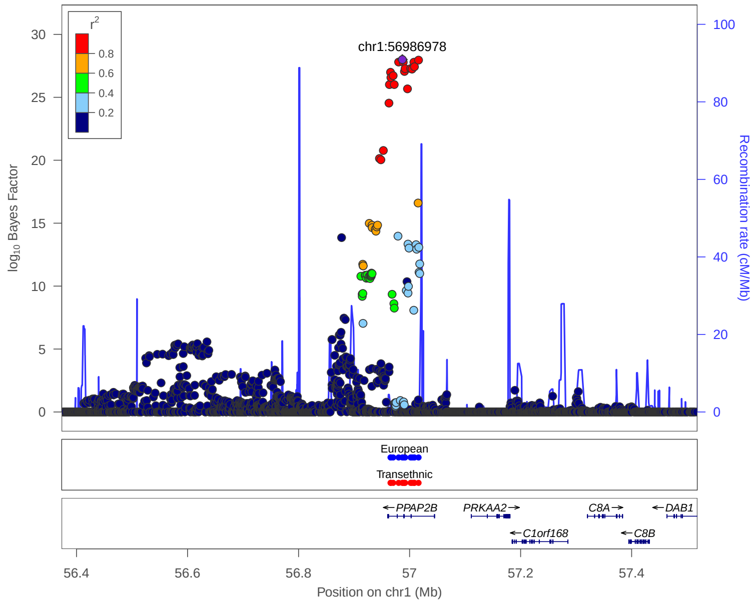


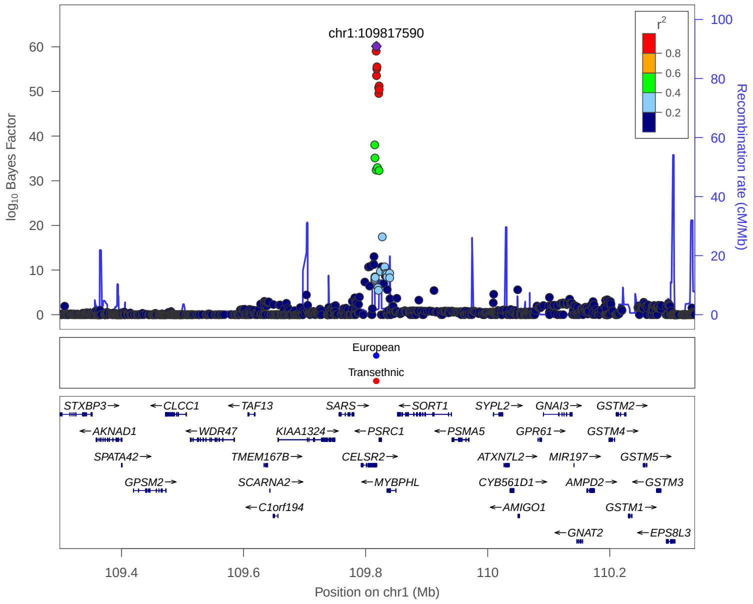


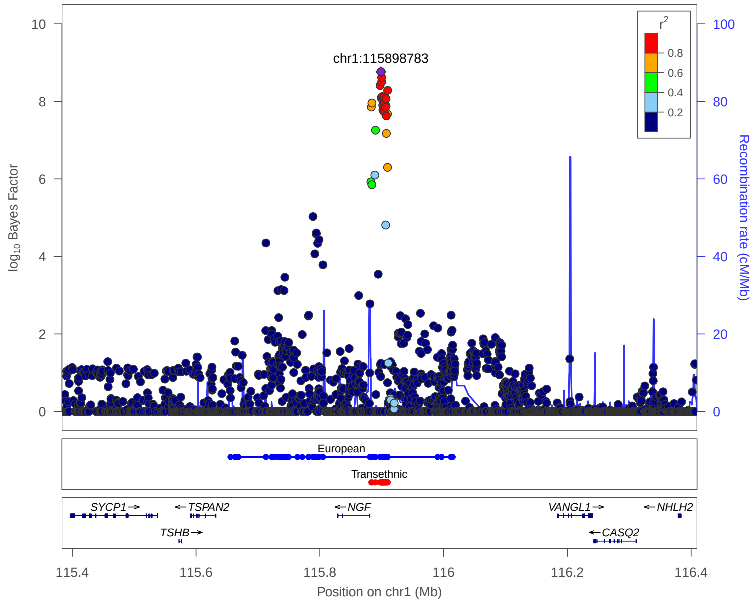


**European:** 99% credible set derived from European meta analysis

**Transethnic:** 99% credible set derived from transethnic meta analysis

### 4. Previously reported 133 loci in Meta analysis

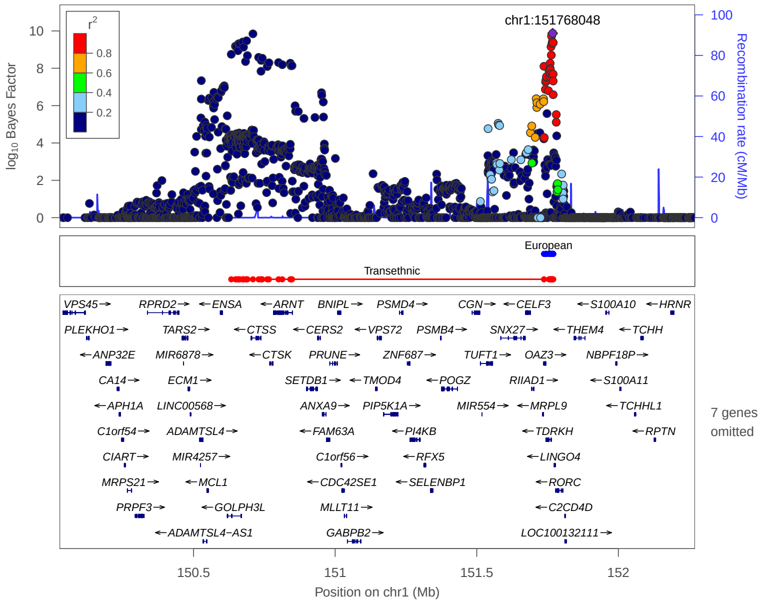


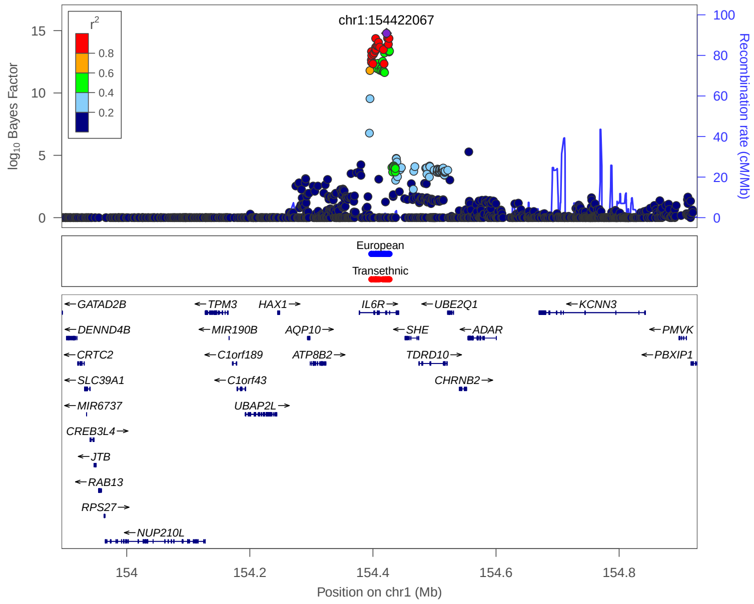


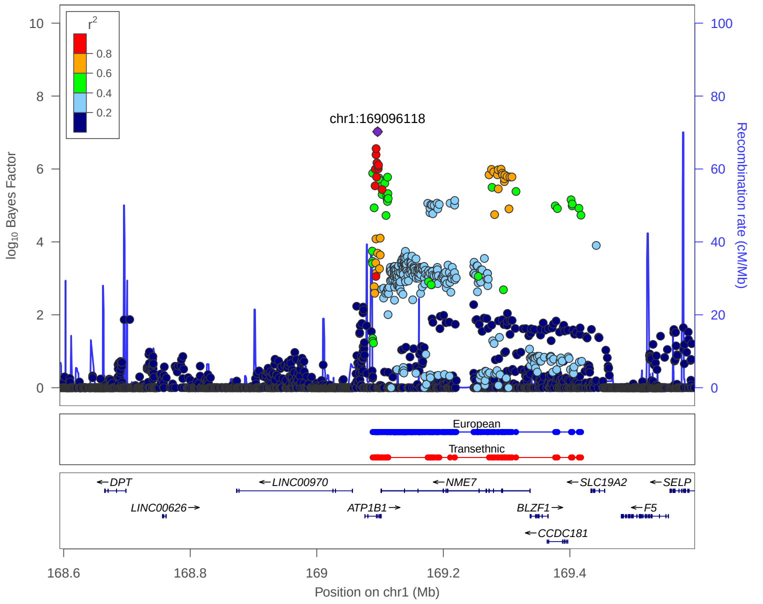


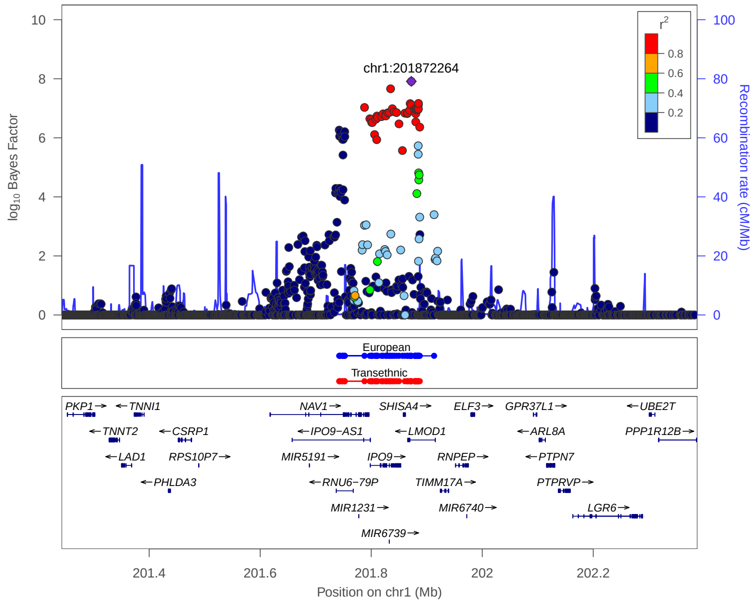


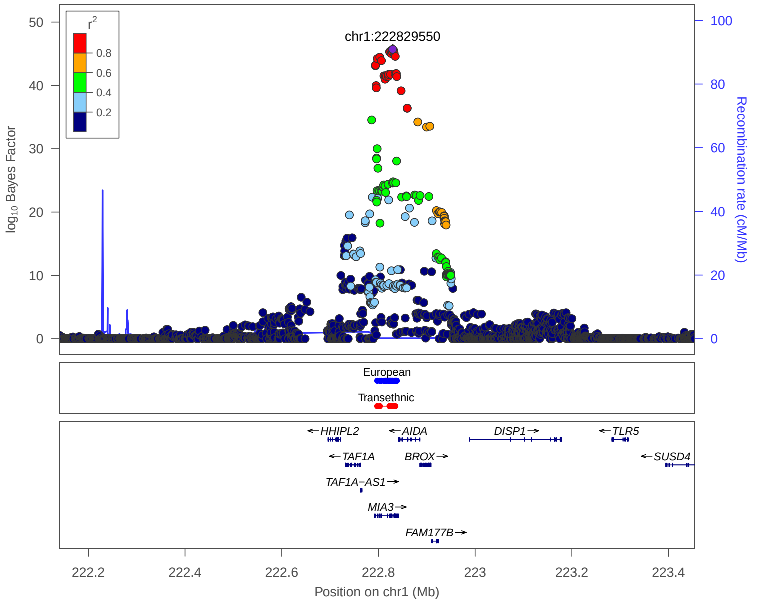


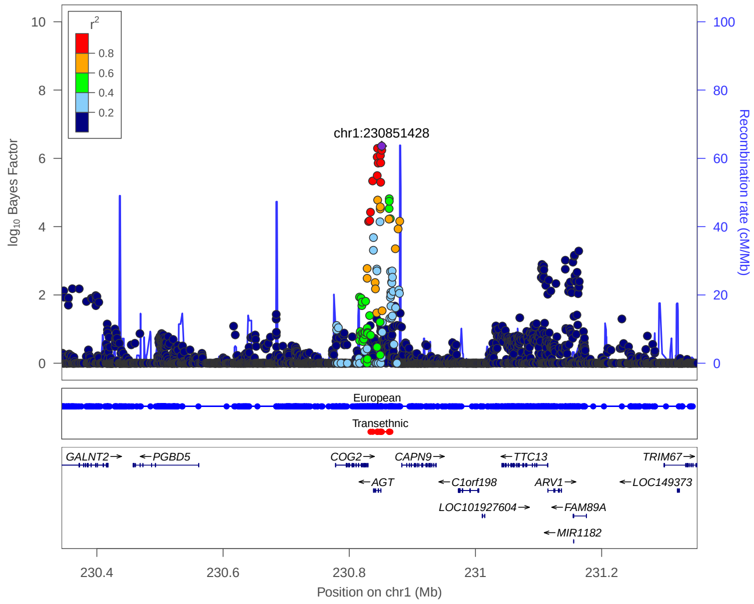


**European:** 99% credible set derived from European meta analysis

**Transethnic:** 99% credible set derived from transethnic meta analysis

### 4. Previously reported 133 loci in Meta analysis

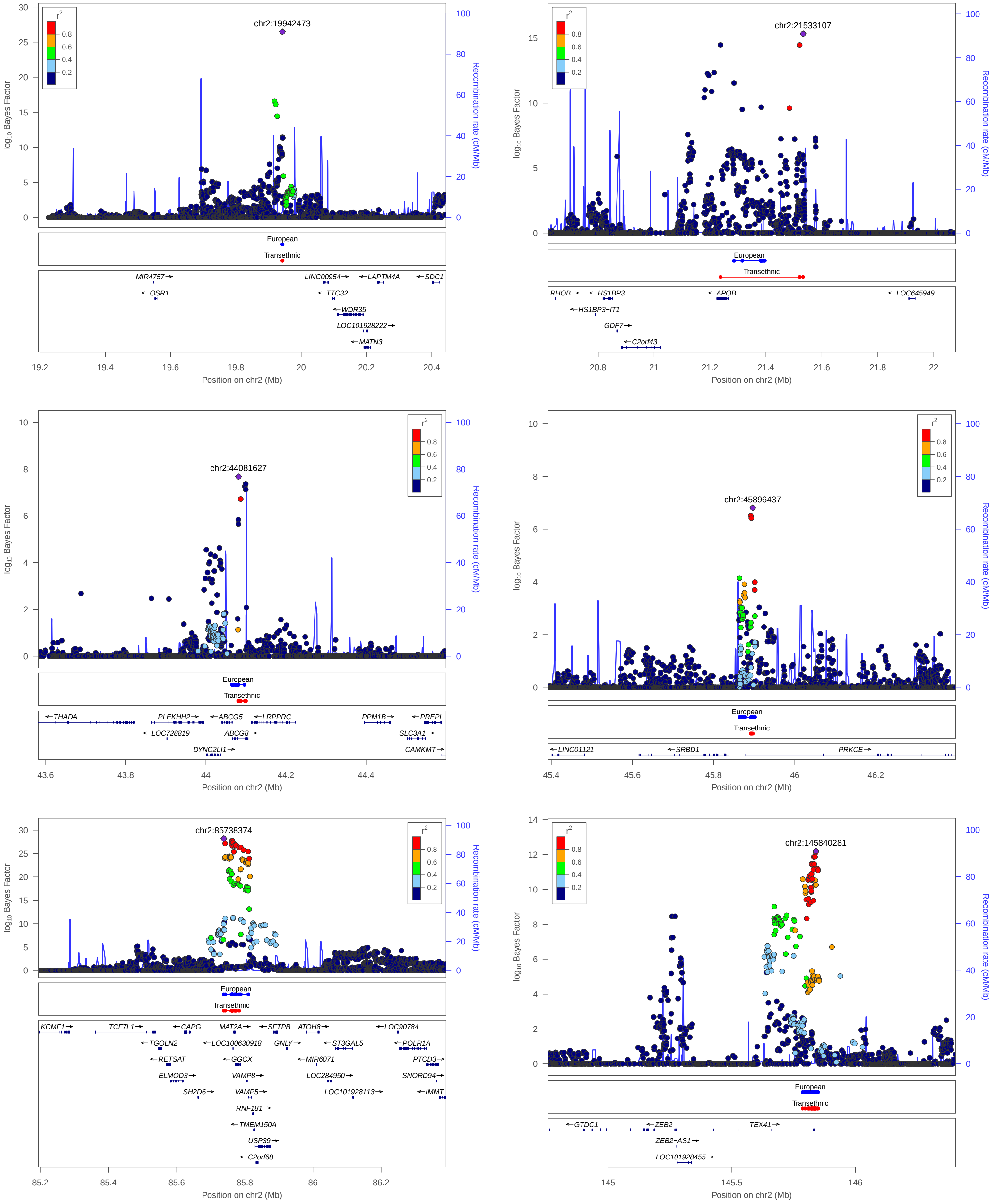


**European:** 99% credible set derived from European meta analysis

**Transethnic:** 99% credible set derived from transethnic meta analysis

### 4. Previously reported 133 loci in Meta analysis

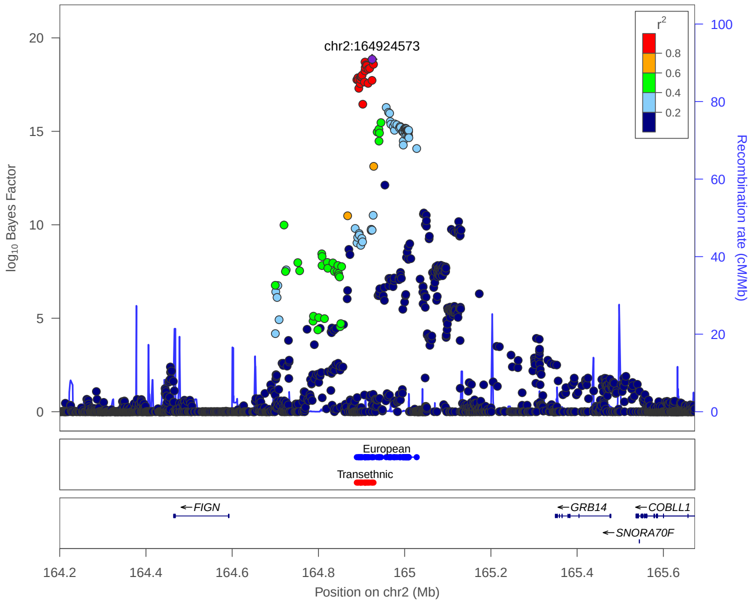


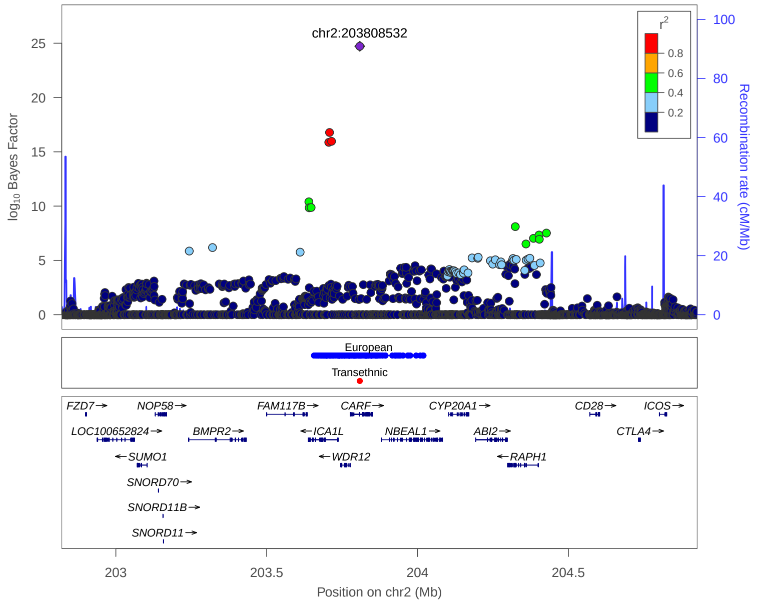


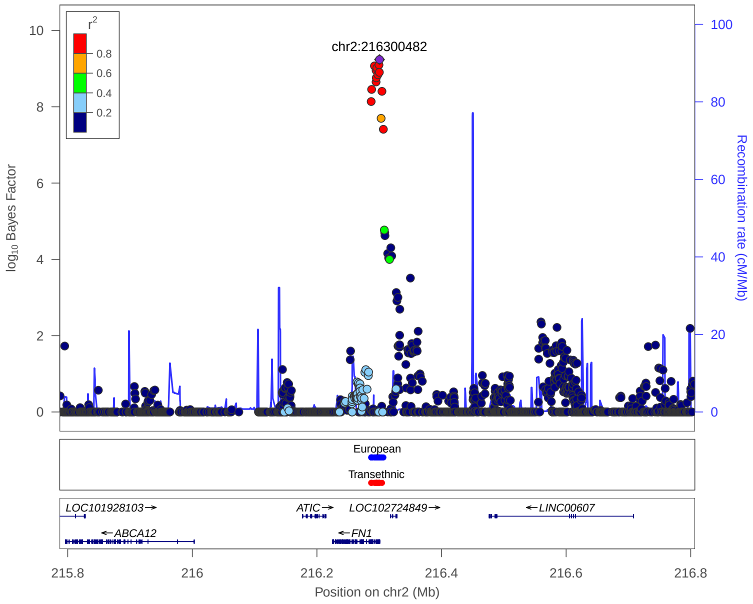


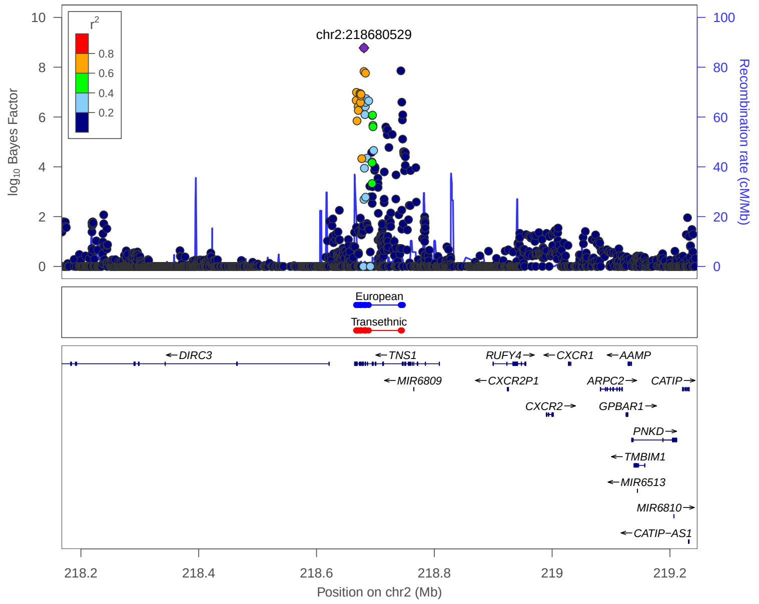


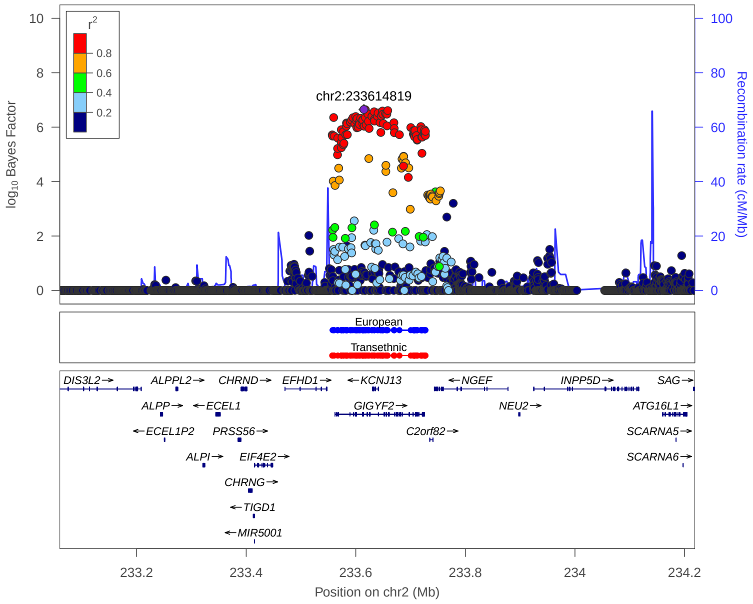


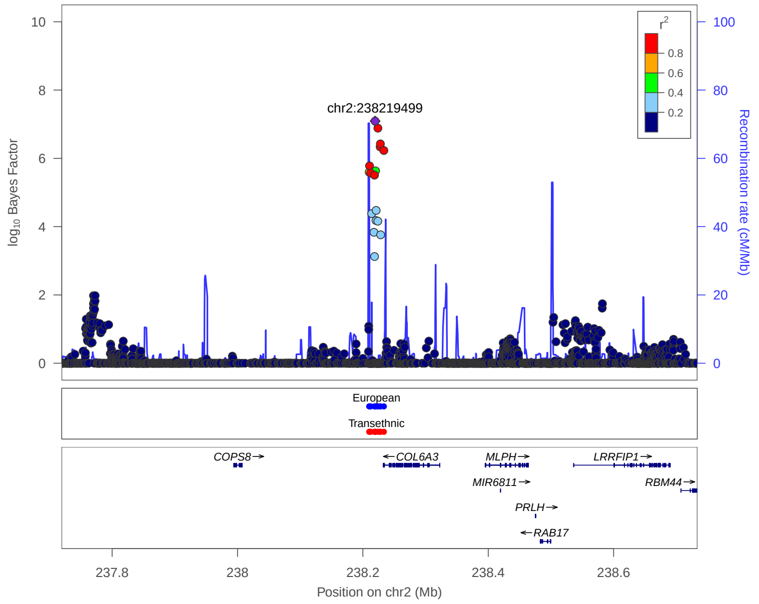


**European:** 99% credible set derived from European meta analysis

**Transethnic:** 99% credible set derived from transethnic meta analysis

### 4. Previously reported 133 loci in Meta analysis

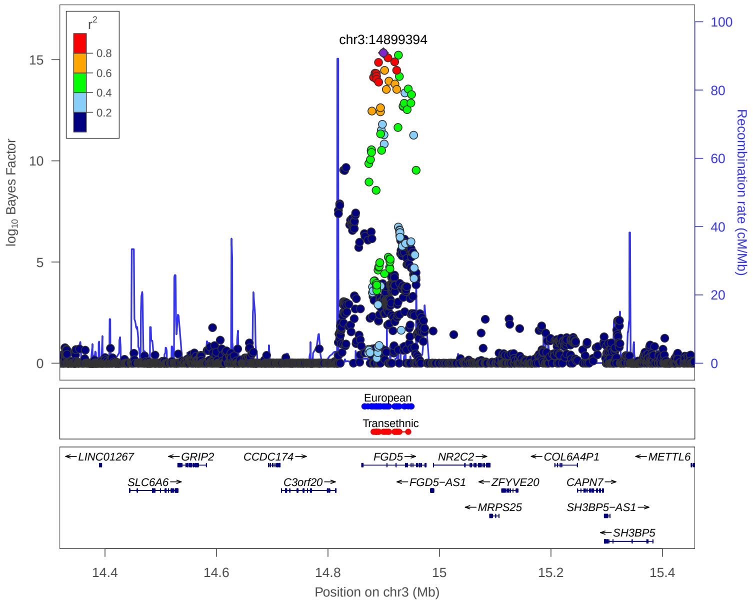


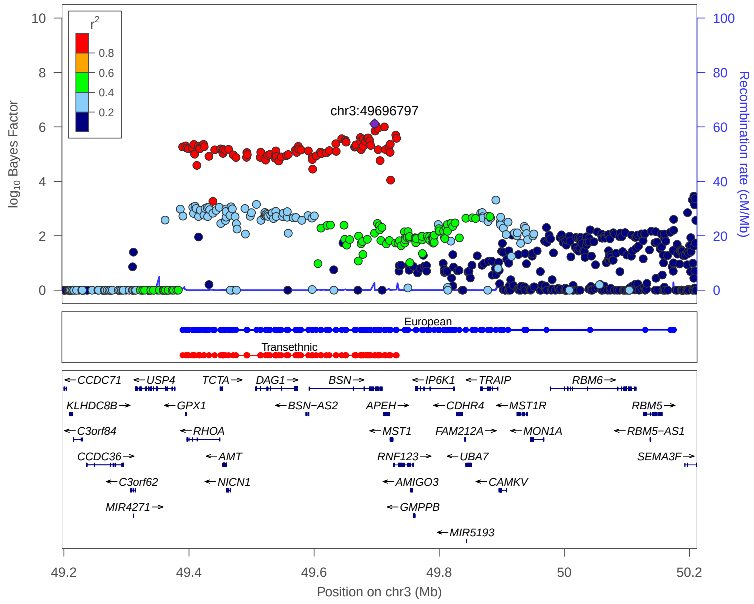


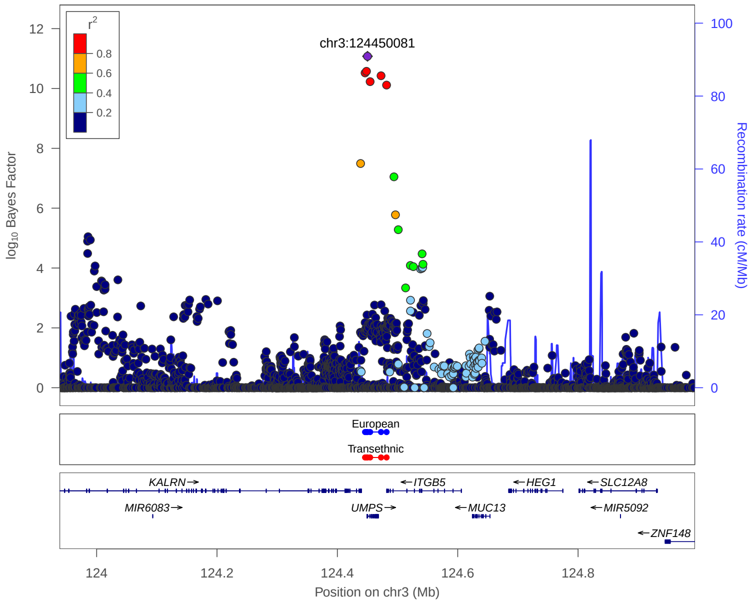


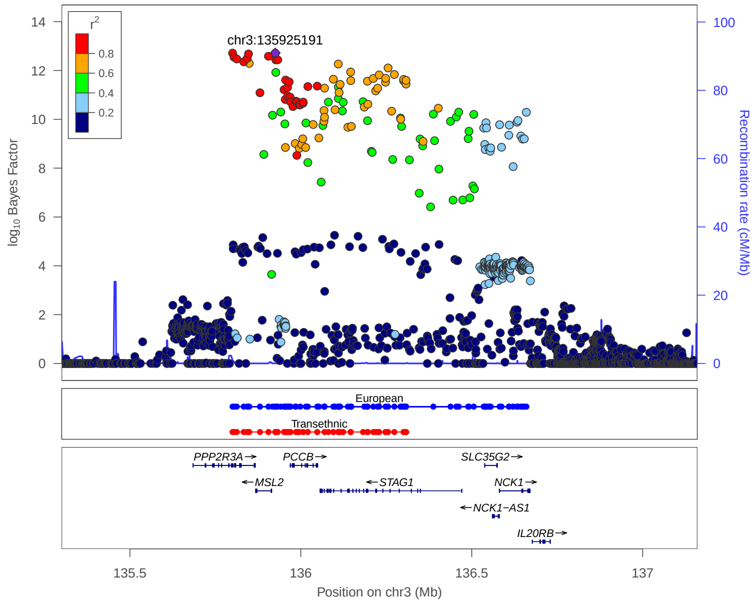


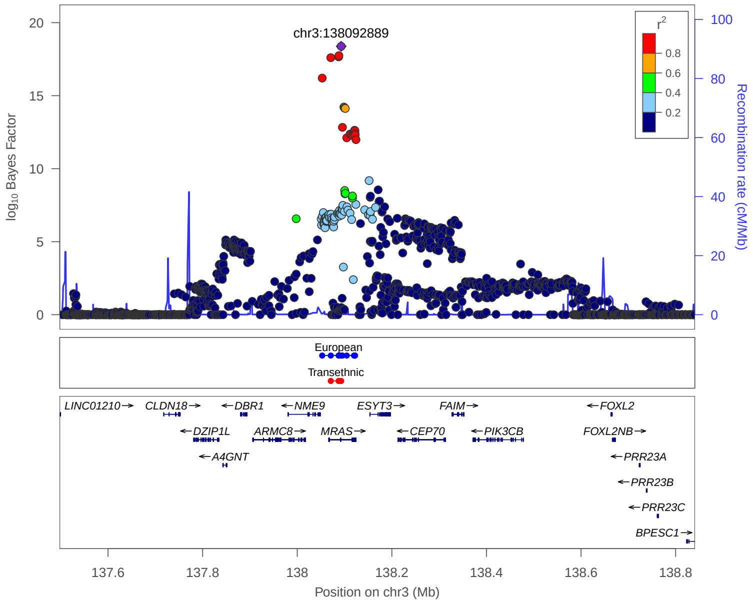


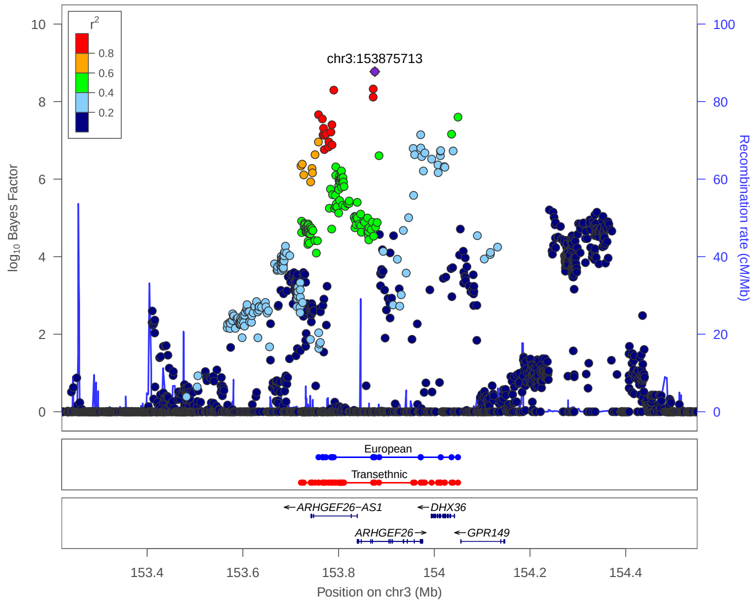


**European:** 99% credible set derived from European meta analysis

**Transethnic:** 99% credible set derived from transethnic meta analysis

### 4. Previously reported 133 loci in Meta analysis

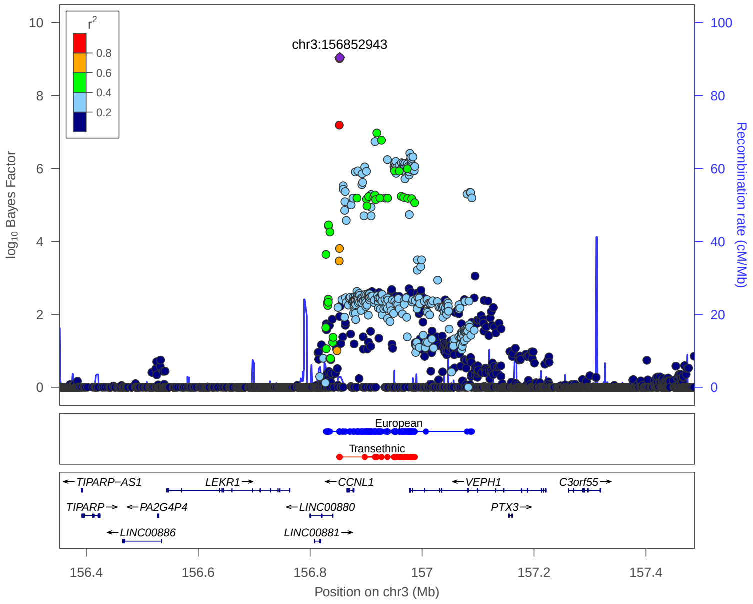


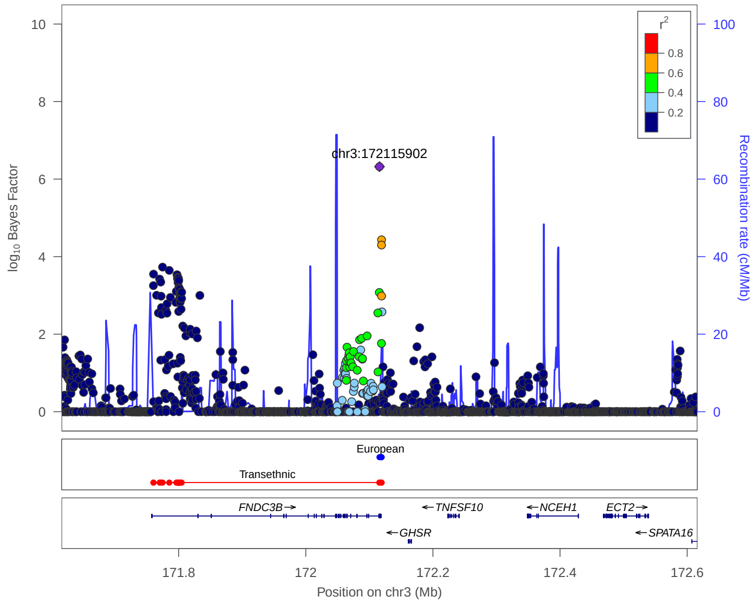


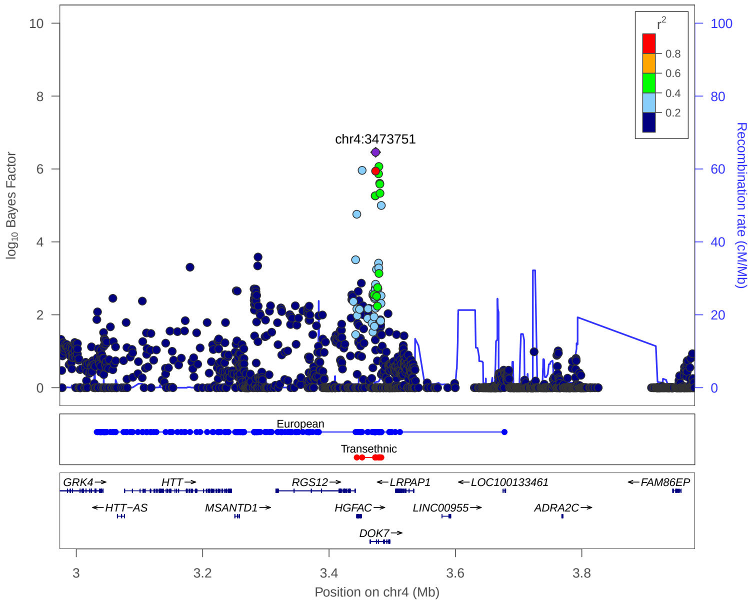


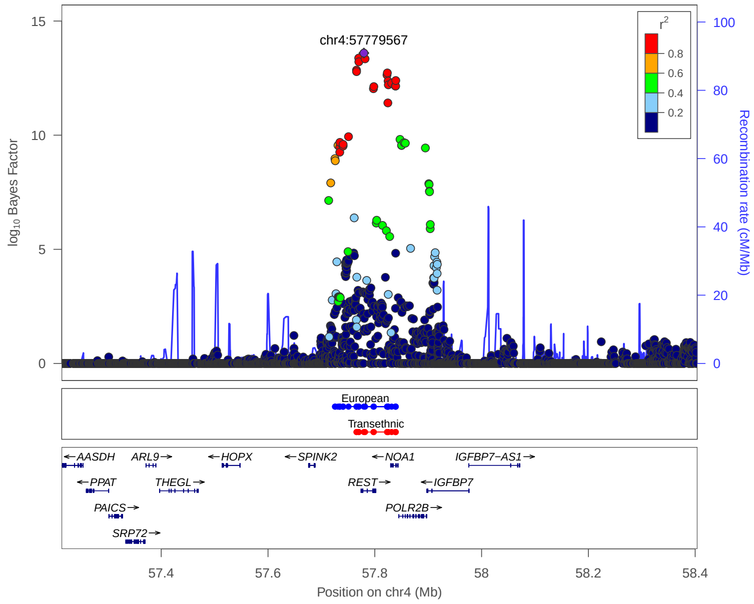


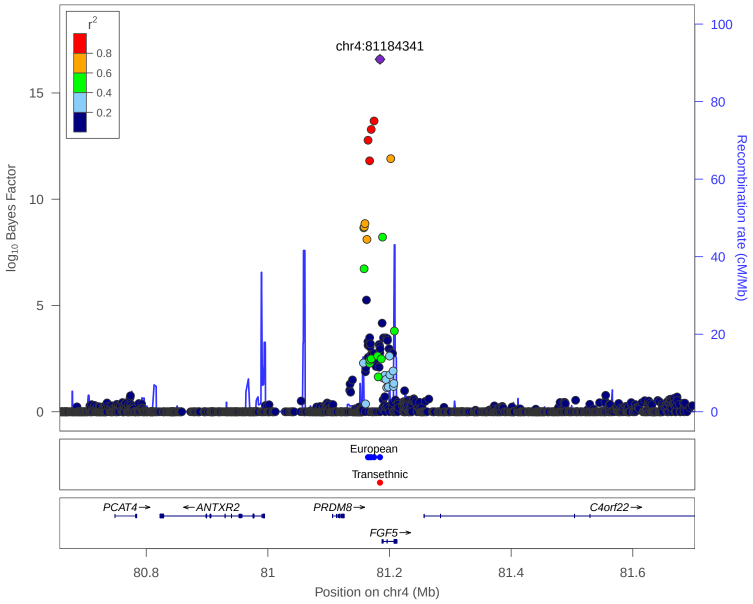


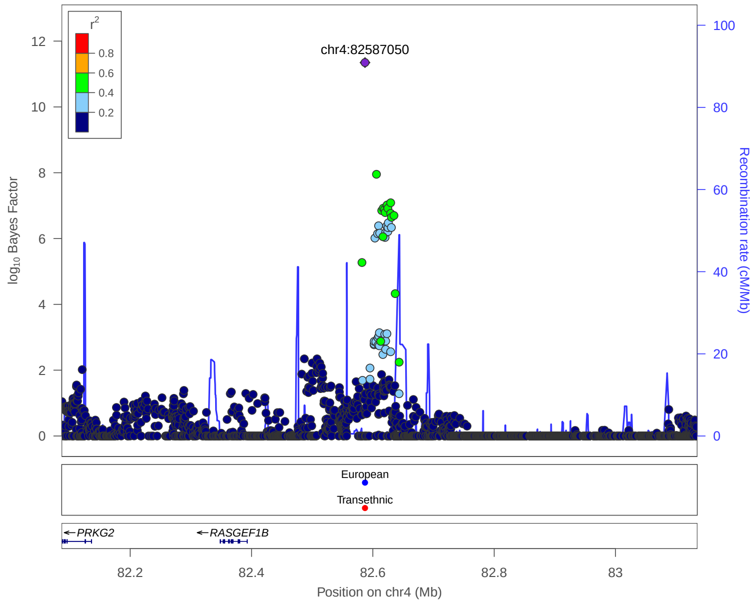


**European:** 99% credible set derived from European meta analysis
**Transethnic:** 99% credible set derived from transethnic meta analysis

### 4. Previously reported 133 loci in Meta analysis

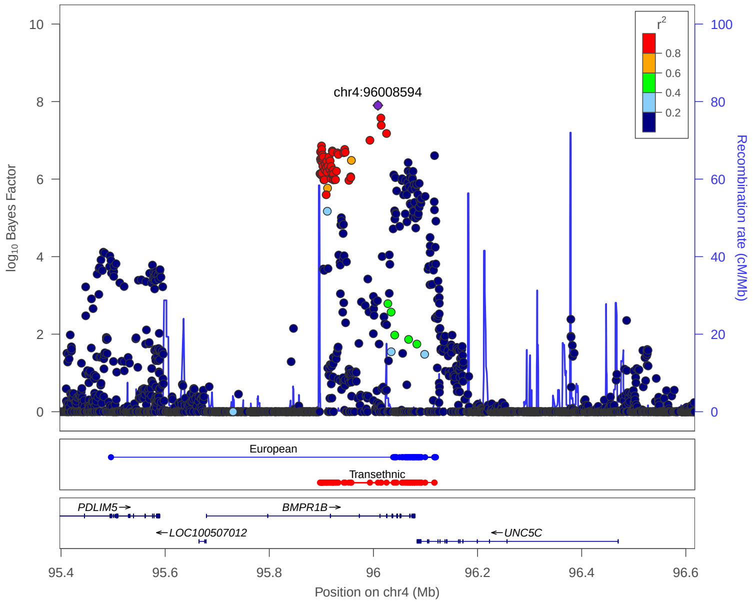


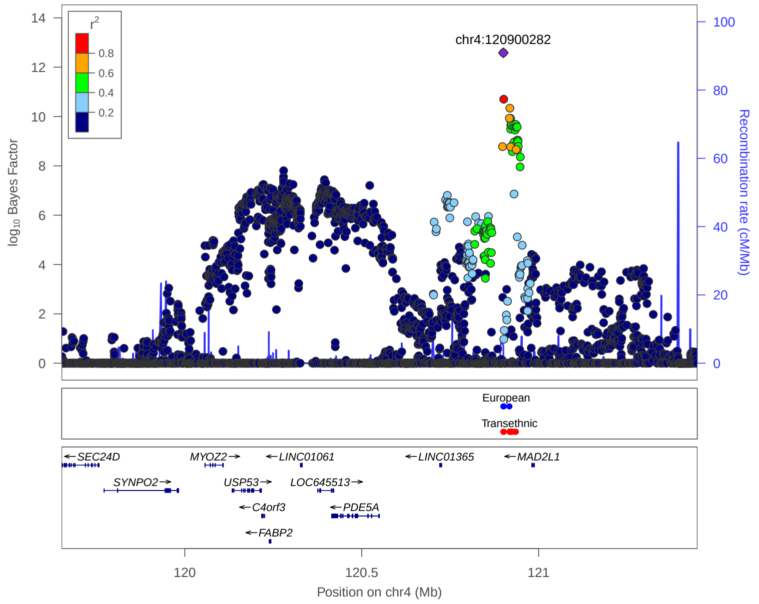


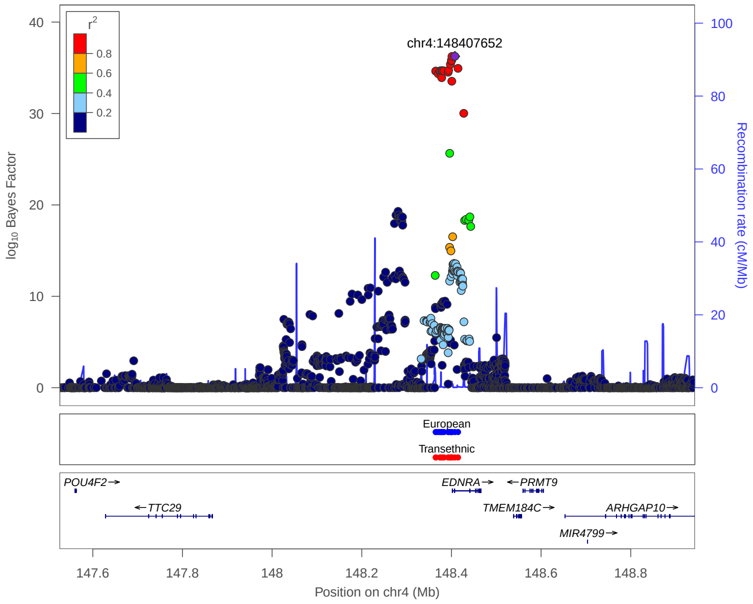


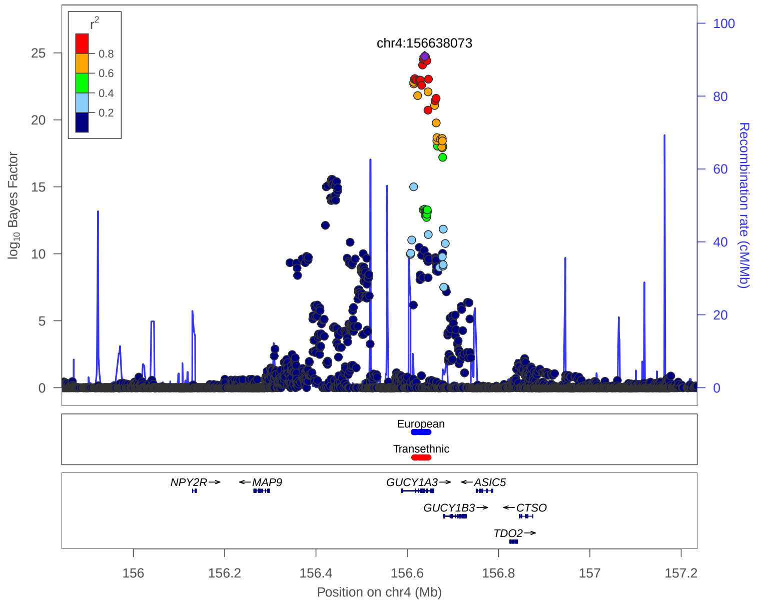


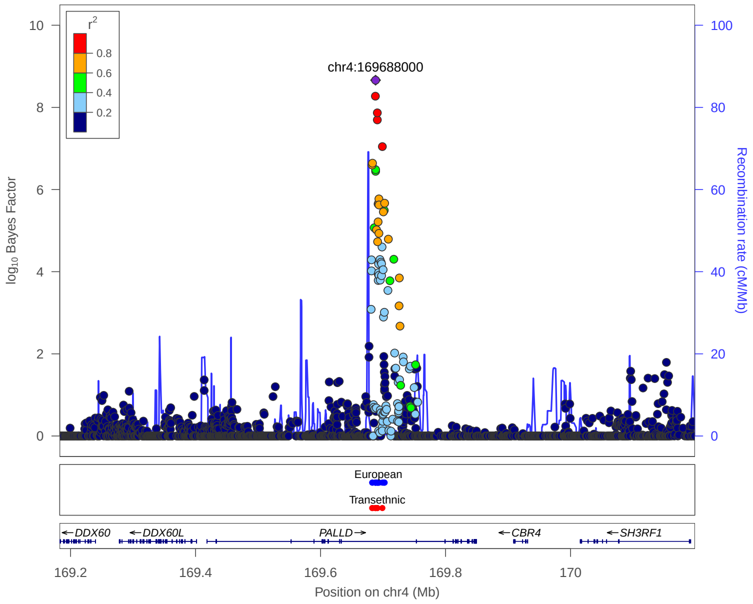


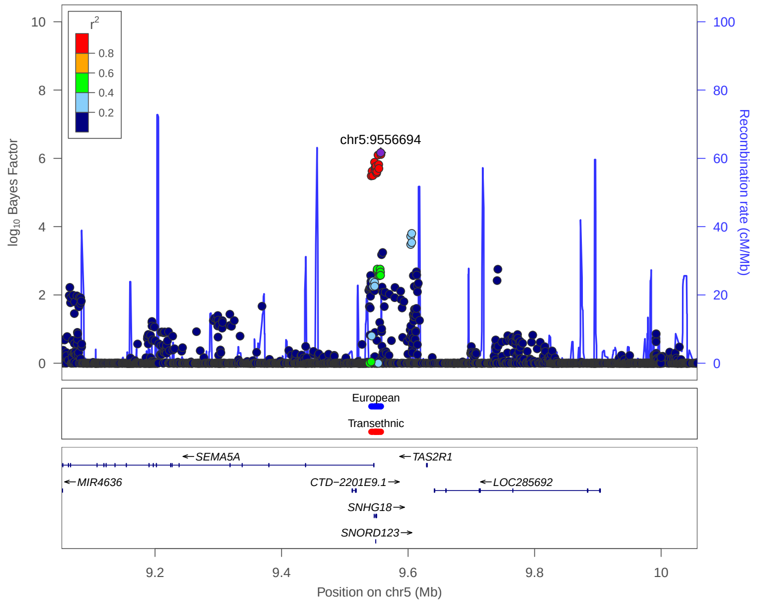


**European:** 99% credible set derived from European meta analysis

**Transethnic:** 99% credible set derived from transethnic meta analysis

### 4. Previously reported 133 loci in Meta analysis

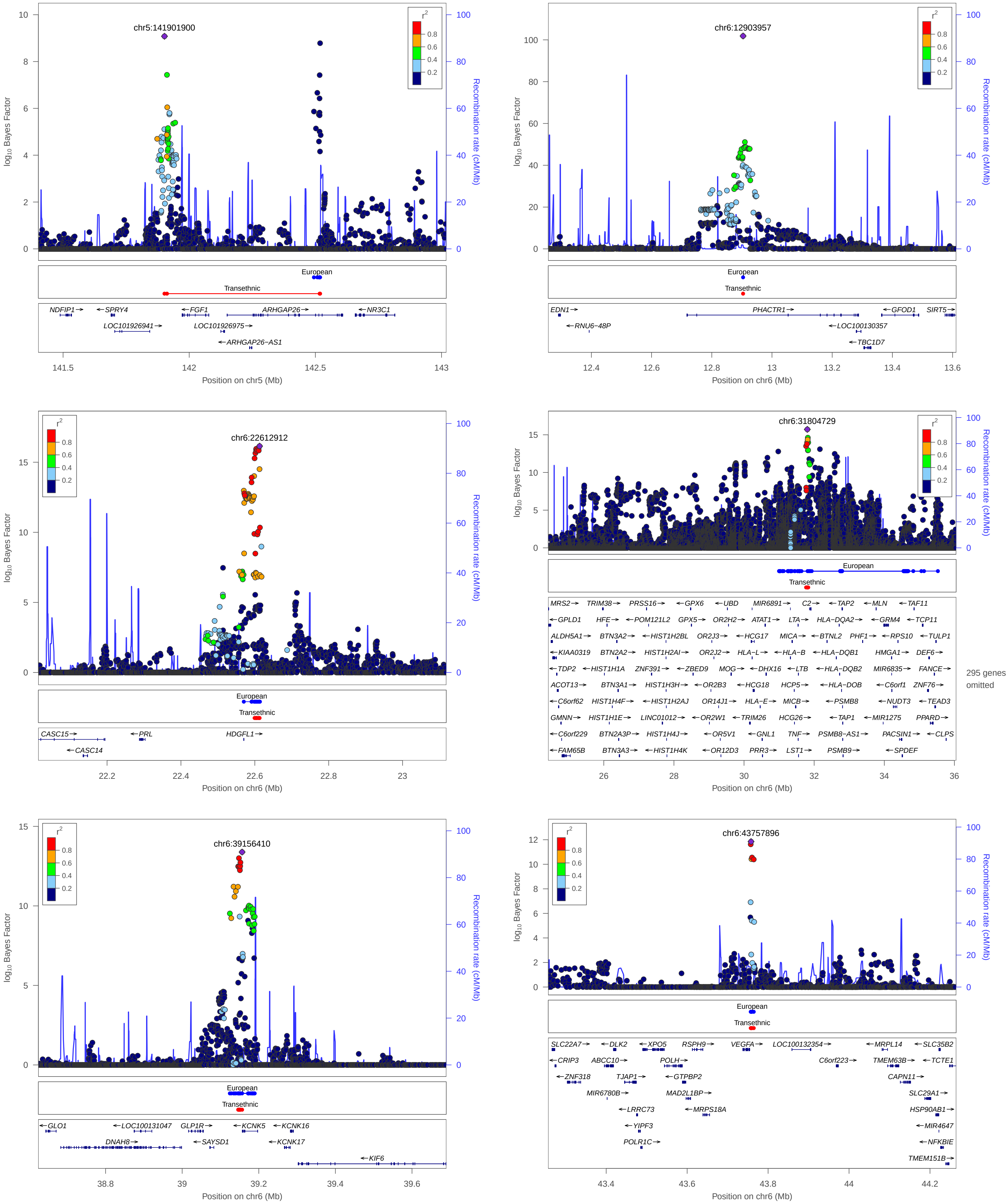


**European:** 99% credible set derived from European meta analysis

**Transethnic:** 99% credible set derived from transethnic meta analysis

### 4. Previously reported 133 loci in Meta analysis

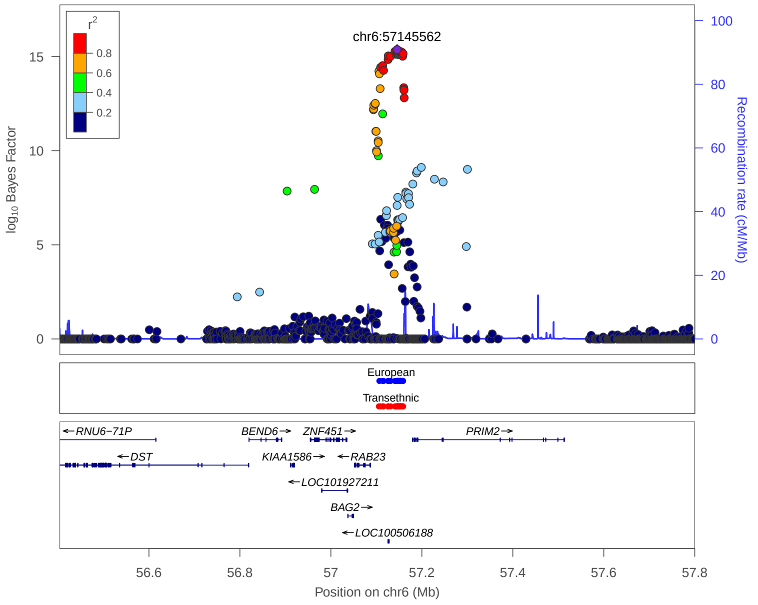


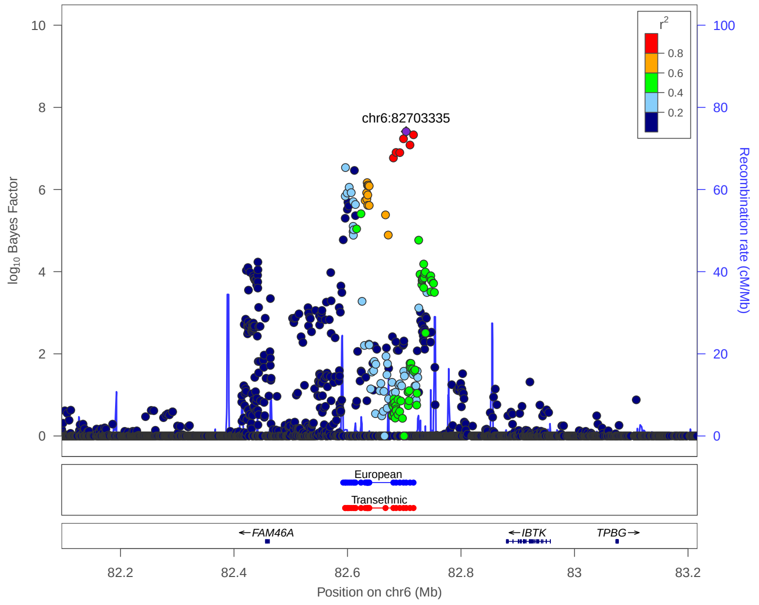


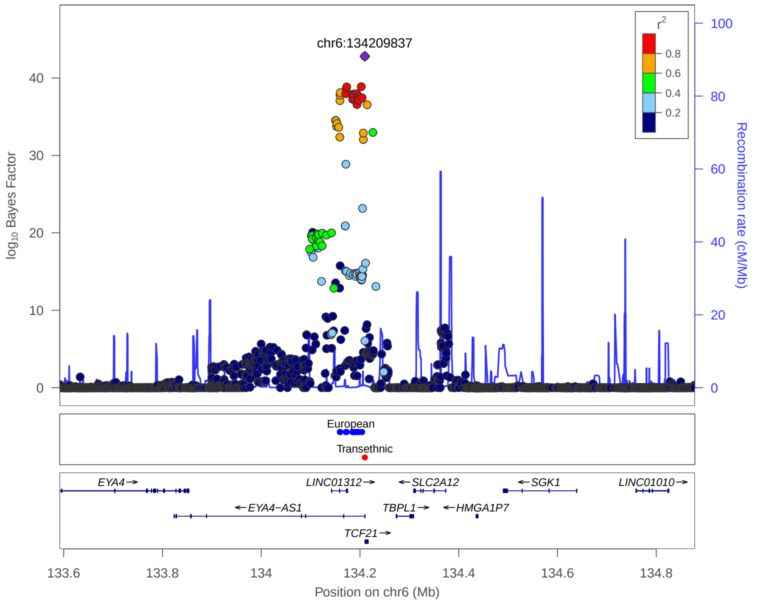


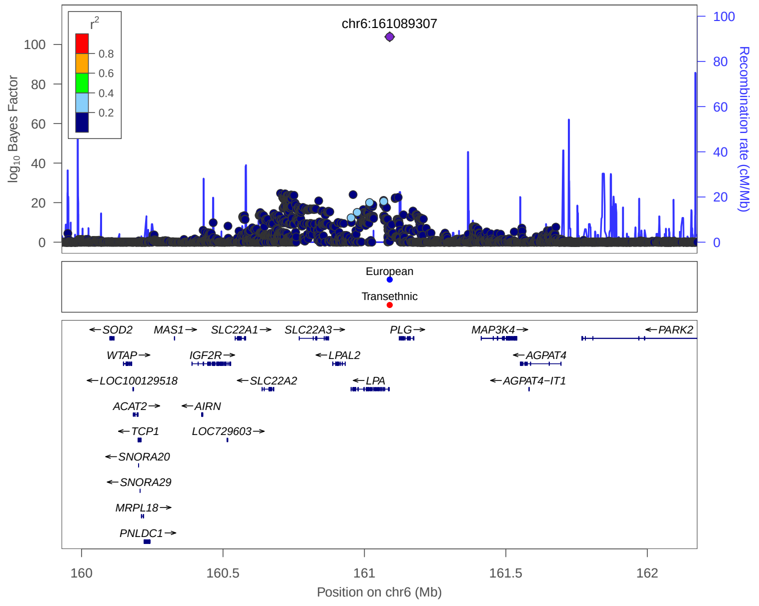


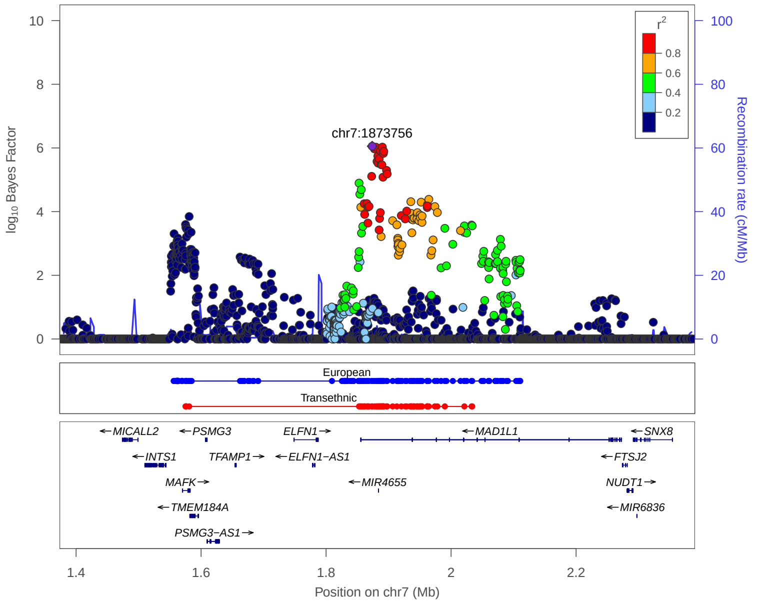


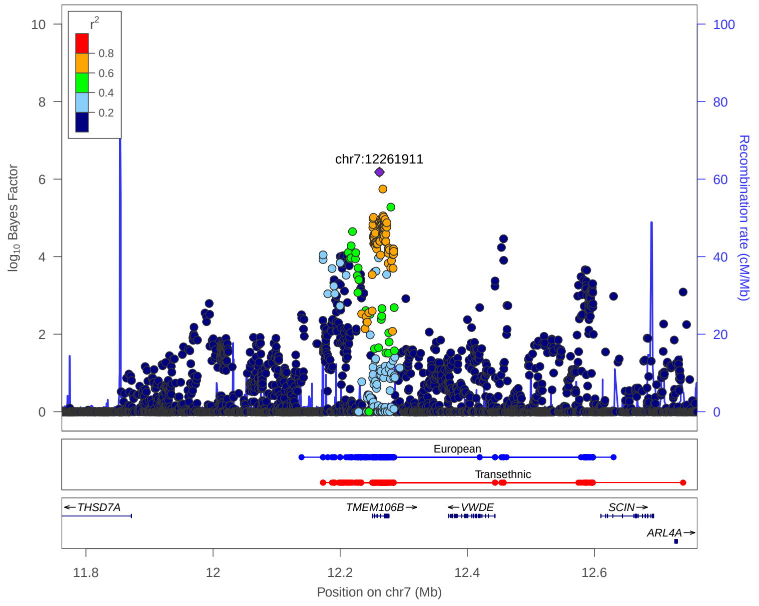


**European:** 99% credible set derived from European meta analysis

**Transethnic:** 99% credible set derived from transethnic meta analysis

### 4. Previously reported 133 loci in Meta analysis

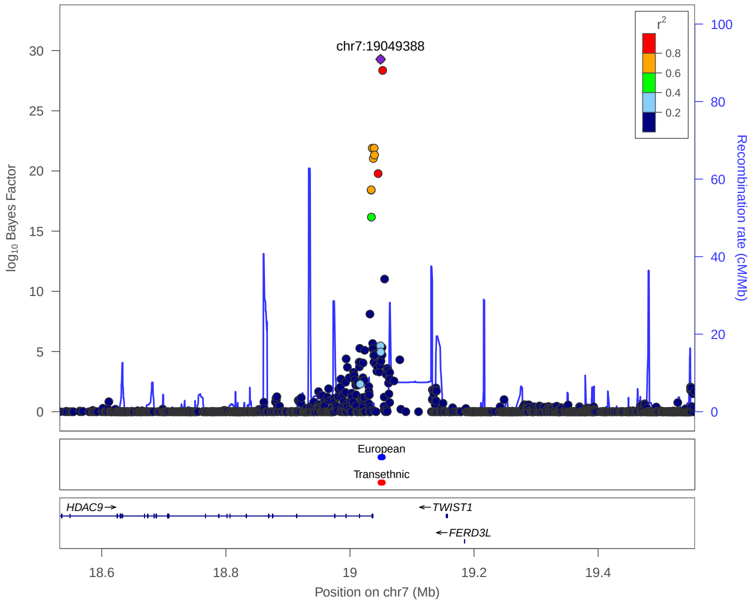


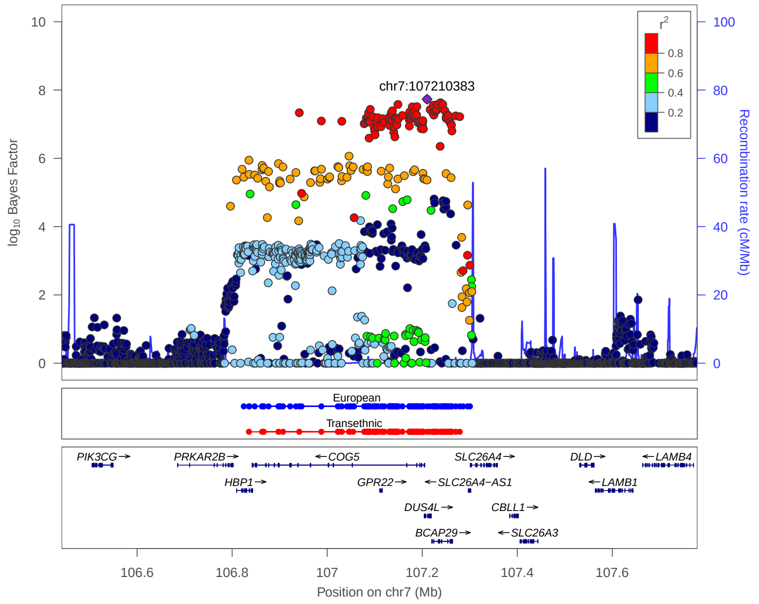


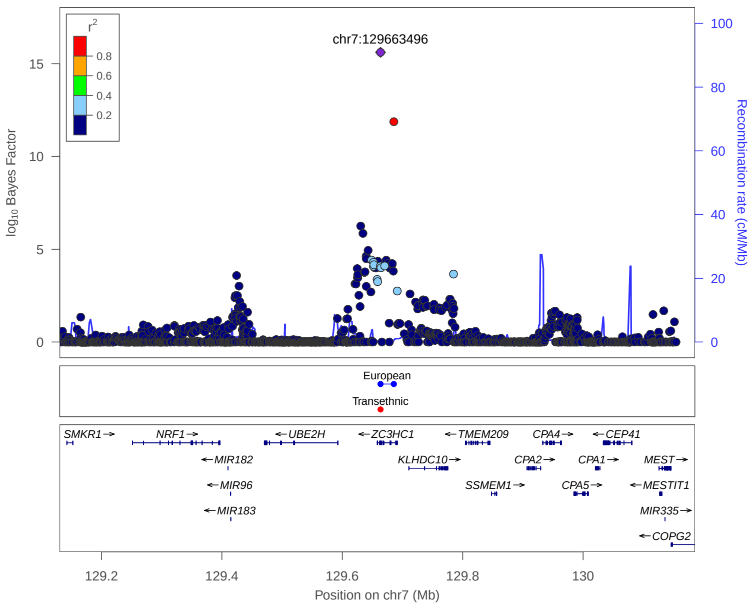


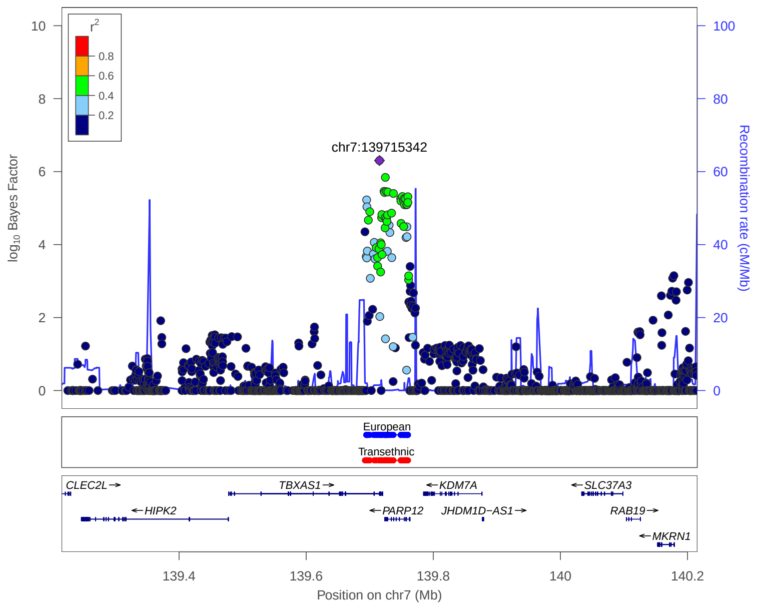


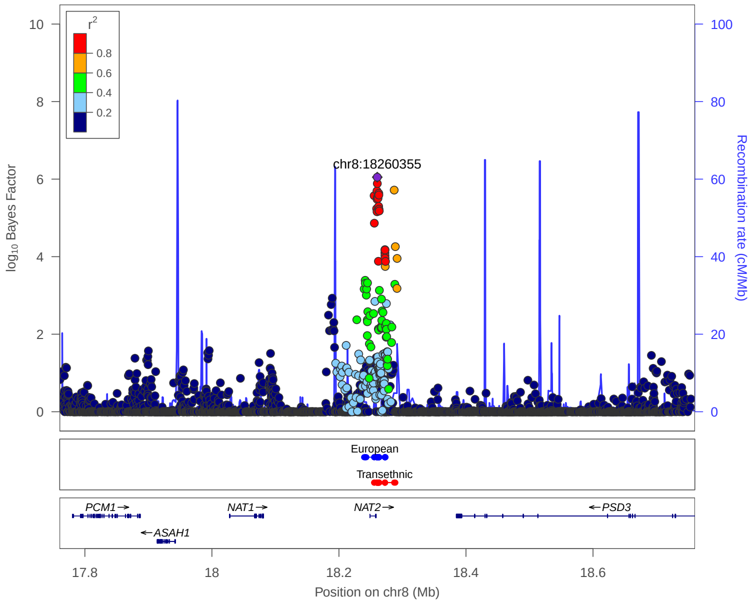


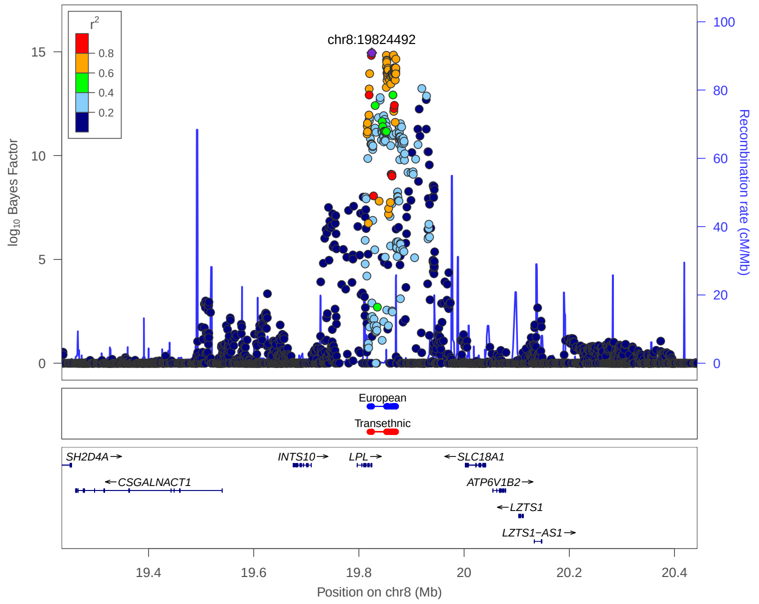


**European:** 99% credible set derived from European meta analysis

**Transethnic:** 99% credible set derived from transethnic meta analysis

### 4. Previously reported 133 loci in Meta analysis

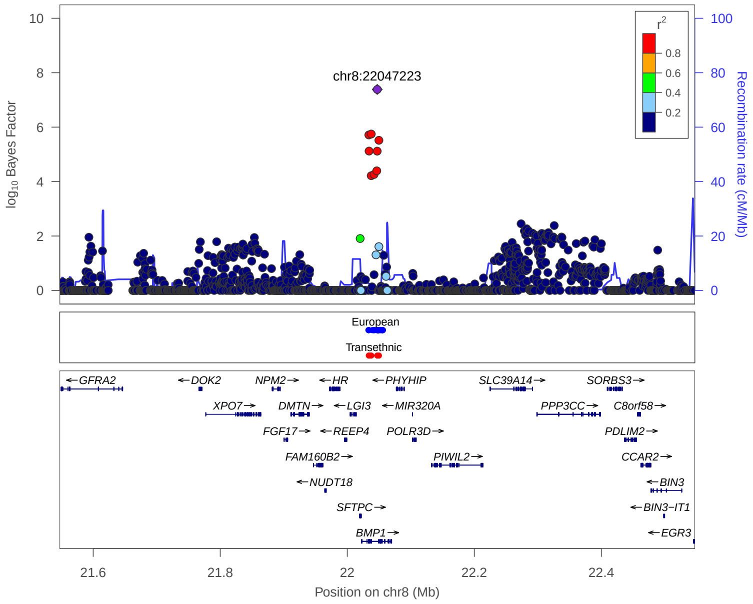


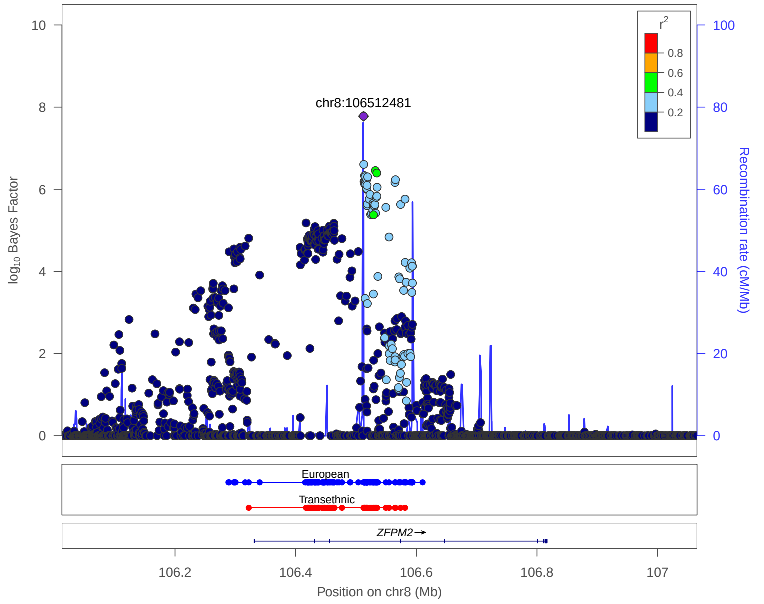


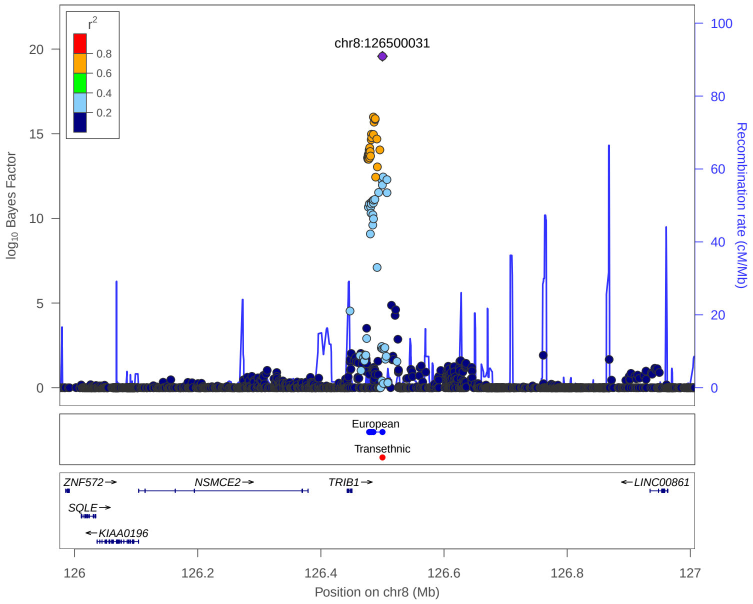


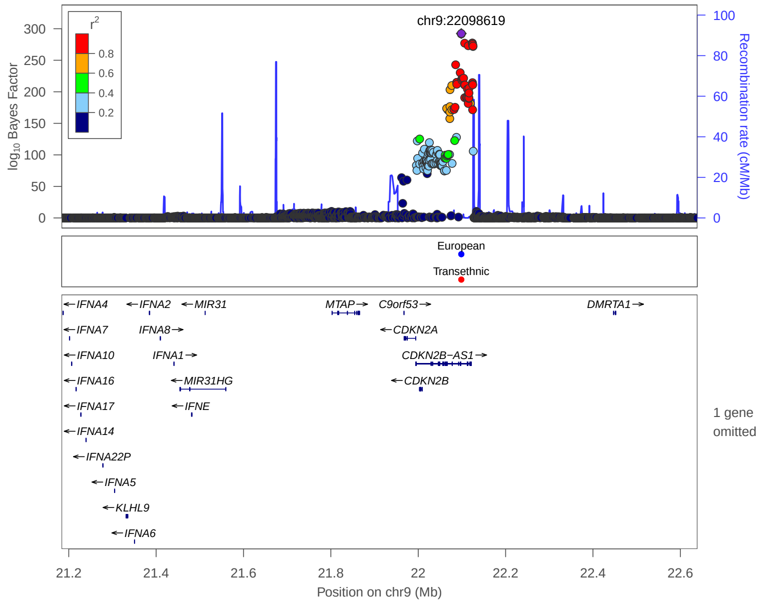


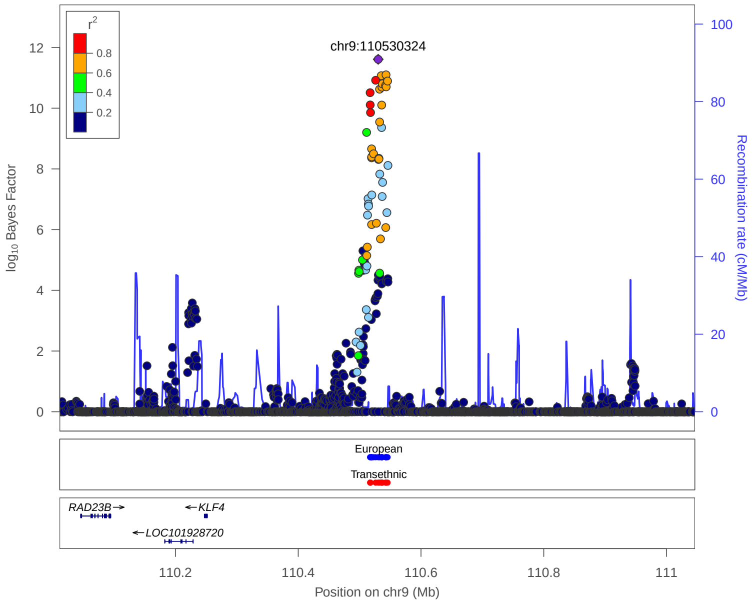


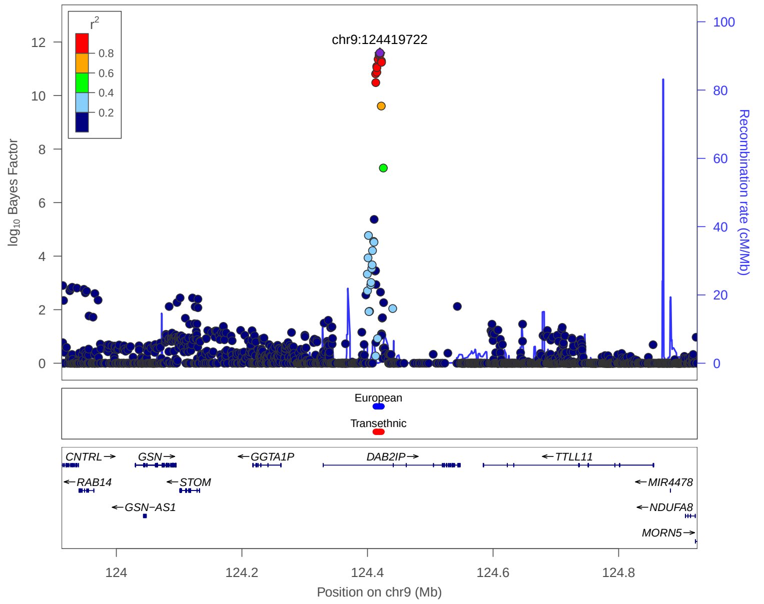


**European:** 99% credible set derived from European meta analysis

**Transethnic:** 99% credible set derived from transethnic meta analysis

### 4. Previously reported 133 loci in Meta analysis

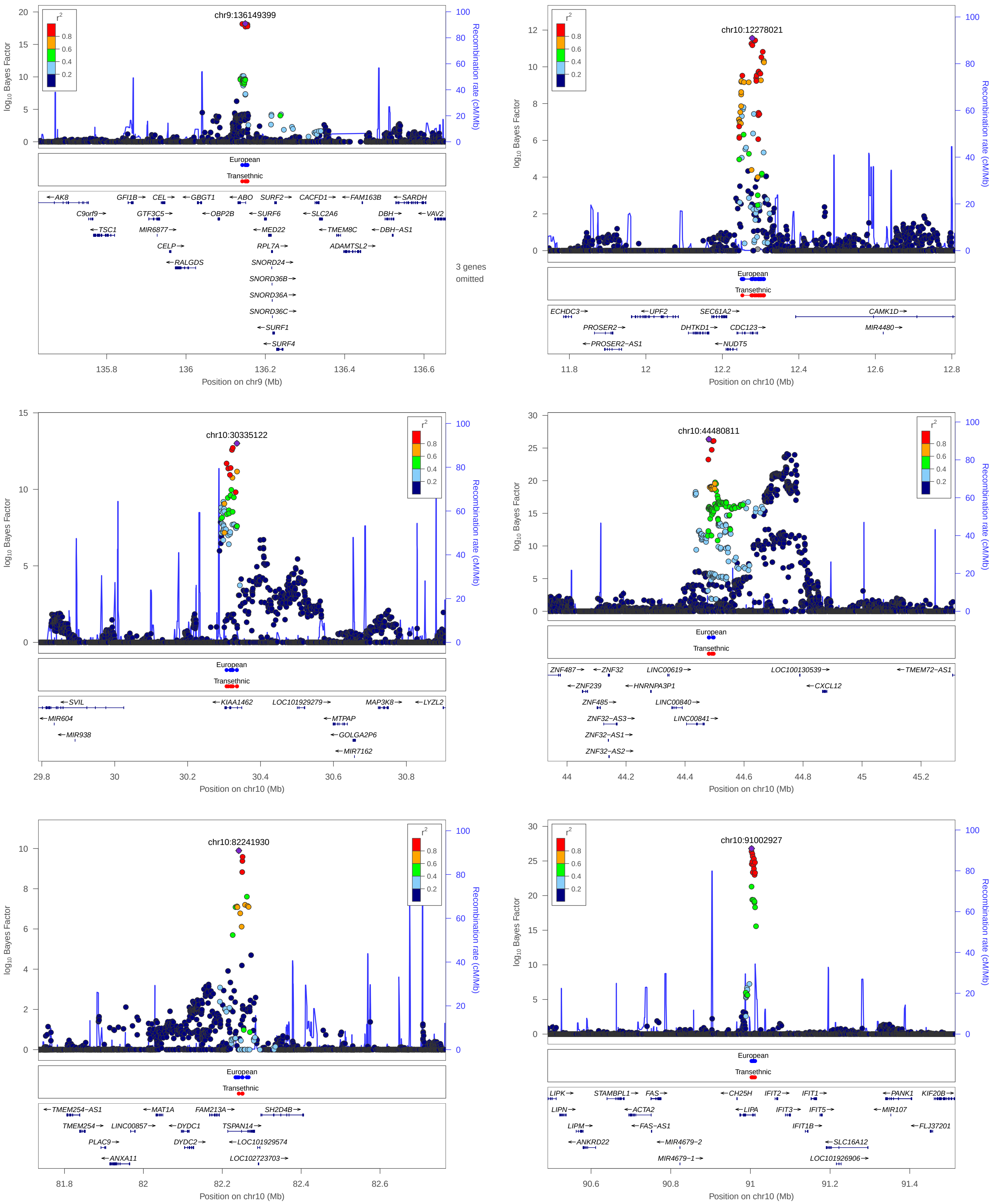


**European:** 99% credible set derived from European meta analysis

**Transethnic:** 99% credible set derived from transethnic meta analysis

### 4. Previously reported 133 loci in Meta analysis

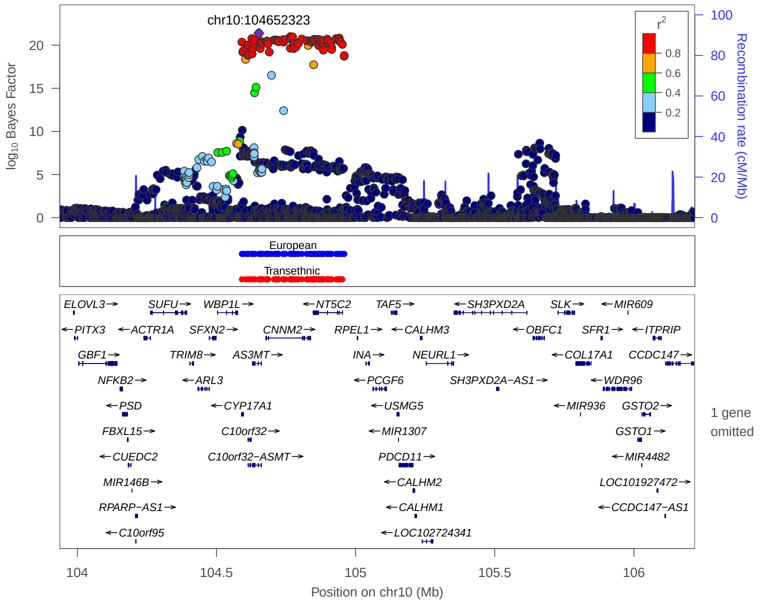


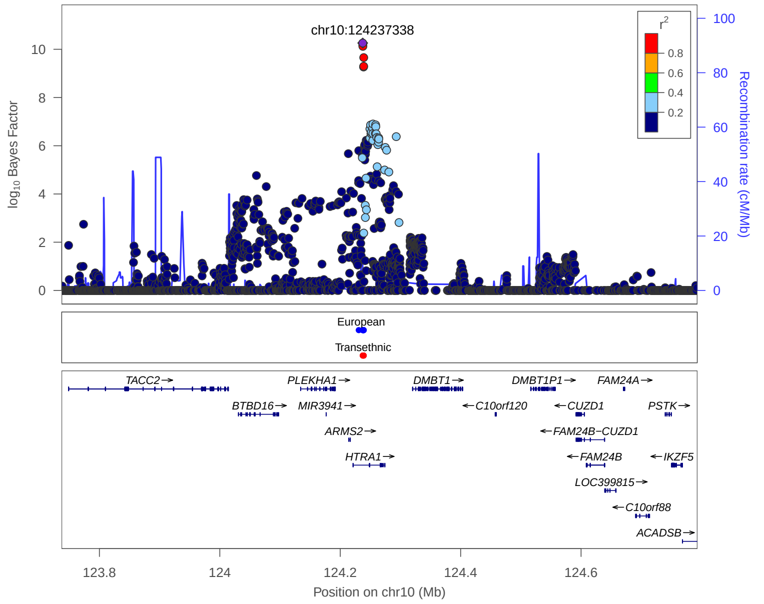


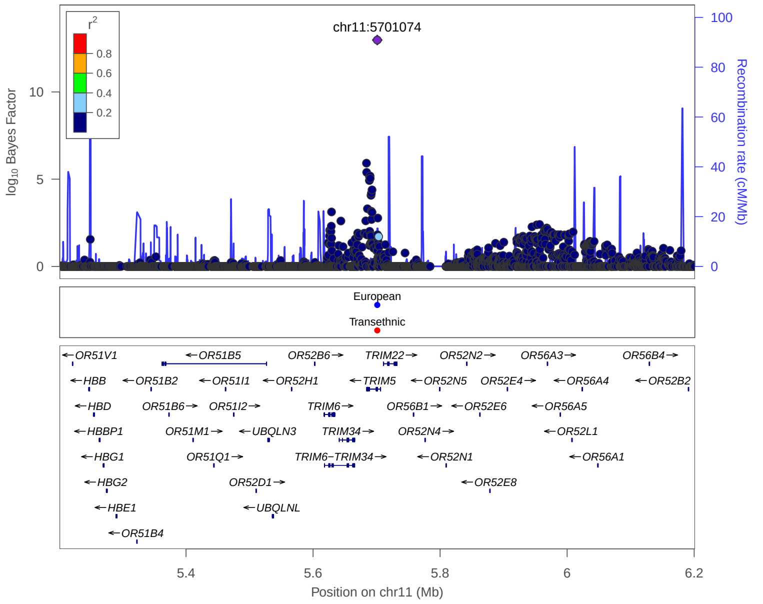


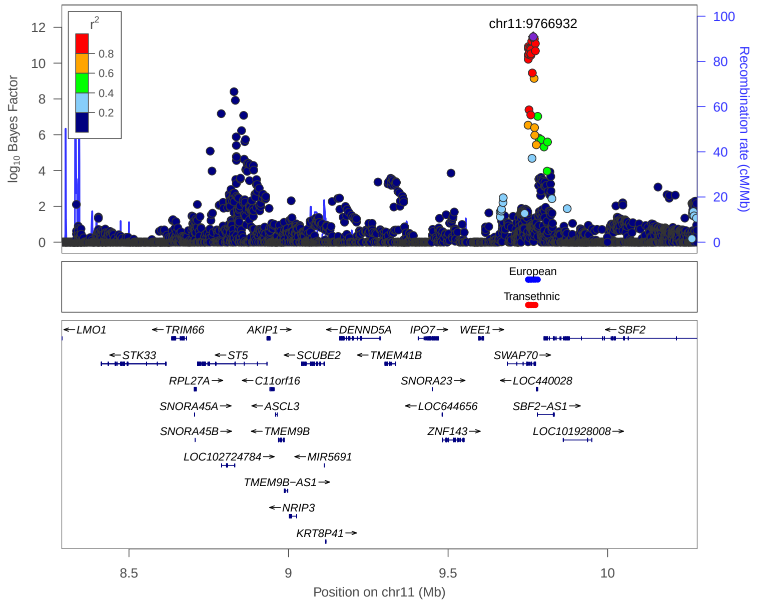


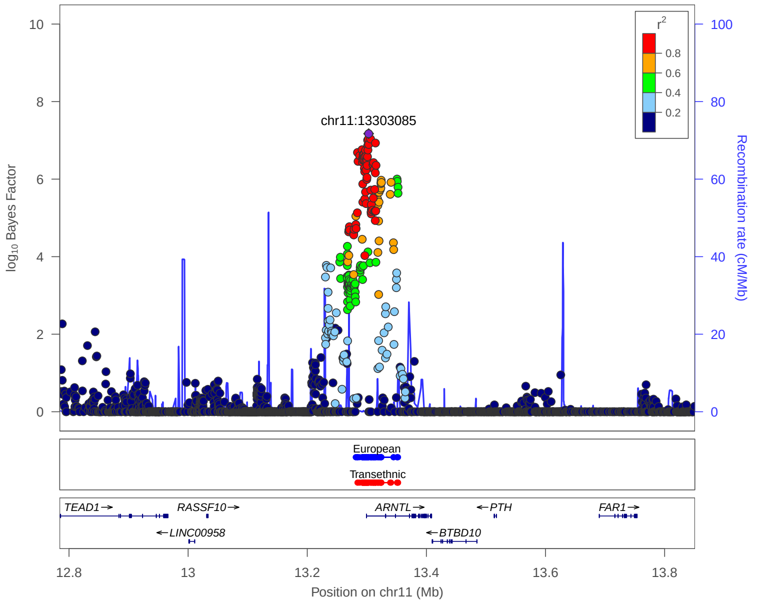


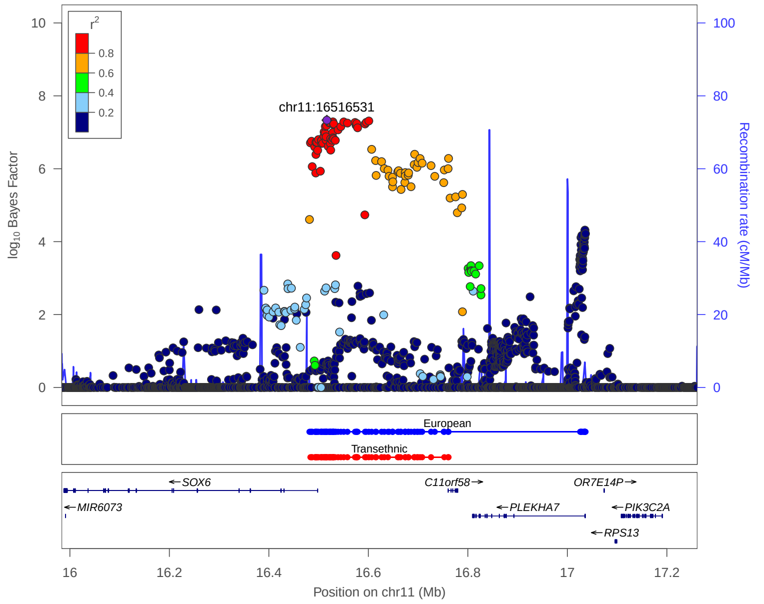


European: 99% credible set derived from European meta analysis

Transethnic: 99% credible set derived from transethnic meta analysis

### 4. Previously reported 133 loci in Meta analysis

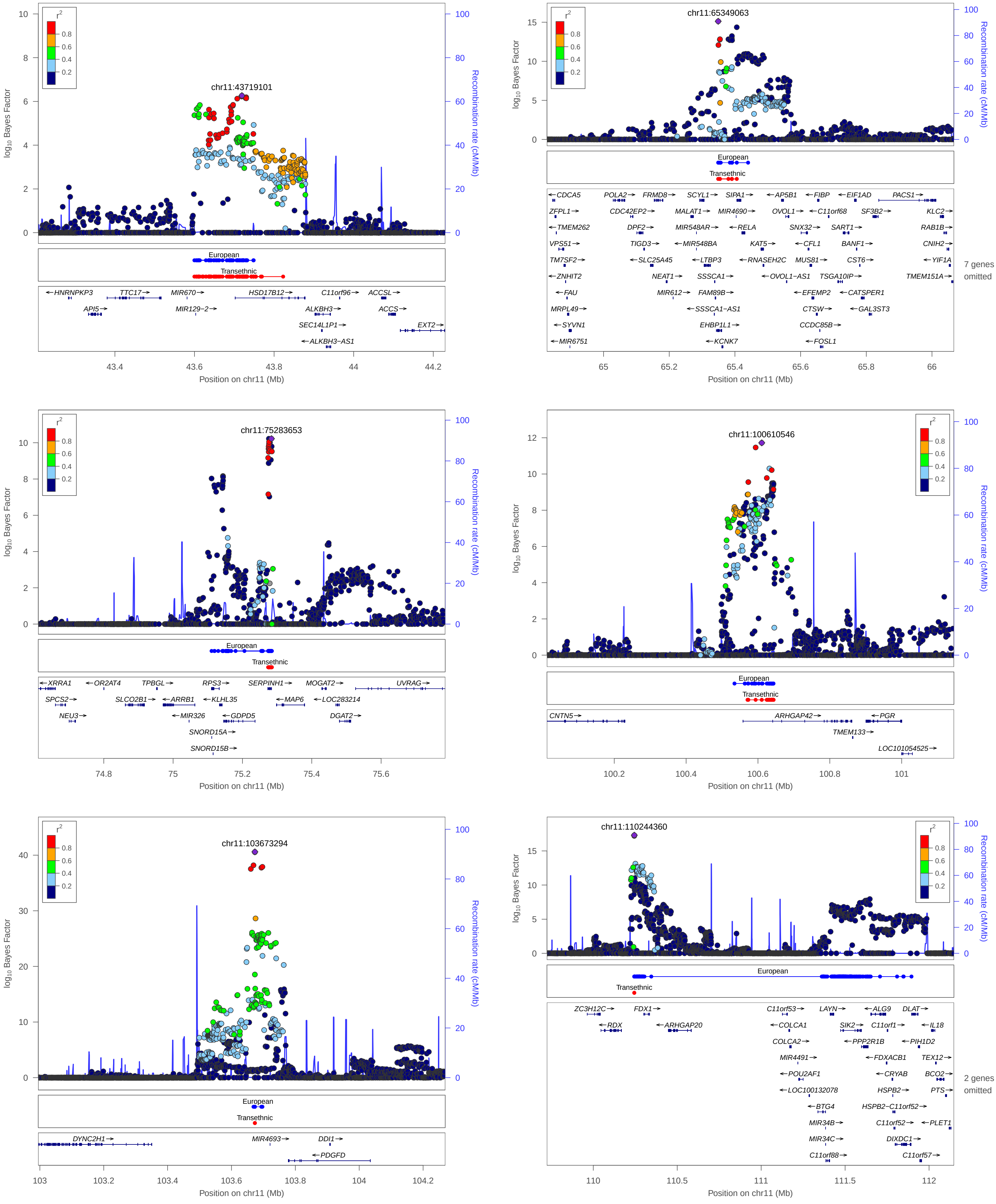


European: 99% credible set derived from European meta analysis

Transethnic: 99% credible set derived from transethnic meta analysis

### 4. Previously reported 133 loci in Meta analysis

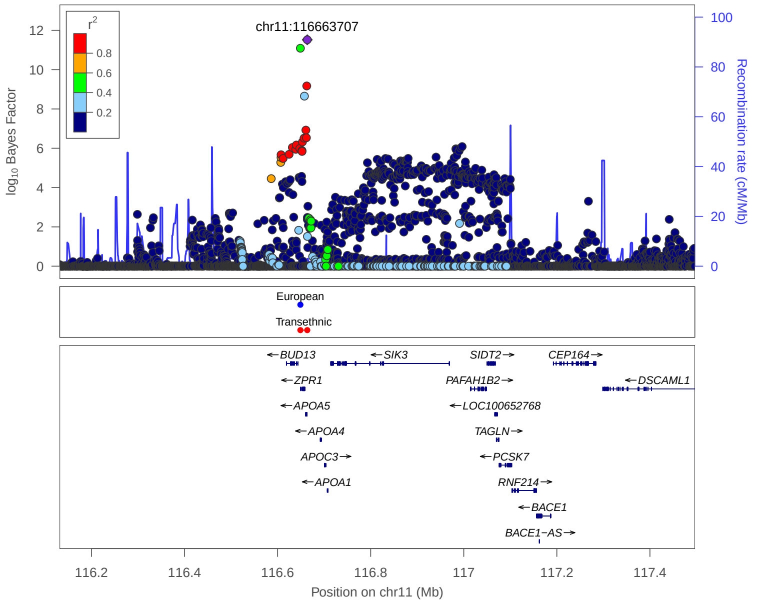


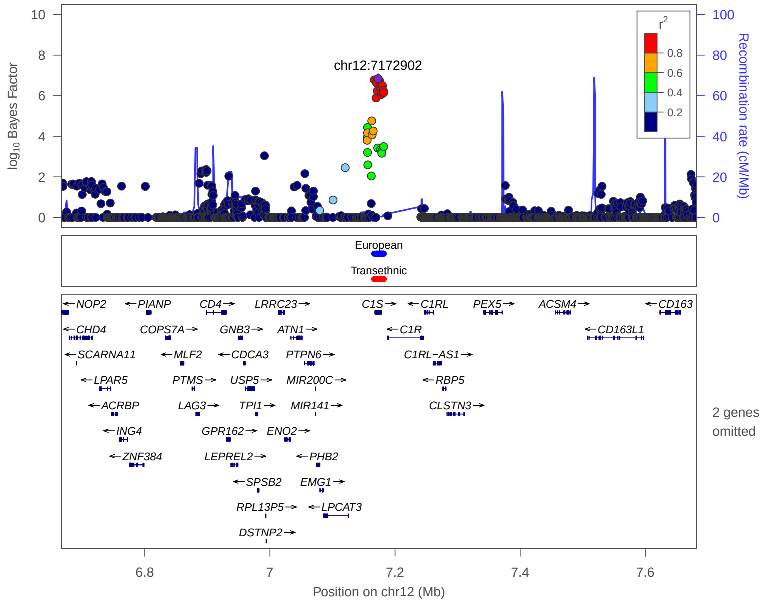


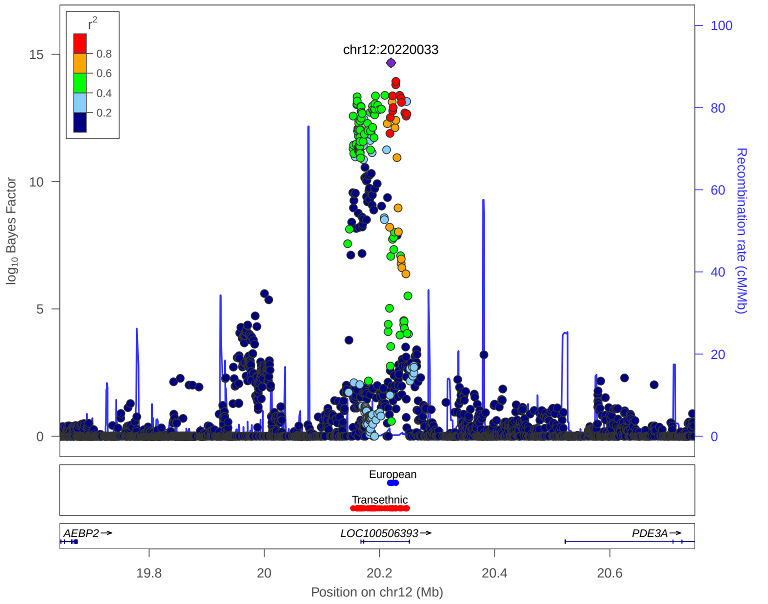


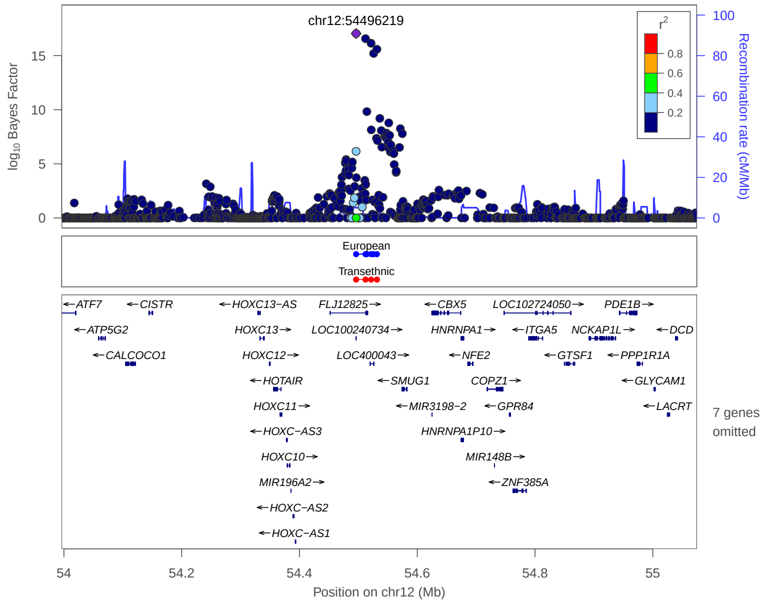


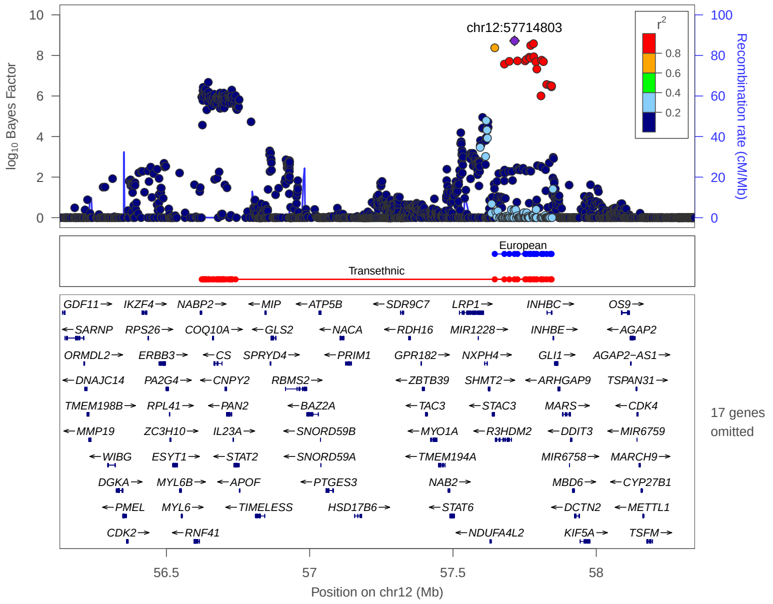


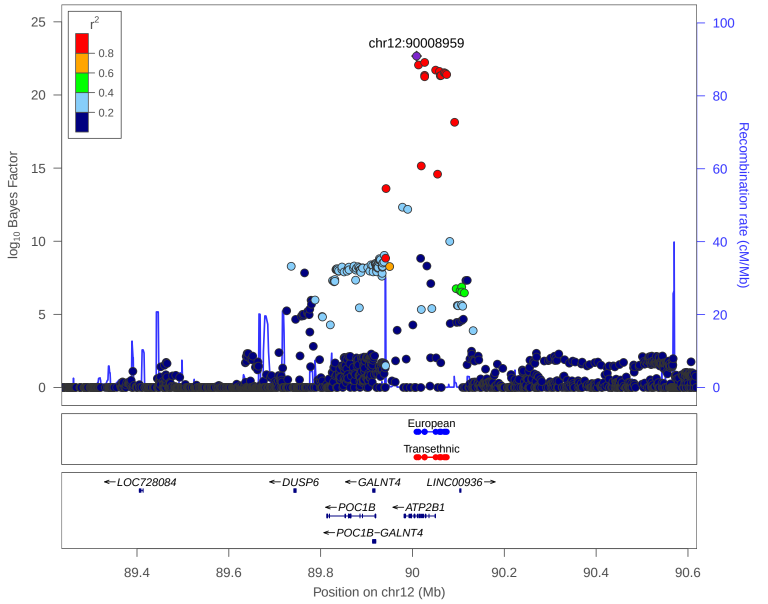


**European:** 99% credible set derived from European meta analysis

**Transethnic:** 99% credible set derived from transethnic meta analysis

### 4. Previously reported 133 loci in Meta analysis

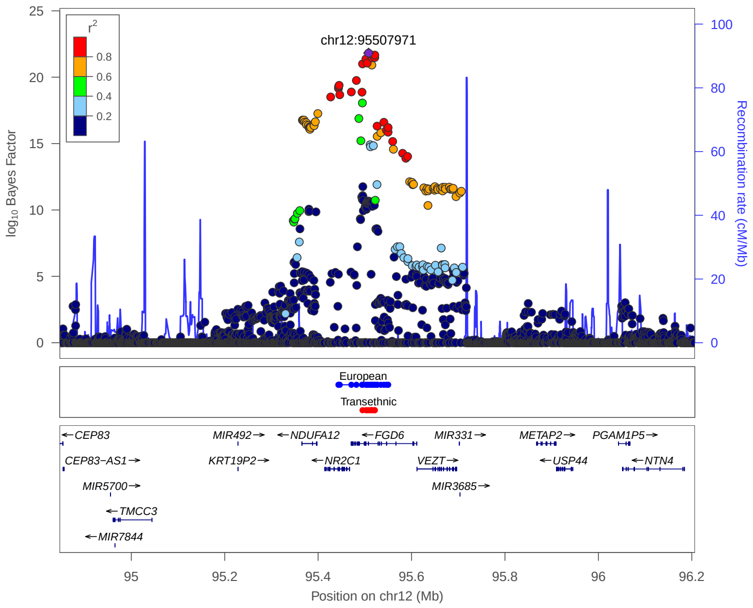


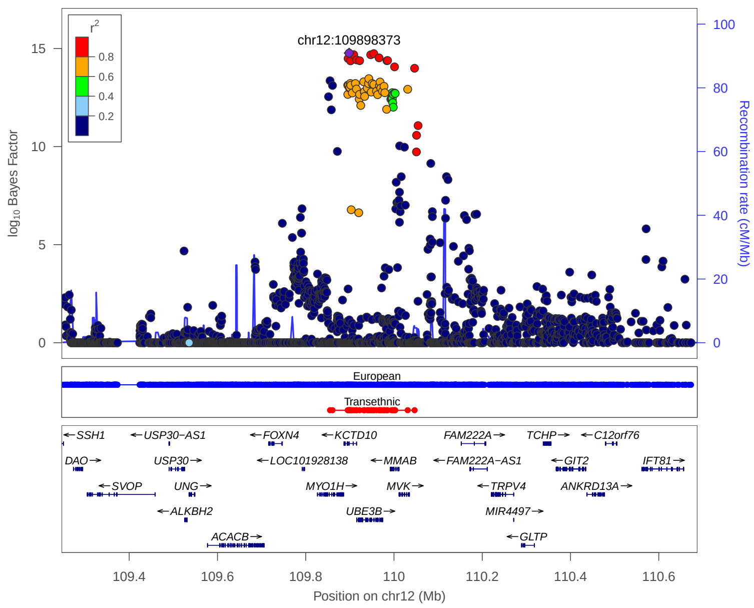


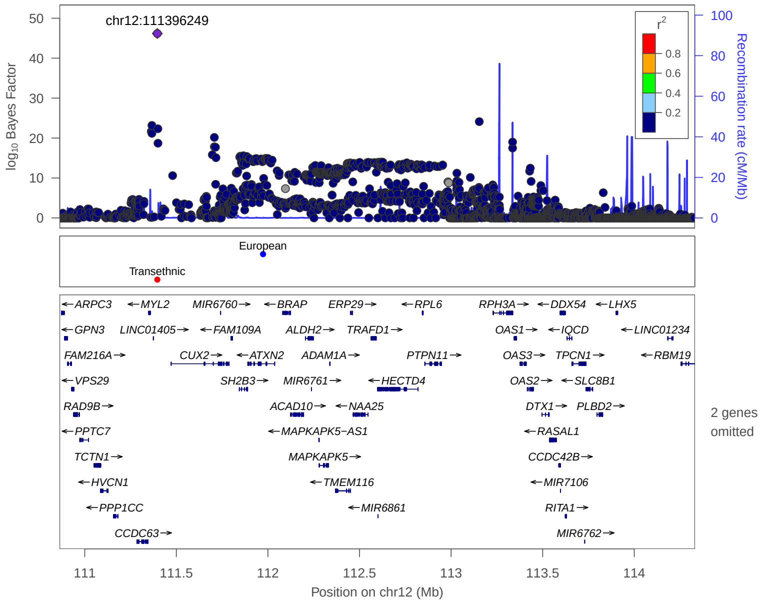


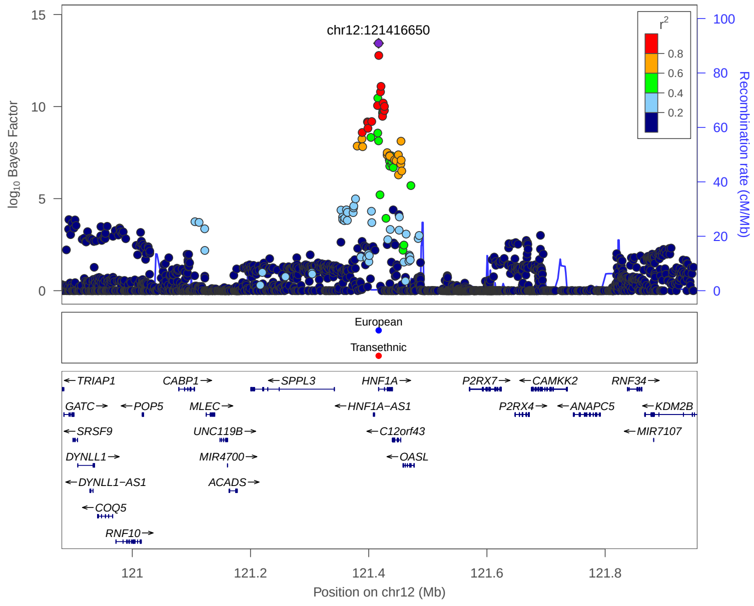


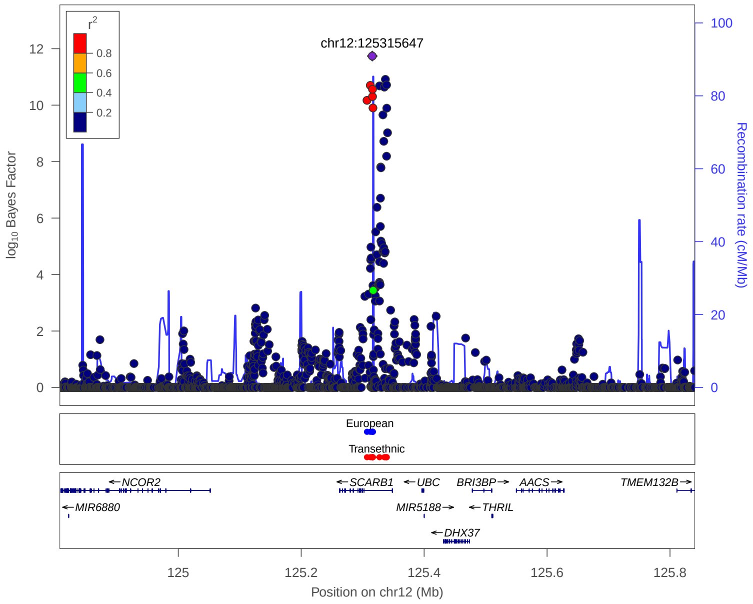


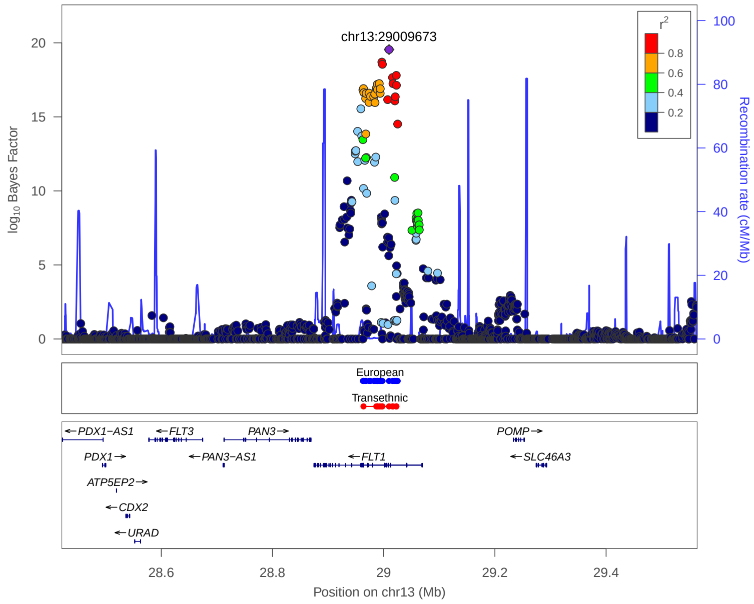


**European:** 99% credible set derived from European meta analysis

**Transethnic:** 99% credible set derived from transethnic meta analysis

### 4. Previously reported 133 loci in Meta analysis

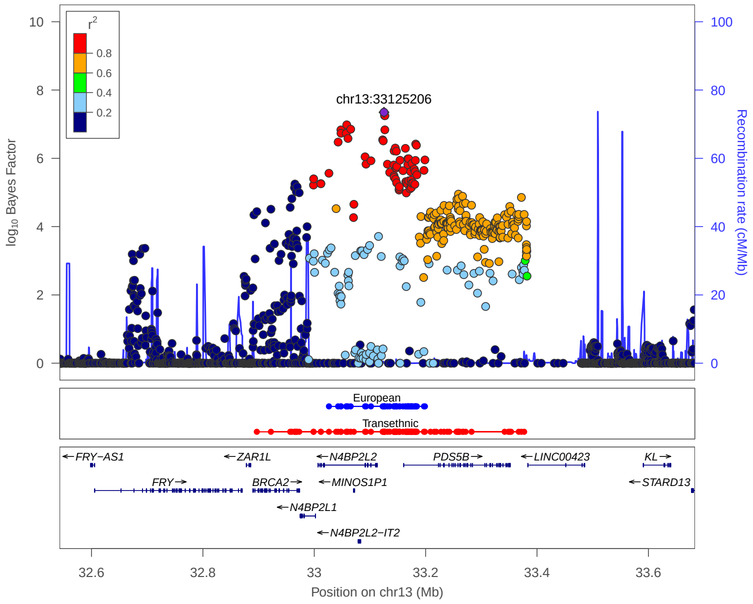


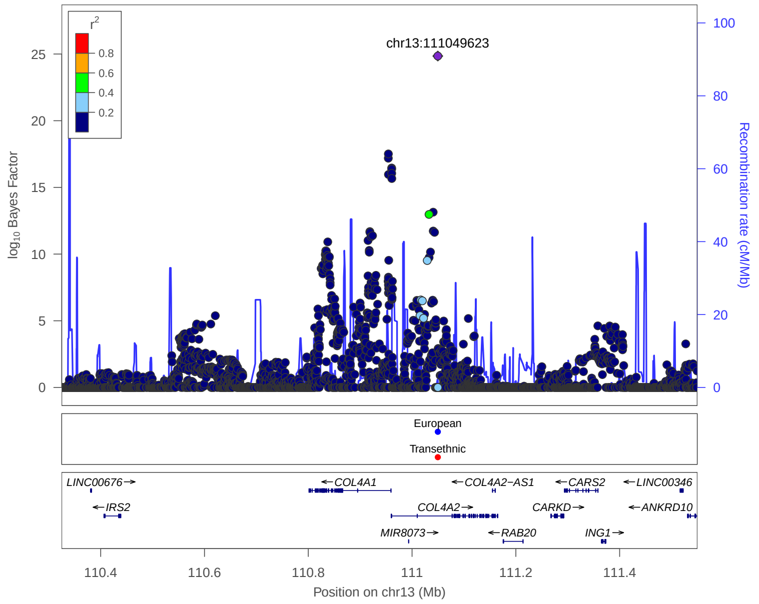


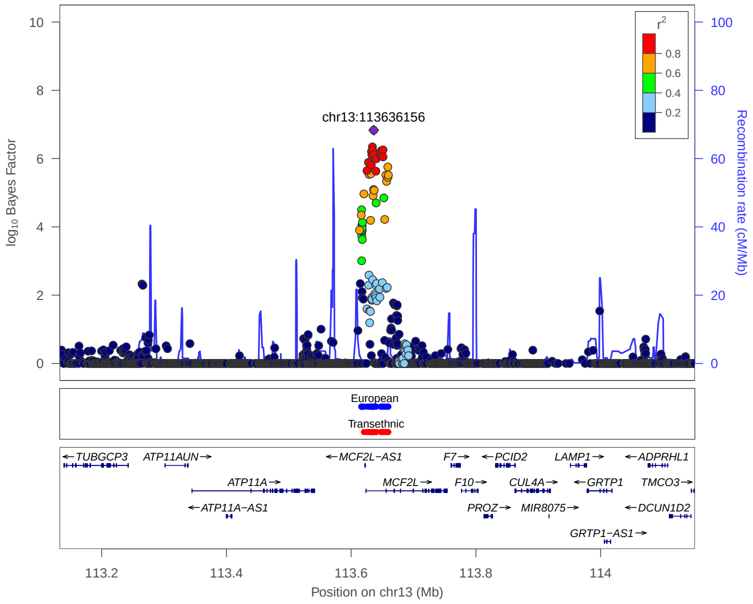


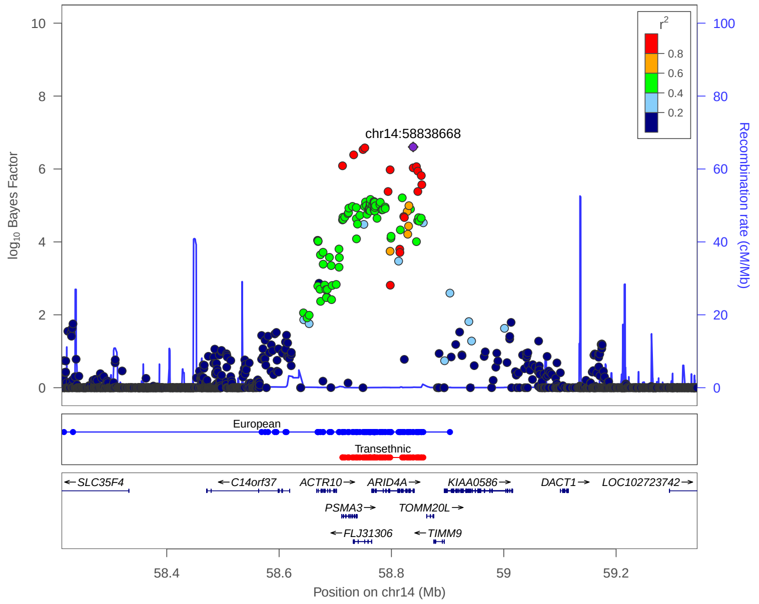


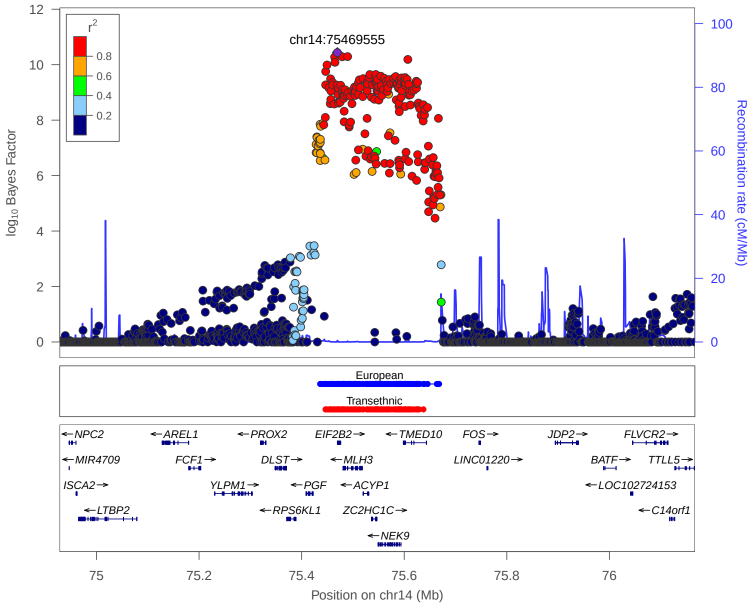


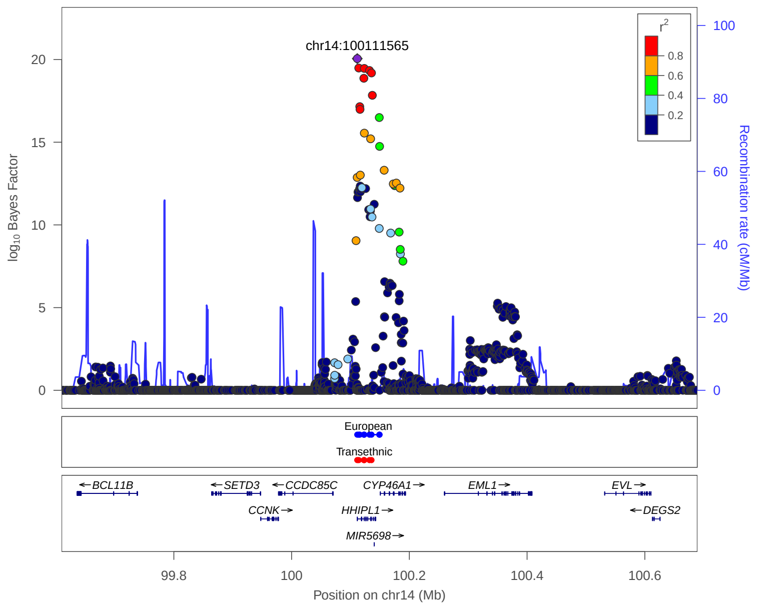


European: 99% credible set derived from European meta analysis

Transethnic: 99% credible set derived from transethnic meta analysis

### 4. Previously reported 133 loci in Meta analysis

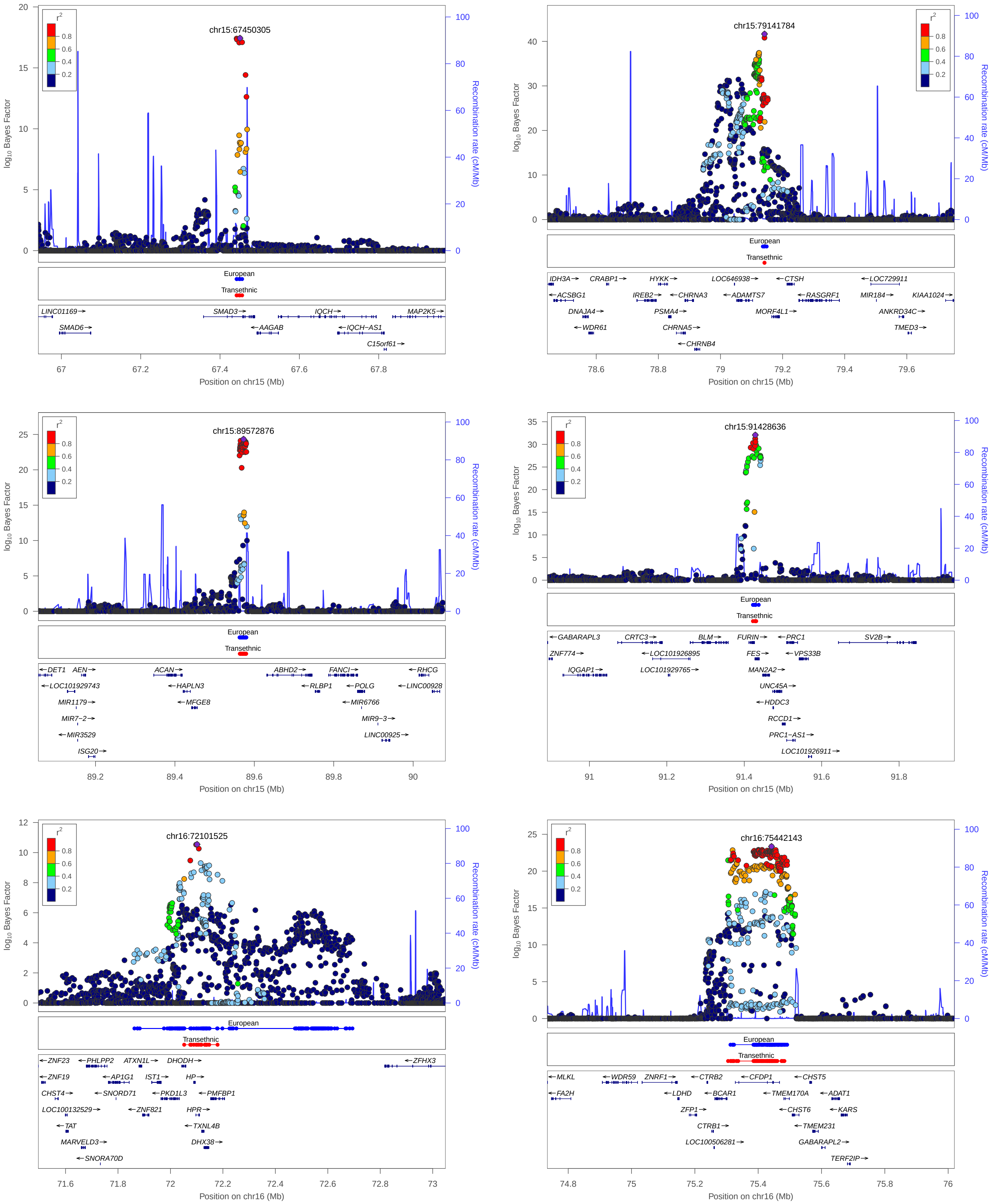


**European:** 99% credible set derived from European meta analysis

**Transethnic:** 99% credible set derived from transethnic meta analysis

### 4. Previously reported 133 loci in Meta analysis

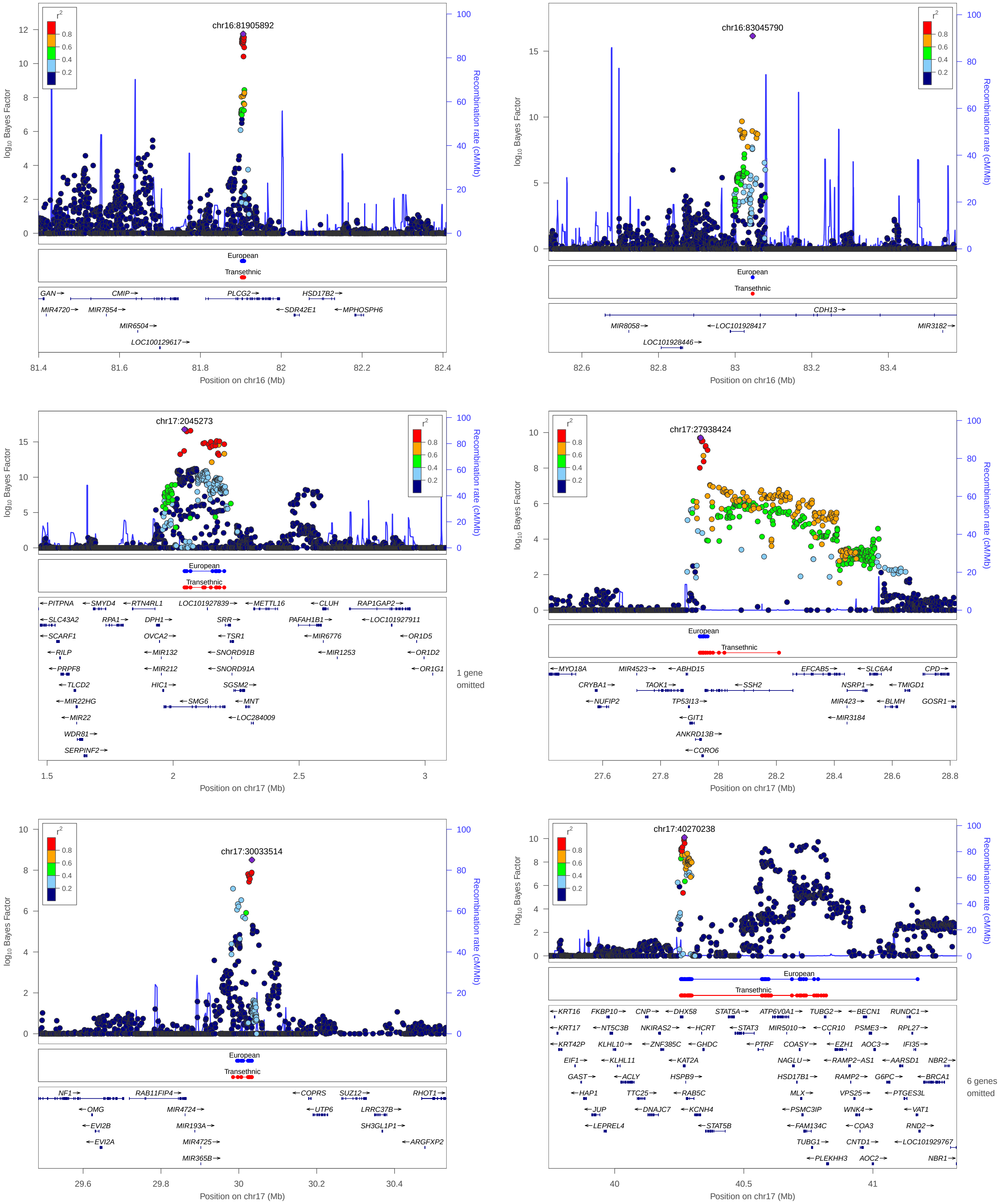


**European:** 99% credible set derived from European meta analysis

**Transethnic:** 99% credible set derived from transethnic meta analysis

### 4. Previously reported 133 loci in Meta analysis

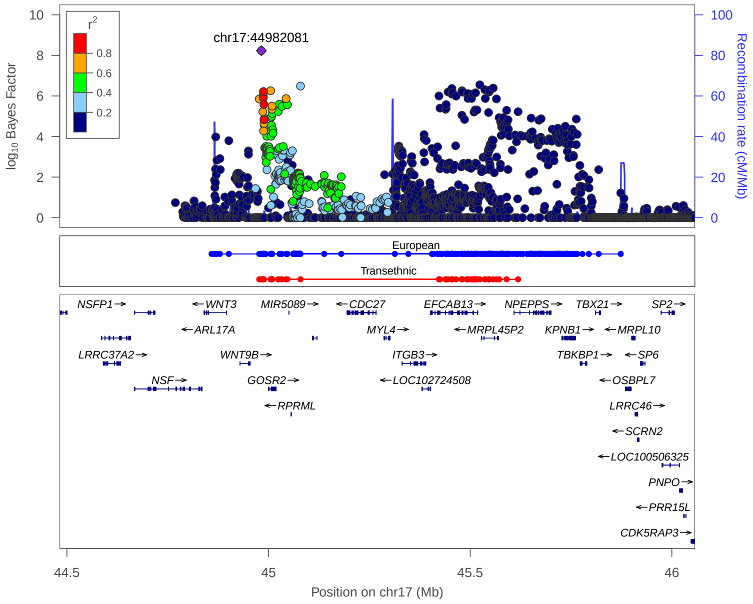


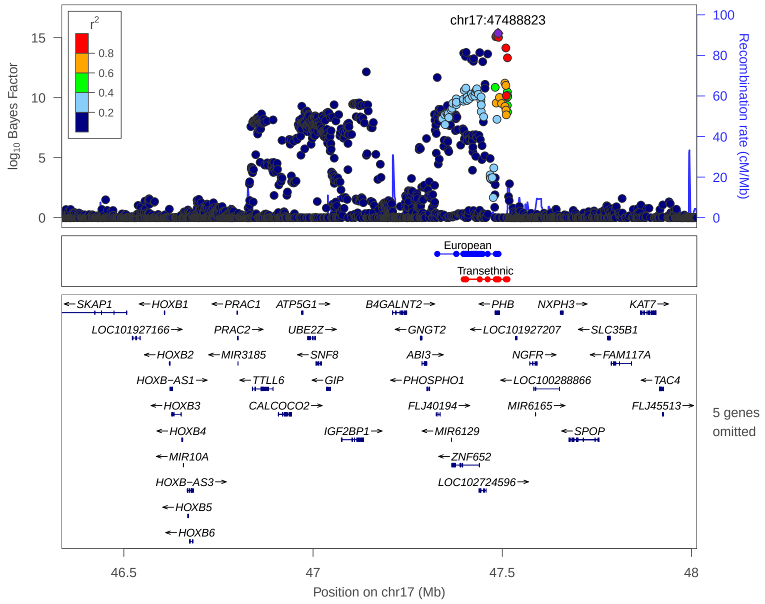


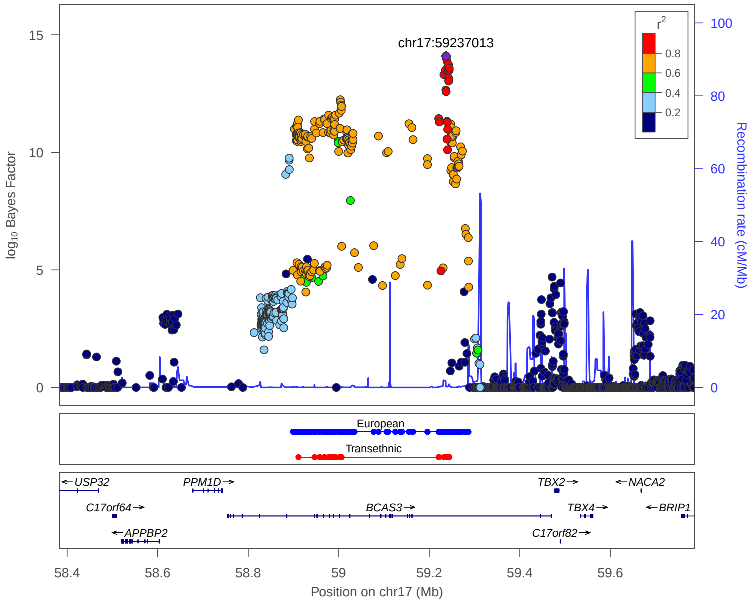


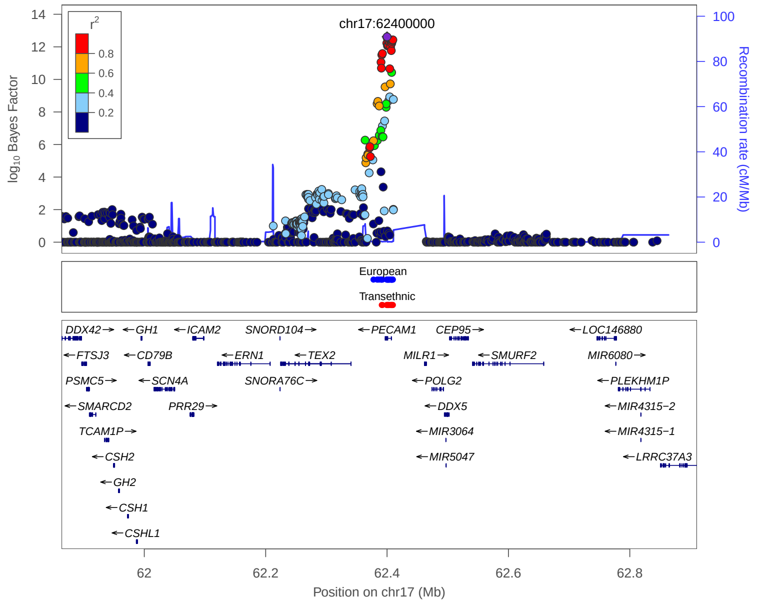


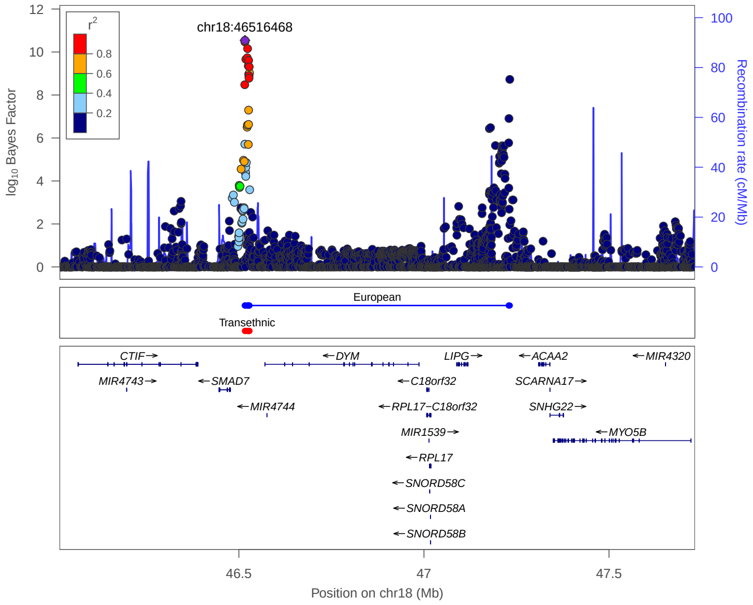


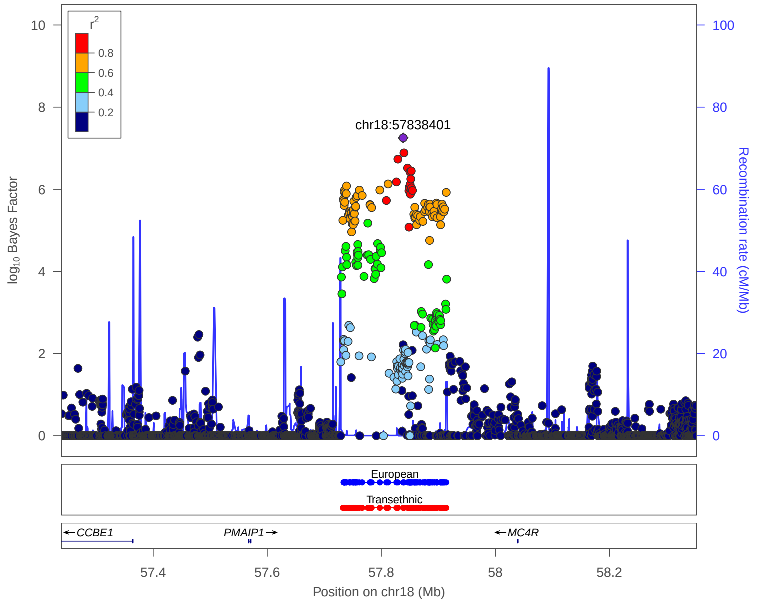


**European:** 99% credible set derived from European meta analysis

**Transethnic:** 99% credible set derived from transethnic meta analysis

### 4. Previously reported 133 loci in Meta analysis

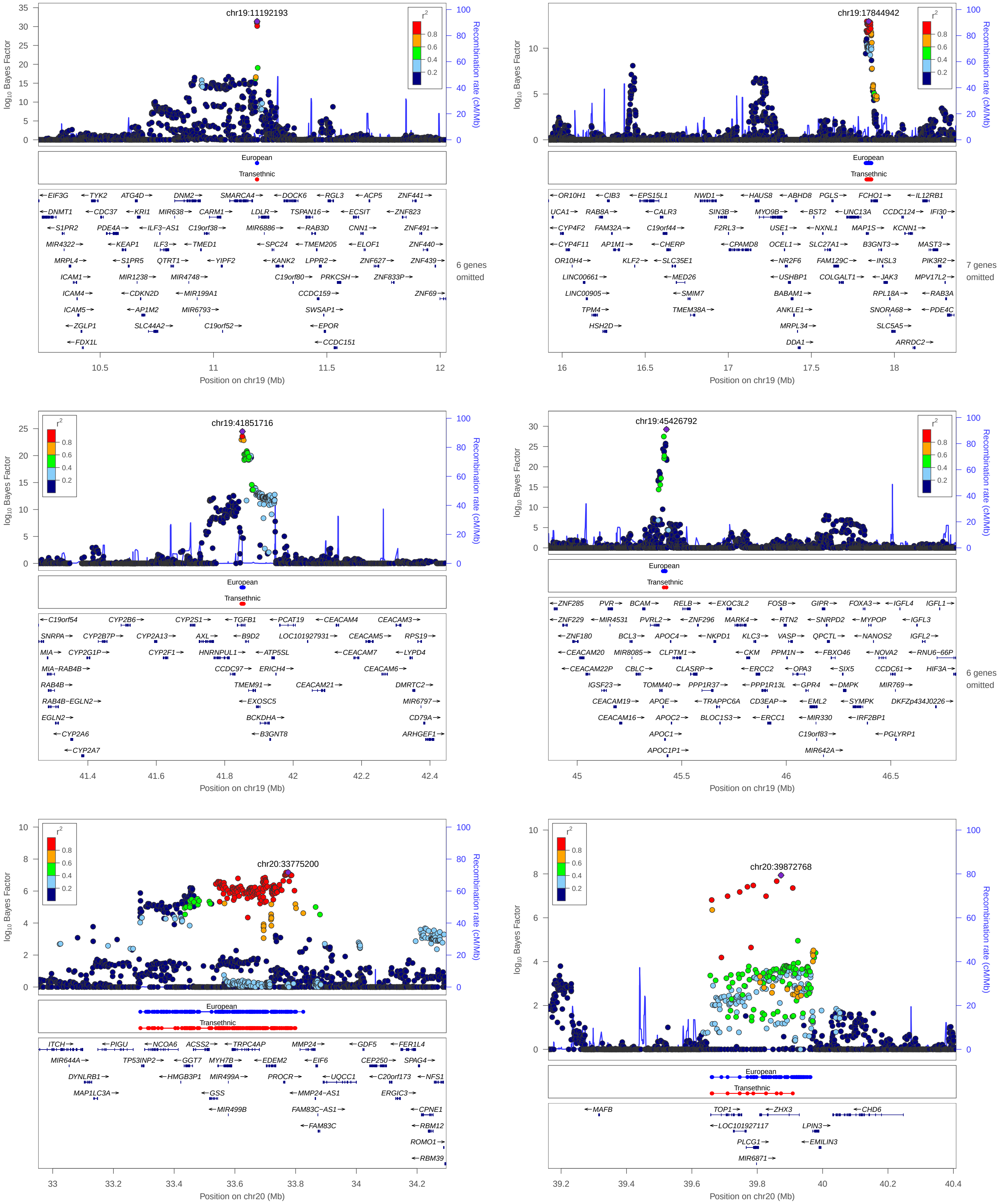


**European:** 99% credible set derived from European meta analysis

**Transethnic:** 99% credible set derived from transethnic meta analysis

### 4. Previously reported 133 loci in Meta analysis

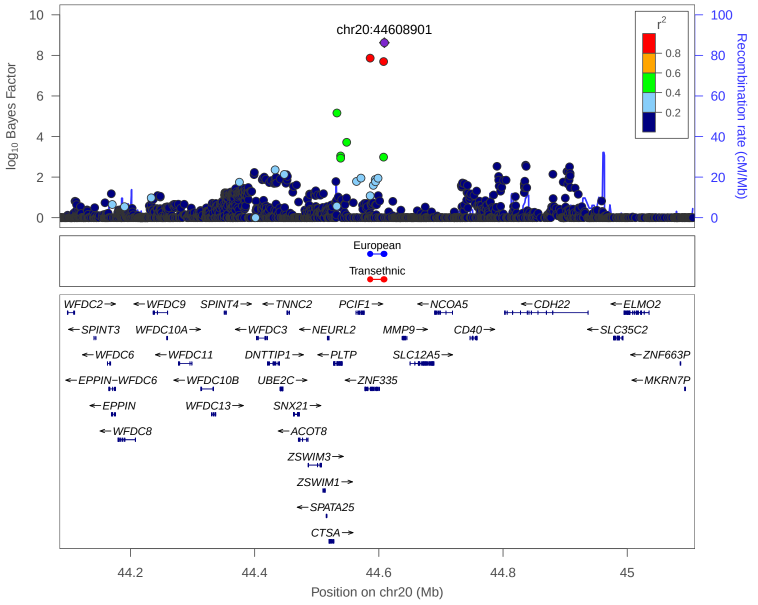


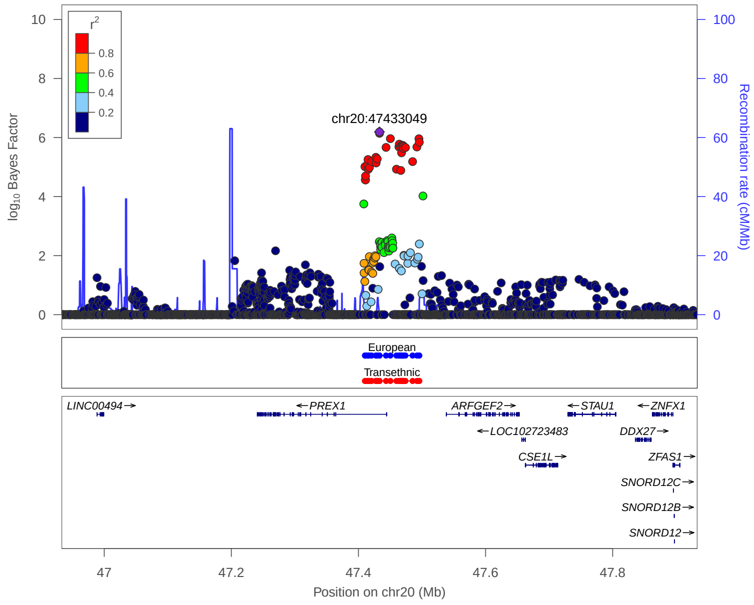


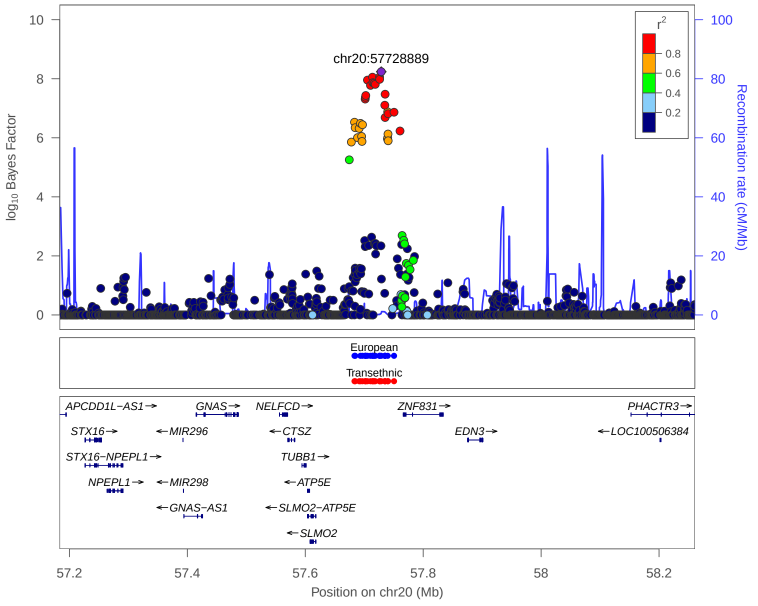


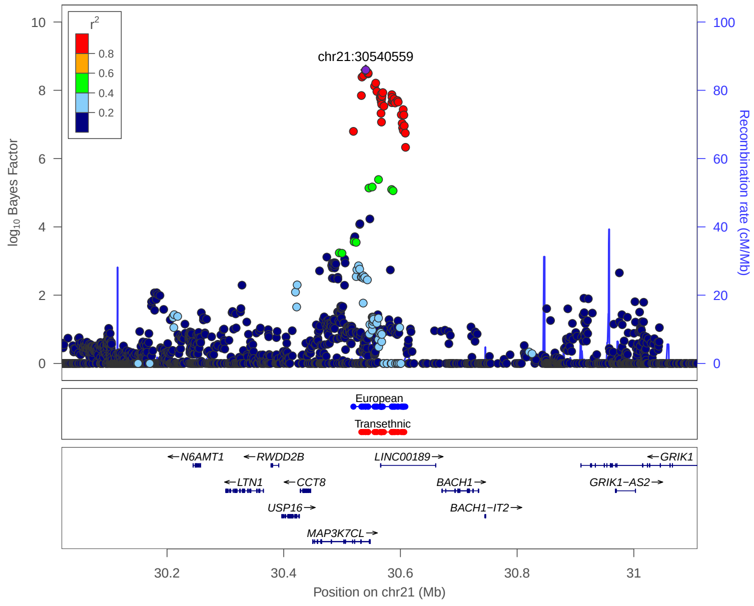


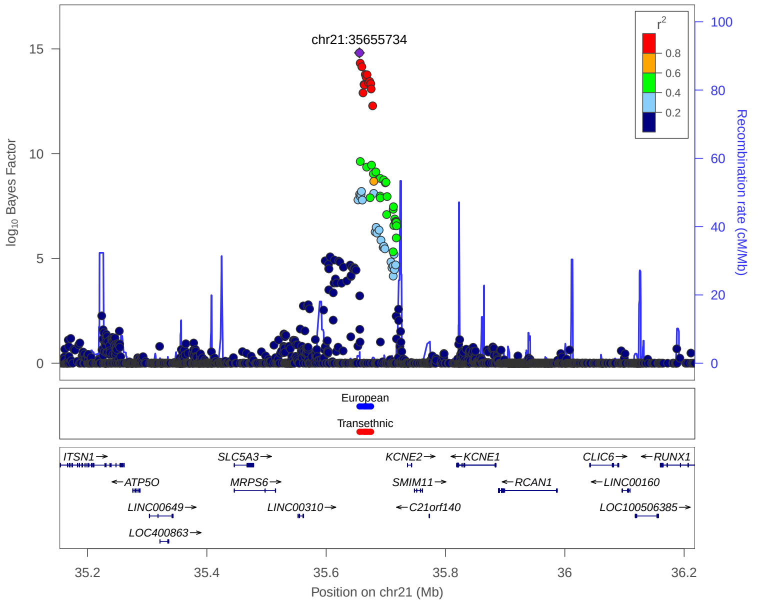


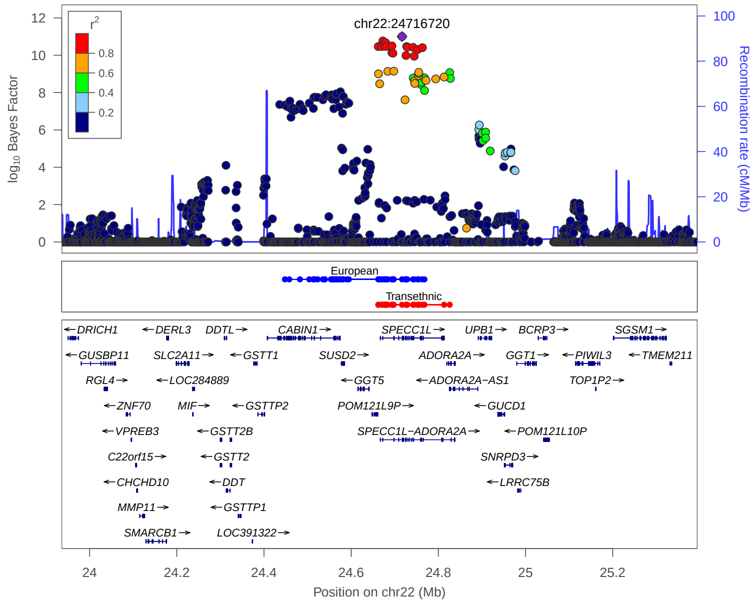


European: 99% credible set derived from European meta analysis

Transethnic: 99% credible set derived from transethnic meta analysis

### 4. Previously reported 133 loci in Meta analysis

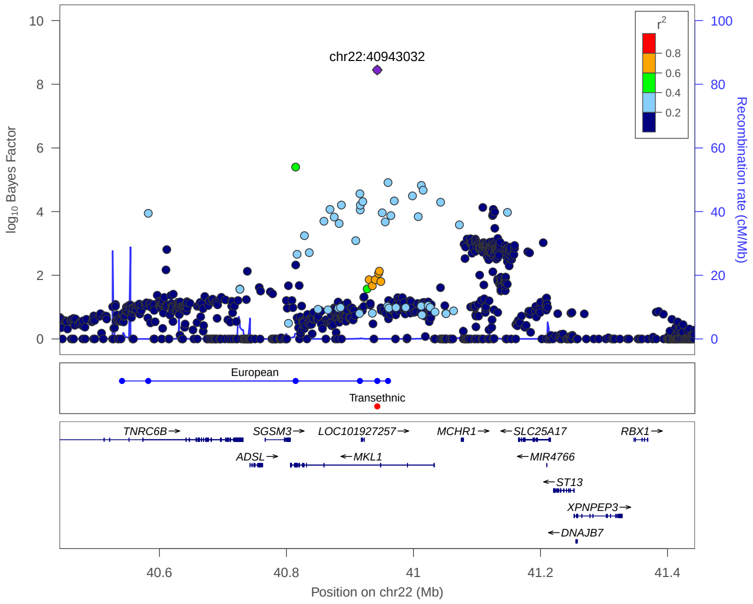


European: 99% credible set derived from European meta analysis

Transethnic: 99% credible set derived from transethnic meta analysis
